# *Trachymyrmex septentrionalis* ants promote fungus garden hygiene using *Trichoderma*-derived metabolite cues

**DOI:** 10.1101/2022.11.12.516288

**Authors:** Kathleen E. Kyle, Sara P. Puckett, Andrés Mauricio Caraballo-Rodríguez, José Rivera-Chávez, Robert M. Samples, Cody E. Earp, Huzefa A. Raja, Cedric J. Pearce, Madeleine Ernst, Justin J.J. van der Hooft, Madison E. Adams, Nicholas H. Oberlies, Pieter C. Dorrestein, Jonathan L. Klassen, Marcy J. Balunas

## Abstract

Fungus-growing ants depend on a fungal mutualist that can fall prey to fungal pathogens. This mutualist is cultivated by these ants in structures called fungus gardens. Ants exhibit weeding behaviors that keep their fungus gardens healthy by physically removing compromised pieces. However, how ants detect diseases of their fungus gardens is unknown. Here, we applied the logic of Koch’s postulates using environmental fungal community gene sequencing, fungal isolation, and laboratory infection experiments to establish *Trichoderma* spp. as previously unrecognized pathogens of *Trachymyrmex septentrionalis* fungus gardens. Our environmental data showed that *Trichoderma* are the most abundant non-cultivar fungi in wild *T. septentrionalis* fungus gardens. We further determined that metabolites produced by *Trichoderma* induce an ant weeding response that mirrors their response to live *Trichoderma*. Combining ant behavioral experiments with bioactivity-guided fractionation and statistical prioritization of metabolites in *Trichoderma* extracts demonstrated that *T. septentrionalis* ants weed in response to peptaibols, a specific class of secondary metabolites known to be produced by *Trichoderma* fungi. Similar assays conducted using purified peptaibols, including the two new peptaibols trichokindins VIII and IX, suggested that weeding is likely induced by peptaibols as a class rather than by a single peptaibol metabolite. In addition to their presence in laboratory experiments, we detected peptaibols in wild fungus gardens. Our combination of environmental data and laboratory infection experiments strongly support that peptaibols act as chemical cues of *Trichoderma* pathogenesis in *T. septentrionalis* fungus gardens.

**Significance Statement:** An extended defense response may exist in any relationship where one partner benefits from defending a mutualistic partner. Such a response is observed in the fungus-growing ant symbiosis, where ants must identify and remove pathogens of their symbiotic fungus gardens. Here we describe the fungal pathogen *Trichoderma* and its associated metabolites, which induce *Trachymyrmex septentrionalis* ant weeding behavior. Ants removed fungus garden pieces inoculated with *Trichoderma* spores or peptaibol-rich *Trichoderma* extracts, and peptaibols as a class cued ant defensive behavior, allowing *T. septentrionalis* to differentiate healthy from diseased fungus gardens. Extended defense responses mediated by chemical cues may be underappreciated mechanisms that structure symbiotic interactions.

## Introduction

The transition to agriculture in humans ∼10,000 years ago is often cited as the key innovation that led to large and complex human civilizations (1, 2). Yet, insects have practiced farming on a much longer timescale. For example, fungus-growing ants have farmed specific “cultivar” fungi (Agaricales: mainly Agaricaceae) for ∼50 million years (3). Fungus-growing ants include over 250 known species, the most complex of which, *Atta* leaf-cutting ants, create colonies that rival large cities in population size (4). Fungus-growing ants obligately rely on their cultivar fungus to digest otherwise inaccessible plant nutrients. In underground fungus gardens, the cultivar fungus breaks down recalcitrant organic material provided by the ants and in return produces specialized structures that the ants consume, making the cultivar fungus a valuable resource for the ants (3, 5).

Valuable crop resources require maintenance and protection. In ant fungus gardens, *Escovopsis* fungi are well known as specialized parasites, especially in the tropics (6, 7). Infection of the fungus garden by *Escovopsis* decreases colony fitness and can lead to colony collapse (6–9). However, *Escovopsis* does not regularly parasitize fungus gardens cultivated by *Trachymyrmex septentrionalis*, the northernmost fungus-growing ant, a finding based on a single culture-based study that detected *Trichoderma* and several other microfungi but not *Escovopsis* in *T. septentrionalis* fungus gardens and that did not test its infectiveness (10). Many other fungi have also been isolated from ant fungus gardens, although the culture-based methods used and the often non-systematic sampling obscures their ecological distribution and symbiotic role (6, 7, 10–15). Of these fungi, only *Trichoderma* and *Syncephalastrum* have been shown to exhibit some degree of pathogenesis towards *Atta* ant fungus gardens [but not towards *T. septentrionalis*; (6, 16, 17)]. More work is needed to fully understand the diversity and ecology of these and other potential fungus garden pathogens and the mechanisms by which ants respond to protect their fungus gardens.

Ants have developed numerous chemical and behavioral mechanisms to avoid infection of their fungus gardens and prevent colony collapse (7). Chemical defenses include the application of antimicrobials from ant metapleural gland and fecal secretions (18–21), and from antibiotic-producing *Pseudonocardia* bacteria that the ants host on their cuticles (22, 23). Ant behavioral defenses include grooming their own bodies and those of their nestmates (24), task partitioning between different members of the colony (25–27), and pre-processing of foraged material before its incorporation into the fungus garden (28, 29). Fungus garden grooming (removal of pathogen spores by moving fungus garden hyphae through ant mouthparts) and especially weeding (removal of infected fungus garden fragments) represent important ant behavioral responses to fungal pathogens that have invaded the fungus garden (17, 27, 30, 31). Although these defensive responses have been described in detail, how fungus-growing ants detect threats to their fungus gardens remains poorly understood (32).

Insects are well-known to communicate using chemical cues. For example, *Mastotermes darwiniensis* termite soldiers respond to the cuticular hydrocarbon *p*-benzoquinone with increased mandible openings, indicating excitement (33). In another example, diverse ant species destroy diseased pupae in response to cuticular hydrocarbons emitted during fungal infection (34). Although the mechanism by which fungus-growing ants detect pathogen infections remains unknown, they do detect and respond to chemical cues that promote other aspects of fungus-garden health. For example, *Acromyrmex lundii* ants use carbon dioxide as a spatial cue to position their fungus gardens at optimum soil depth (35). Based on unfavorable CO_2_ levels, ants will relocate their gardens, a remarkable demonstration of their sensitivity to small molecule cues. Pathogenic fungi also produce metabolites that can affect fungus-growing ant health and behavior such as shearanine D produced by *Escovopsis* fungus garden pathogens, which reduces ant movement, inhibits *Pseudonocardia* growth, and directly causes ant death (36, 37). Thus, chemical communication mechanistically underpins diverse symbiotic interactions in ant fungus gardens.

In this study, we sought to identify the molecular cues that induce hygienic weeding behavior during infections of ant fungus gardens. We established that *Trichoderma* fungi are widespread opportunistic pathogens of *T. septentrionalis* ant fungus gardens and that *T. septentrionalis* ants weed their fungus gardens in response to treatments with live *Trichoderma* spores, *Trichoderma* chemical extracts and fractions, and *Trichoderma*-derived pure compounds. Our results suggest that peptaibol metabolites are produced by *Trichoderma* fungi during fungus garden infection and cue ant weeding behaviors that promote fungus garden hygiene. This study fills the gap between the well-studied hygienic behavioral responses of fungus-growing ants and the hitherto unknown molecular cues that induce them. Such chemical cues are likely widespread in other agricultural systems where they are used by hosts to detect and prevent pathogen infections.

## Results

### *Trichoderma* spp. are common natural inhabitants of *T. septentrionalis* fungus gardens

We used internal transcribed spacer region 2 (ITS2) community amplicon sequencing to investigate microfungal communities in field-sampled and apparently healthy *T. septentrionalis* fungus gardens from across the Eastern USA (*<u>SI Appendix</u>*, Fig. S1, Table S1). Reads classified as *Trichoderma*, a common genus of mycoparasites that also includes some saprophytes and plant mutualists (38), were both the most abundant and prevalent non-cultivar reads in field-sampled *T. septentrionalis* fungus gardens, with a median relative abundance of 1.2% and a maximum of 68.6% in the most extreme case (Fig. 1*A*). Other non-cultivar fungi in these fungus garden samples were only rarely abundant (Fig. 1*A*, *<u>SI Appendix</u>*, Fig. S2), and no reads in this dataset matched the common tropical fungus garden pathogen *Escovopsis*. Only four samples out of 83 were dominated by a non-cultivar fungus; two of these were dominated by *Meyerozyma*, one by *Trichoderma*, and one by an unclassified member of the family *Stephanosporaceae* (Basidiomycota). Although the sampled fungus gardens did not visually appear to be diseased at the time of their collection, our ITS sequencing results suggest that *Trichoderma* spp. may be a low-level but constant threat to *T. septentrionalis* fungus gardens *in situ*.

**Figure 1.**
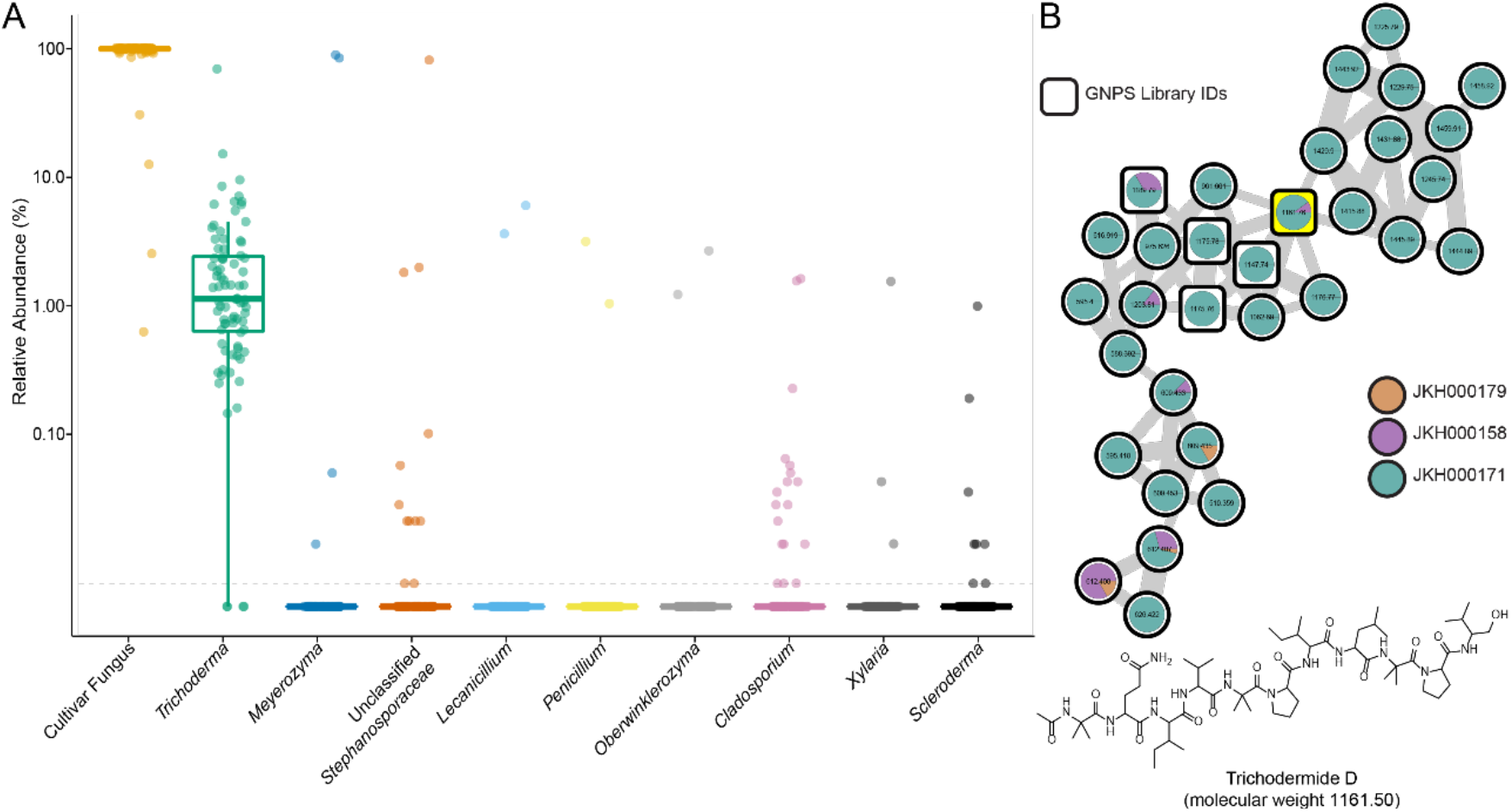
Molecular evidence supporting *Trichoderma* spp. presence in wild *T. septentrionalis* fungus gardens. (A) Relative abundance of ITS2 community amplicon sequencing ASVs shows *Trichoderma* to be the most abundant and most prevalent fungus in wild fungus gardens (n = 83) besides the cultivar fungus. Boxplot center bars show the median relative abundance of ITS2 amplicon reads grouped by genus and the 1st/3rd quartiles are shown by the top/bottom notches respectively. Boxplot whiskers are 1.5 times the interquartile range from the top and bottom notch. Only genera with ASVs that are present at ≥ 1% in at least one sample are shown. The limit of detection (dotted gray line) was determined to be 100*(1/12531). Taxa on the x-axis are ordered from left to right by decreasing mean abundance. (B) Molecular family of peptaibols detected from environmentally-collected fungus garden extracts (*<u>SI Appendix</u>*, Fig. S3), including the node with *m/z* 1161.76 (highlighted in yellow) whose molecular weight and fragmentation pattern were consistent with trichodermide D (*<u>SI Appendix</u>*, Fig. S4), a peptaibol previously isolated from *Trichoderma* spp. (40). This molecular family contains related peptaibol metabolites from three fungus garden colonies, all from North Carolina. Square nodes represent spectral matches in the GNPS public libraries.

In a parallel analysis, we also generated environmental metabolomes from 53 field-sampled *T. septentrionalis* fungus gardens, 18 of which were the same as those sampled for our ITS dataset. Using untargeted LC-MS/MS and Global Natural Products Social (GNPS) molecular networking (39), we identified chemical evidence for the presence of *Trichoderma* spp. in *T. septentrionalis* fungus gardens. After searching our network of specialized metabolites from these freshly collected fungus gardens (*<u>SI Appendix</u>*, Fig. S3) for metabolites produced by potential pathogens, we identified one cluster that contained a feature whose molecular weight and fragmentation pattern were consistent with the peptaibol trichodermide D (Fig. 1*B*, *<u>SI Appendix</u>*, Fig. S4) along with a suite of related peptaibols, most of which were originally isolated from *Trichoderma virens* CMB-TN16 (40). A related series of peptaibol features were detected in three fungus gardens collected from North Carolina, as shown by the nodes in the network that were closely related to known peptaibol features (Fig. 1*B*). Given that peptaibols are characteristic of mycoparasitic members of the Hypocreales and especially prevalent in *Trichoderma* (41, 42), our detection of peptaibols from these samples further supports that *Trichoderma* is present and metabolically active in field-sampled *T. septentrionalis* fungus gardens.

### *Trichoderma* spp. can infect *T. septentrionalis* fungus gardens

To experimentally test our hypothesis that some *Trichoderma* spp. can infect *T. septentrionalis* fungus gardens, we performed infections using lab-acclimated *T. septentrionalis* colonies. Three different *Trichoderma* strains (JKS001879, JKS001884, and JKS001921; most closely related to *T. koningiopsis*, *T. virens*, and *T. simmonsii*, respectively; *<u>SI Appendix</u>,* Table S2, Fig. S5) were used for these infection experiments. These strains were all isolated from *T. septentrionalis* colonies that had naturally become diseased after the removal of ants in our laboratory. By four days post-treatment, all *Trichoderma*-treated fungus gardens without ants became visibly infected with these *Trichoderma* (confirmed using full length ITS sequencing). However, those with ants and the mock-inoculated control fungus gardens (both with and without ants) remained uninfected (Fig. 2*A*, *<u>SI Appendix</u>*, Fig. S6). Ants weeded their fungus gardens in response to the application of all three *Trichoderma* strains (*<u>SI Appendix</u>*, Fig. S6), which presumably helped to maintain fungus garden health. In addition, peptaibols were produced by all three *Trichoderma* strains, as determined using untargeted metabolomics of the infected fungus gardens (*<u>SI Appendix</u>*, Fig. S7). These experiments demonstrate the ability of three different *Trichoderma* spp. to infect *T. septentrionalis* fungus gardens and further link *Trichoderma* infection with the presence of peptaibols.

**Figure 2.**
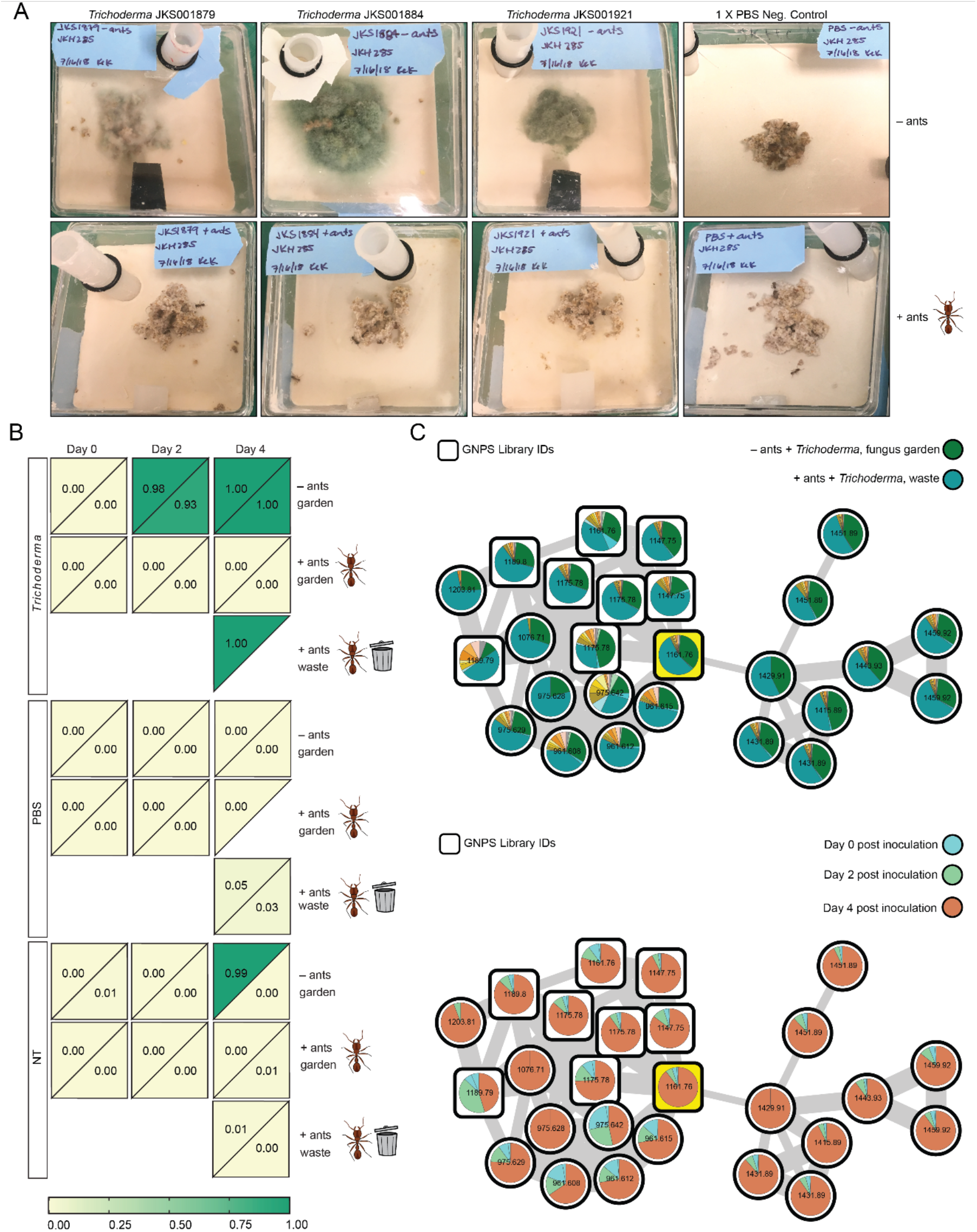
*Trichoderma* spp. can infect *T. septentrionalis* fungus gardens. (A) *Trichoderma* spp. infected *T. septentrionalis* laboratory fungus gardens without ants but not fungus gardens with ants at 4 days post treatment. The uninoculated PBS control treatment both with and without ants showed no visible signs of disease. Attached sideboxes where ants can deposit weeded pieces of fungus garden as waste are shown in *<u>SI Appendix</u>*, Fig. S6 for the treatments with ants. (B) ITS2 community amplicon sequencing detects the growth of *Trichoderma* sp. JKS001884 over time following infection. Duplicate subcolonies of JKH000292 were inoculated with *Trichoderma* sp. JKS001884 spores in PBS, sterile PBS, or no treatment and sampled for DNA at days 0, 2, and 4 post inoculation. The relative abundance of *Trichoderma* ITS reads are shown as a heatmap labelled with each value (sample replicates are triangles forming half a square per sample; only half-square triangles are shown when replicates failed to sequence). *Trichoderma* relative abundance was high in the *Trichoderma*-treated fungus gardens without ants, in the waste of *Trichoderma*-treatments with ants, and in one NT -ants replicate. *Trichoderma* was detected at lower relative abundance in the waste samples of the mock and non-inoculated +ants treatments and not at all in remaining samples. See *<u>SI Appendix</u>*, Fig. S7 for experimental images. (C) LC-MS/MS molecular networks of peptaibol-like features detected in the time-course JKS001884 infection. The top network has node pie charts colored according to feature abundance in each experimental treatment (control treatment colors are not shown in the legend for simplicity; see *<u>SI Appendix</u>*, Fig. S9 for the full color legend that includes all treatments and the complete molecular network). The bottom network shows node pie charts colored according to feature abundance from each time point post-inoculation (see *<u>SI Appendix</u>*, Fig. S11 for the complete molecular network). Peptaibol-like features were most abundant in *Trichoderma*-treated fungus gardens lacking ants and in the waste of *Trichoderma* treatments that included ants, and these features increased in abundance over infection time. The highlighted node is consistent with trichodermide D (*<u>SI Appendix</u>*, Fig. S4) and square nodes represent spectral matches in the GNPS public libraries.

We conducted an additional time-course experiment to further characterize *Trichoderma* infections of ant fungus gardens. By day 4 post-inoculation, *Trichoderma* sp. JKS001884 was visible in *Trichoderma*-treated fungus gardens in subcolonies that lacked ants, and in the waste piles created by the ants in the subcolonies that included them. All uninoculated negative controls lacked visible signs of disease throughout the time-course (*<u>SI Appendix</u>*, Fig. S8). These visual observations were verified using community ITS2 gene sequencing (Fig. 2*B*), in which *Trichoderma* ITS reads were detected 2 and 4 days post-inoculation in the *Trichoderma*-treated fungus gardens in subcolonies that lacked ants, and in the waste generated by ants following *Trichoderma* treatments (waste was sampled only at day 4 post-treatment). All uninoculated fungus gardens (with and without ants) and *Trichoderma*-treated fungus gardens in subcolonies that included ants lacked *Trichoderma* ITS reads, except for one untreated fungus garden lacking ants that contained mostly *Trichoderma* reads by day 4 post treatment, likely due to a bloom of *Trichoderma* that was naturally present in the subcolony and whose suppression was released by ant removal. Day 4 waste samples from uninoculated fungus gardens tended by ants also contained low abundances of non-cultivar fungi (including *Trichoderma*) in control treatments. These results indicate that *Trichoderma* can persist for several months in ant fungus gardens even when colonies were fed only sterile corn meal throughout that time. Molecular networking confirmed that peptaibol abundance was associated with the presence and abundance of *Trichoderma* during these infections (Fig. 2*C*, *<u>SI Appendix</u>,* Fig. S9-S10), mainly in the waste produced by ants following *Trichoderma* treatment and in *Trichoderma*-treated fungus gardens lacking ants. Peptaibol abundance also increased over time following infection in these samples (Fig. 2*C*, *<u>SI Appendix</u>,* Fig. S11). These results confirmed the ability of *Trichoderma* spp. to infect *T. septentrionalis* fungus gardens and linked the presence of peptaibols with that of *Trichoderma*.

### *Trichoderma* metabolites promote ant weeding behavior

To identify the underlying mechanisms by which *Trichoderma* inoculation induced waste production by the ants, we exposed *T. septentrionalis* fungus gardens to crude extracts of *Trichoderma* sp. JKS001884. Colonies were tested multiple times, including both intra- and inter-colony replicates (*<u>SI Appendix</u>*, Table S4). In all tests, ant waste production was greater in extract-treated fungus gardens (mean weeded mass of 395 mg ± 40.9) compared to the negative controls treated only with DMSO (91.7 mg ± 19.4) or left untreated (41.9 mg ± 14.2; Fig. 3*A*, *<u>SI Appendix</u>*, Fig. S12). These results suggest that ant waste production was induced by metabolites from the *Trichoderma* extracts and that ant responses were not solely due to the physical presence of *Trichoderma* cells.

**Figure 3.**
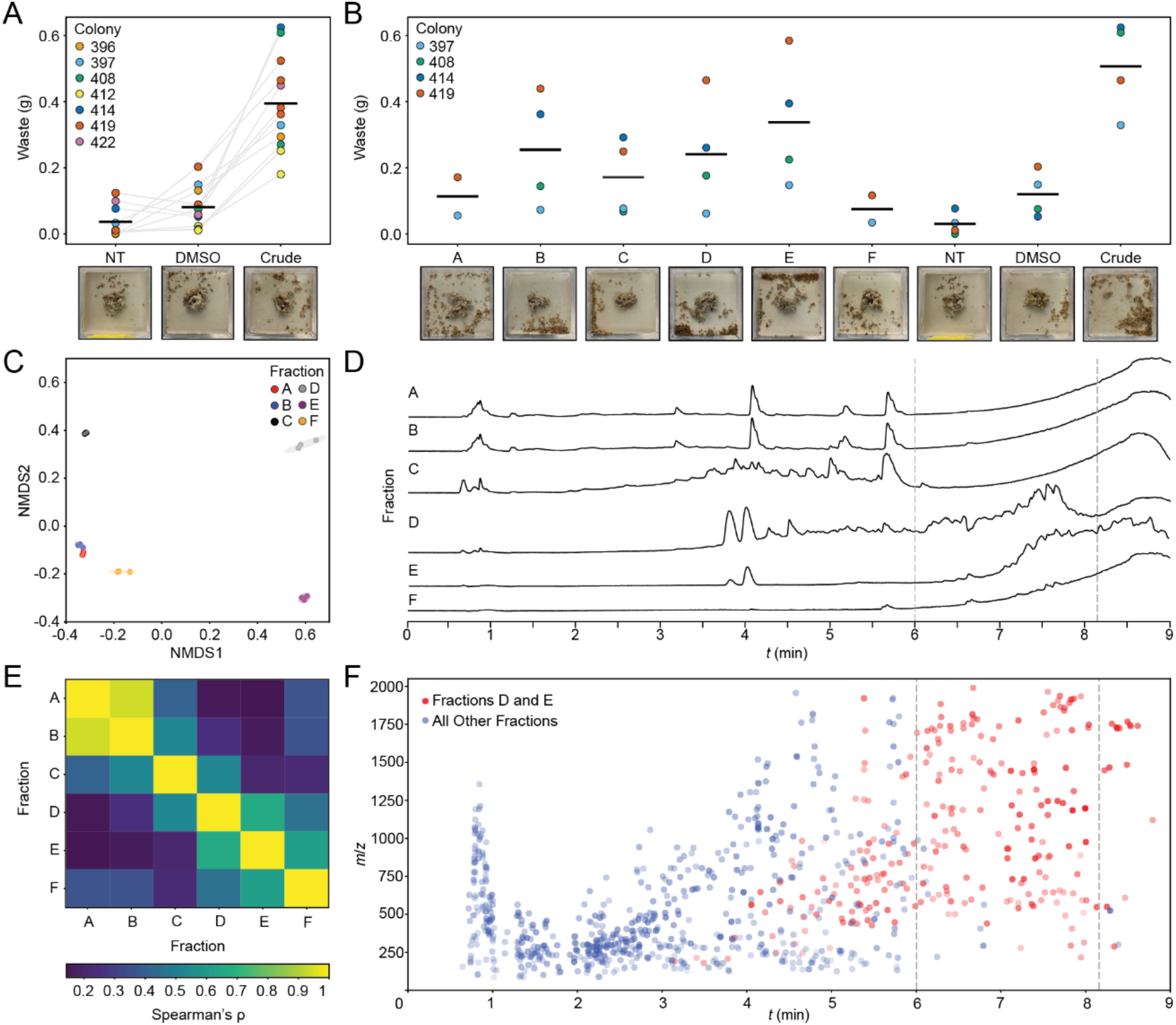
*Trichoderma* metabolites promote ant weeding behavior. (A) *T. septentrionalis* ants deconstructed (“weeded”) their fungus gardens when exposed to crude *Trichoderma* extract (DMSO = solvent control, NT = no treatment). Colors indicate unique ant colonies, and values from biologically independent trials are connected. Representative images are shown (see Data Availability for full dataset). (B) Exposure to fractions D and E of *Trichoderma* extracts resulted in substantive weeding behavior. Fraction B also resulted in weeding but was deprioritized based on metabolomic similarity with inactive fraction A (seen in C and D). Using LC-MS/MS and statistical analyses, fractions D and E grouped together along NMDS1 (C), distinct from other fractions. (D) Total ion chromatograms (TICs) indicated considerable overlap in peaks from fractions D and E, especially those with elution times/solvent systems consistent with peptaibol metabolites (dashed lines). Fractions D and E also exhibited high chemical similarity as seen via Spearman correlations (E). (F) Many of the features that co-occur in the bioactive fractions D and E (shown in red) have molecular weights above 1000 *m/z*, also consistent with peptaibol metabolites (blue features indicate features found in all other fractions).

To identify the metabolites responsible for this bioactivity, we evaluated semi-purified fractions of our *Trichoderma* extract for their ability to induce ant behavioral responses. Fractions B, D, and E induced the greatest amount of ant waste production (255 mg ± 87.0, 241 mg ± 85.1, and 338 mg ± 97.1, respectively; Fig. 3*B*, *<u>SI Appendix</u>*, Fig. S13), although further analyses determined that fraction B was chemically dissimilar to fractions D and E (Fig. 3*C-F*, *<u>SI Appendix</u>*, S15-17) and was instead highly similar to fraction A (Fig. 3*C-F*), one of the least bioactive fractions (Fig. 3*B*). Given that such early fractions often contain pan-assay interference compounds (PAINs)-type compounds (43, 44), we prioritized fractions D and E for further analysis.

Comparative metabolomics was used to prioritize and identify metabolites that were highly abundant in fractions D and E. These fractions grouped together along NMDS axis 1, distinct from all other fractions (Fig. 3*C*), demonstrating their high chemical similarity, as also determined using Spearman’s correlation (Fig. 3*E*). Comparisons of total ion chromatograms (TICs; Fig. 3*D*, *<u>SI Appendix</u>*, Fig. S18) indicated considerable overlap of peaks in fractions D and E especially between 7-8 minutes, retention times at which numerous peptaibols elute. In addition, a large number of features that co-occurred in fractions D and E had molecular weights above 1,000 Da, consistent with peptaibol metabolites (Fig. 3*F*). This motivated further chemical analysis to identify features that may underpin the ant behaviors induced by fractions D and E.

### Metabolomic and statistical analyses reveal peptaibols from *Trichoderma* induce ant waste production

Based on our prioritization of fractions D and E (Fig. 3), we generated a heat map to identify features shared between fractions D and E and dereplicated these features using NP Atlas (45) (Fig. 4*A*, full heatmap in *<u>SI Appendix</u>*, Fig. S16). Fractions D and E exclusively shared 118 features (*<u>SI Appendix</u>*, Fig. S15), although an additional suite of features was highly enriched in fractions D and E but present in lower abundances in other fractions. Based on their molecular weights, retention times, and fragmentation patterns, many of these shared features were consistent with the structure of peptaibols. Three peptaibol-like features were exceptionally abundant in fractions D and E (Fig. 4*C*, *<u>SI Appendix</u>*, Table S8): [M + Na]^+^ peaks *m/z* 1197.7557 and 1183.7406, and [M + H]^+^ peak *m/z* 1452.8756. Together, the ion abundances of these three features represented a combined 35.5% and 75.5% of total metabolite abundance in fractions D and E, respectively. Further exploration of these features confirmed that all three represent peptaibols, two of which have masses consistent with the peptaibols trichodermides B/C and D/E (*<u>SI Appendix</u>*, Fig. S19), although several other peptaibols have similar masses.

**Figure 4.**
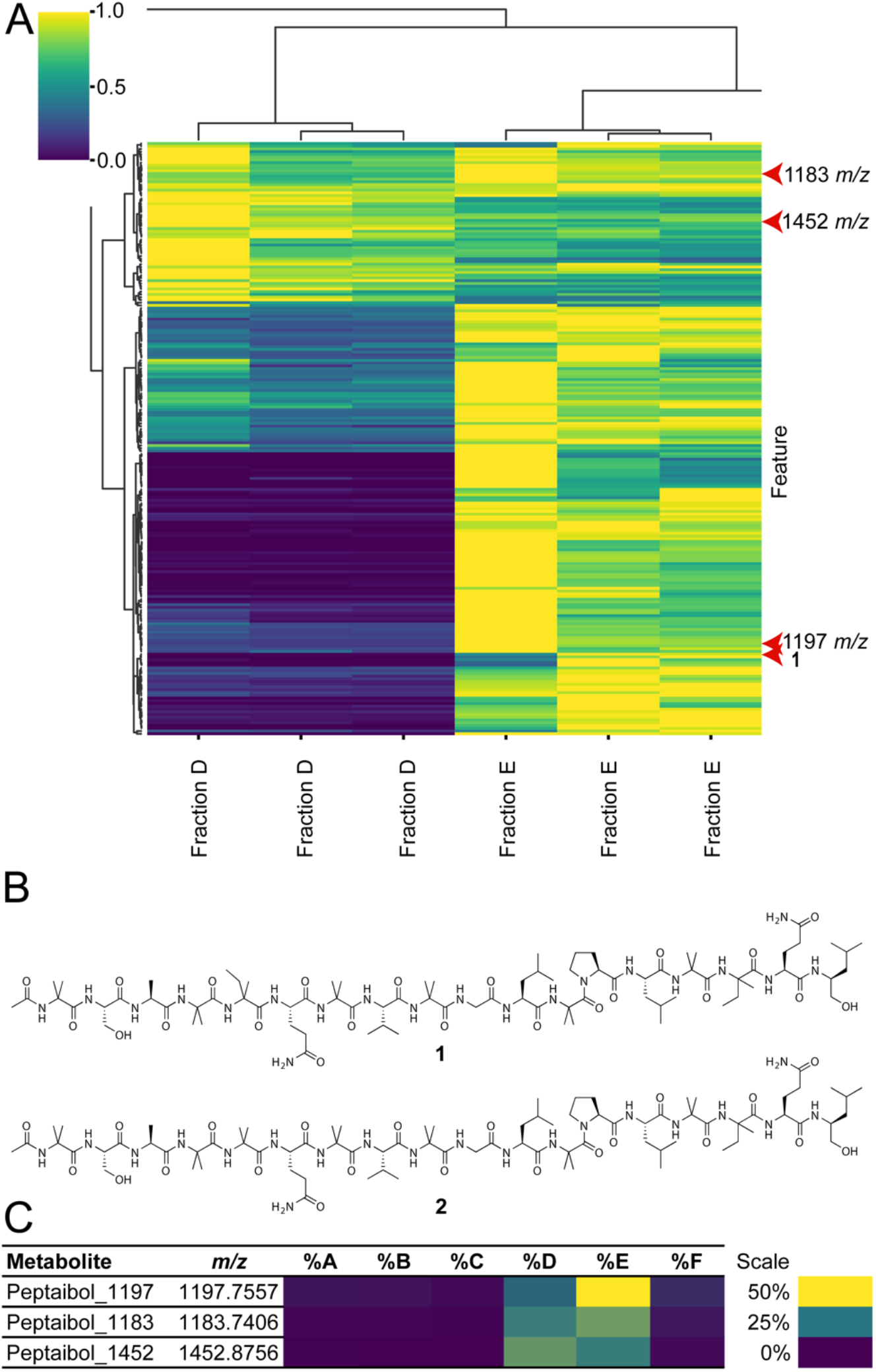
Peptaibols from *Trichoderma* induce ant waste production. (A) Fractions D and E from the *Trichoderma* extract exhibit considerable similarity in the peptaibol-rich portion of a heatmap of feature abundance (full heatmap in *<u>SI Appendix</u>*, Fig. S16). Three of these features include the peptaibol-like features with [M + Na]^+^ peaks *m/z* 1197.7557 and 1183.7406, and [M + H]^+^ peak *m/z* 1452.8756. (B) Two newly identified peptaibol metabolites, compounds **1** and **2**, were also found to induce ant waste production (*<u>SI Appendix</u>*, Fig. S14), and were most abundant in fraction E (*<u>SI Appendix</u>*, Table S8). (C) Three of the most abundant metabolites in fractions D and E were determined to be peptaibols, representing a combined 35.5% and 75.5% of fractions D and E, respectively. Two of these had masses consistent with trichodermides B/C and D/E, although further confirmation is needed for conclusive identification.

These results prompted further ant weeding assays using a small library of purified peptaibols to determine if specific, individual peptaibols induce ant behavior and if this behavior is a result of collective and/or non-specific peptaibol metabolites. We tested six peptaibols isolated from *Trichoderma arundinaceum* and one purchased peptaibol to evaluate their ability to induce ant waste production. Although these purified peptaibols varied in their bioactivity, all induced ant waste production, some with higher levels of bioactivity than the crude extract (*<u>SI Appendix</u>*, Fig. S14). Two of these peptaibol metabolites, **1** and **2** (Fig. 4*B*), are new compounds given the trivial names trichokindins VIII and IX, respectively. Mass spectrometric analysis confirmed that both metabolites were present in our crude *Trichoderma* extract and in the bioactive fractions D and E (*<u>SI Appendix</u>*, Figs. S20-21). Full characterization of **1** and **2** (*<u>SI Appendix</u>*, Figs. S22-40) indicated that these new compounds have a classical peptaibol structure, being composed entirely of amino acids, including characteristic α-aminoisobutyric acid (Aib) moieties. Because each isolated peptaibol induced ant weeding, this behavior is likely characteristic of the peptaibol class, in general, and may not be specific to individual peptaibols.

Our results strongly suggest that ant waste production is induced by *Trichoderma*-derived peptaibol metabolites, however, we cannot discount that other fungal secondary metabolites were present in our *Trichoderma* extracts. Using comparative metabolomics, we identified a feature suggestive of the fungal metabolite roselipin 1A (46), although this feature had similar abundances across fractions D, E, and F. Because fraction F did not induce substantial ant waste production, this feature did not likely cause the observed ant weeding behavior. Features with masses matching two other common fungal metabolites, gliovirin (47) and heptelidic acid (48, 49), exhibited similar patterns of abundance in both inactive and bioactive fractions, and thus were also unlikely to induce the observed ant weeding behavior. Thus, the unique correlation between peptaibol-enriched fractions and increased ant waste production underscores the relationship between *Trichoderma*-derived peptaibols and ant waste production.

## Discussion

Our data show that *Trichoderma* spp. are common in wild *T. septentrionalis* fungus gardens sampled from across a broad geographic range (Fig. 1*A*), which suggests that at least some *Trichoderma* spp. may naturally cause fungus garden disease. Supporting this hypothesis, our isolation of *Trichoderma* spp. from diseased gardens, experimental infection of healthy gardens, and detection of *Trichoderma* during experimental infections using ITS2 amplicon sequencing all provide experimental evidence that *Trichoderma* spp. can be opportunistic pathogens of *T. septentrionalis* fungus gardens (Fig. 2*A-B*) and fully satisfy the experimental aspects of Koch’s postulates for disease causality, which include the isolation of a pathogen from a diseased host, experimental infection of a naive host using that isolate, and reisolation of that disease-causing isolate (50–52). The ecological aspects of Koch’s postulates (detection of the pathogen in diseased but not healthy hosts) are partially fulfilled in this study by our detection of *Trichoderma* in environmental fungus gardens, even though these were not visually diseased at the time of collection, a difficult to detect event due to acute disease leading to rapid colony collapse and our detecting colonies to collect based on the presence of active ants, likely indicative of colony health. We also note that fungus garden diseases can be present but not visually apparent (e.g., on day 2 of our infection experiments; Fig. 2), making it challenging to definitively link *Trichoderma* presence to disease in the field, and that a focus on pathogen presence/absence without consideration of pathogen load, microbiome composition, or environmental conditions is a noted weakness of Koch’s postulates (50, 53, 54). Although the species identity, ecological source, and relationship to disease of the low levels of *Trichoderma* present in *T. septentrionalis* fungus gardens remains unclear, our results indicate the consistent presence of these potential disease-causing agents that could at least in some cases necessitate defensive responses by the ants. Together, our ecological and experimental data support our conclusion that *Trichoderma* spp. are opportunistic pathogens of *T. septentrionalis* fungus gardens.

Metabolomics analyses of lab-reared fungus gardens inoculated with *T. septentrionalis*-isolated *Trichoderma* spp. revealed the presence of peptaibols in infected gardens (Fig. 2*C*, *<u>SI Appendix</u>*, Fig. S7, S9-11). Wild fungus gardens contained similar peptaibols, indicating their ecological relevance, consistent with peptaibol-producing *Trichoderma* being an opportunistic pathogen of *T. septentrionalis* fungus gardens (Fig. 1*B*). In our laboratory experiments, two of the *Trichoderma* fractions with the highest abundance of peptaibols, fractions D and E, induced some of the strongest *T. septentrionalis* weeding response (Fig. 3), and purified peptaibols also induced a weeding response similar to that of the crude *Trichoderma* extract (*<u>SI Appendix</u>*, Fig. S14). Given that the *Trichoderma* fractions and purified peptaibols were derived from different strains and represent a diversity in peptaibol composition, the similarities in the observed ant behavioral activity are thus likely to be unrelated to specific peptaibols but rather generally attributable to the peptaibol class of metabolites. Furthermore, the presence of peptaibols in environmental fungus gardens lends credence to their ecological relevance and together with the experimental data parallels the logic of Koch’s postulates, suggesting that peptaibols are likely produced during *Trichoderma* infections of ant fungus gardens and induce ant defensive behaviors in response.

Peptaibols are produced by fungi in the order Hypocreales, especially by members of the mycoparasitic family *Hypocreaceae*, which contains both *Trichoderma* and *Escovopsis*, a specialized ant fungus garden pathogen (41, 42). Peptaibols have been hypothesized to be important for mycoparasitism (55, 56), which we show here can include infections of ant fungus gardens. Interestingly, the genes needed to synthesize peptaibols are encoded within the genome of *Escovopsis* (57), and *Escovopsis* has been shown to induce a strong ant weeding response in tropical leaf-cutting fungus-growing ants (17), similar to the *T. septentrionalis* weeding behaviors that we observed in response to *Trichoderma* and its peptaibols. We therefore hypothesize that peptaibol-induced weeding behaviors are conserved among diverse fungus-growing ants and may reflect an ancient means of pathogen detection and defense. Future work should test if ant weeding intensity is directly correlated with pathogen load, virulence, peptaibol production, or other contributing environmental factors, and also compare the behavioral responses of diverse fungus-growing ant species to diverse fungal pathogens with varying levels of virulence and specialization towards ant fungus gardens, e.g., as in (58).

This study demonstrates how *T. septentrionalis* ants protect their cultivar mutualist from opportunistic *Trichoderma* pathogens by sensing and responding to peptaibols as specific molecular cues that induce an ant weeding response. These cues included two new bioactive peptaibol metabolites that we identified in this study. Future research will investigate whether ants directly sense peptaibols or indirectly respond to an intermediate signal produced by the cultivar in response to peptaibols, in addition to characterizing other potential signaling molecules that are unlikely to be present in our *Trichoderma* extracts (e.g., volatile compounds). In contrast to the canonical logic of host immune responses, in which hosts directly respond to infections, *T. septentrionalis* responses to peptaibol signaling molecules comprise an extended defense response whereby *T. septentrionalis* ants respond to infections of their cultivar mutualist. Such extended defense responses may be a widespread but poorly recognized mechanism that increases host health indirectly by preventing harm to their beneficial symbionts.

## Materials and Methods

### Colony collections

Colonies of *T. septentrionalis* were collected during the summers of 2014-2017 from state parks and forests throughout the eastern USA (*<u>SI Appendix</u>*, Table S1) and transferred to the Klassen laboratory as described elsewhere (59, 60). Collection permits include: Louisiana Department of Wildlife and Fisheries permit WL-Research-2016-10, Florida Department of Agriculture and Consumer Services unnumbered Letter of Authorization, Georgia Department of Natural Resources State Parks & Historic Sites Scientific Research and Collection Permit 032015, Georgia Department of Natural Resources Wildlife Resources Division unnumbered Letter of Authorization, North Carolina Division of Parks and Recreation Scientific Research and Collecting Permit 2015_0030, State of New Jersey Department of Environmental Protection Division of Parks and Forestry State Park Service Unnumbered Letter of Authorization, New York State Department of Environmental Conservation License to Collect or Possess: Scientific # 915; County of Suffolk Department of Parks, Recreation and Conservation “Permit for Research in Suffolk County Parklands” (unnumbered), and USDA permit P526P-14-00684 for the transport of colonies. Fungus garden samples were frozen directly in the field in either dimethyl sulfoxide-ethylenediamine tetraacetic acid-saturated salt solution (DESS) (59) for mycobiome sequencing (dataset “Environmental ITS2”; *<u>SI Appendix</u>*,Table S1) or without added preservative for metabolomic analysis (dataset “Environmental LC-MS/MS”; *<u>SI Appendix</u>*,Table S1). Due to the prospective nature of these collections, only a subset of colonies was used for both analyses. Three additional colonies were collected from New Jersey during the summer of 2014 and used to isolate the *Trichoderma* strains used in this study (dataset “*Trichoderma* isolations”; *<u>SI Appendix</u>*, Table S1). Two more colonies were collected during the summer of 2018 from New Jersey and used for laboratory infection experiments (dataset “*Trichoderma*/Time-course infection”; *<u>SI Appendix</u>*, Table S1). Colonies were collected in 2019 under permit North Carolina Division of Parks and Recreation Scientific Collection and Research Permit 2019_0322 and used for preliminary chemical extract experiments in 2019 (dataset “2019 extract tests”; *<u>SI Appendix</u>*, Table S1). Further chemical extract testing was performed with additional colonies collected from New Jersey during the summer of 2020 (dataset “2020 extract tests”; *<u>SI Appendix</u>*, Table S1).

### Trichoderma isolations

Fungal pathogen strains JKS001879, JKS001884, and JKS001921 (*<u>SI Appendix</u>*, Table S2) were isolated from pieces of recently collected fungus gardens with their ants removed. Non-cultivar fungi that overgrew these unprotected fungus gardens were collected using tweezers and plated on Potato Dextrose Agar (PDA, Difco) + antibiotics (ABX, 50 mg/L penicillin and 50 mg/L streptomycin; both Fisher Scientific) and grown at 25 °C. Fresh hyphal growth from the edge of each colony was sub-cultured several times to obtain morphologically and microscopically pure cultures. These strains were then identified by Sanger sequencing of several marker genes including the full ITS region, translation elongation factor 1-α (TEF1), and the RNA polymerase II subunit RPB2. DNA was extracted (see below) and PCR amplified using primers ITS1/ITS4, EF1/EF2, or fRPB2-5f/fRPB2-7cr (*<u>SI Appendix</u>*, Table S3). The ITS region was amplified using thermocycling conditions: 3 min at 95 °C, 30 cycles of 30 s at 95 °C, 45 s at 50 °C, and 45 s at 72 °C, with a final extension for 2 min at 72 °C [modified from (61)]. The TEF gene was amplified following (62). The RPB2 gene was amplified using thermocycling conditions: 3 min at 95 °C, 15 cycles of 45 s at 95 °C, 1 min at 60 °C and decreasing by 0.5 °C each cycle, and 1 min at 72 °C, followed by 30 cycles of 95 °C for 45 s, 54 °C for 1 min, and 72 °C for 1 min, with a final extension at 72 °C for 5 min [modified from (62, 63)]. Amplified products were purified using Agencourt AMPure XP beads (Beckman Coulter) following the manufacturer instructions for PCR purification. Purified PCR products were then sequenced at Eurofins Genomics (Louisville, KY) and resulting data were analyzed using Geneious Prime v. 2019.1.3 to generate consensus sequences that were then processed with MIST (64) (http://mmit.china-cctc.org/index.php accessed: February 22, 2022) in the order ITS, TEF1, and then RPB2. This protocol was also used to sequence the full ITS genes of the pathogens that bloomed in the *Trichoderma* infection experiment (described below) to confirm that each matched the inoculated strain.

### Trichoderma assays

#### Spore isolation

*Trichoderma* strains JKS001879, JKS001884, and JKS001921 were grown on PDA+ABX plates at 25 °C for ∼1 week until green colored growth covered the plate, indicating sporulation. Spores were isolated by flooding each plate with up to 5 mL of sterile phosphate buffered saline pH 7.4 (PBS) and gently agitating with a sterile loop to suspend the spores. Avoiding the remaining hyphae, suspensions were collected from each plate and diluted with PBS until a phase separation between the heavy particulates and floating hyphae could be visualized after vortexing. The center phase, enriched with spores, was collected and visualized using 1000x oil immersion light microscopy to verify the presence of spores and general absence of hyphae. Spore concentrations were measured using the optical density at 600 nm (OD_600_), and spore suspensions were diluted to OD_600_ 0.2 and stored at 4 °C.

#### Trichoderma infection

Subcolonies of *T. septentrionalis* colony JKH000285 were created using small portions of fungus garden (∼20 cm^3^) in small (7.5 x 7.5 x 3 cm) plaster-lined plastic boxes, manipulating each garden as little as possible. Spore suspensions were warmed to room temperature and 500 µL of either *Trichoderma* sp. JKS001879, JKS001884, or JKS001921 spore suspension or PBS as a negative control were dripped onto fungus gardens, either with or without 8 worker ants present, mirroring previous work that describes other fungus garden pathogens (6, 16). Although treatments lacking ants represent an extreme perturbation that is unlikely to occur naturally, they provide a unique experimental means to differentiate the relative contributions of ants and fungus gardens to colony defense. Pictures were taken at 0, 1, 3, and 4 days post-treatment (*<u>SI Appendix</u>*, Fig. S6, Data Availability). Fungus garden samples were frozen at -80 °C in DESS or without buffer for DNA sequencing and metabolomic analysis, respectively, 17 days post treatment. <u>Time-course infection</u>: *Trichoderma* sp. JKS001884 was chosen for further investigation because it grew the fastest and produced the most spores in the initial infection experiment. Subcolonies of *T. septentrionalis* colony JKH000292 were created as described above, and treated with *Trichoderma* sp. JKS001884 (500 µL of spores suspended in 1X PBS, OD_600_ 0.2) or 500 µL PBS alongside no-treatment controls, both with and without 12 worker ants present and with parallel replicates. Treatments were applied by spraying each fungus garden from ∼2” above using an atomizer (Teleflex MAD300). Four ants and ∼1/3 of the fungus garden were frozen at -80 °C in DESS or without buffer dry for DNA and metabolomic analysis, respectively, at each time point (days 0, 2, and 4). Waste from +ants subcolonies was also sampled at day 4 post-treatment. Pictures were taken at each sampling time point.

#### *Trichoderma* extract experiments

Ant weeding behavioral responses to *Trichoderma* extracts (extraction procedures described below) were assessed qualitatively in 2019 (August 2 - September 30) using video analysis of ant subcolonies created as described above from three different *T. septentrionalis* colonies and with 10 worker ants each. Weeding severity was determined by visually comparing the amount of waste removed from subcolony fungus gardens and discarded around the edges of the box, and thus the extent of fungus garden disturbance in response to treatment. Treatment with *Trichoderma* crude extract (10 mg/mL) induced more weeding compared to DMSO and NT controls (*<u>SI Appendix</u>*, Movie S1). During the 2020 season (July 16 - September 24) waste generated by ant weeding was quantified by weighing the mass of waste produced by ants in response to various *Trichoderma* extract/compound treatments. Subcolonies were created as described above using 10 worker ants per subcolony, except side chambers were omitted due to the short duration of each experiment and Plaster of Paris (50 g) was poured into each box 8-48 h before each experiment to minimize variation in humidity. Immediately prior to each experiment, the plaster was saturated with deionized water to create controlled, high humidity conditions for each box. Each box was weighed before and after adding fungus gardens to calculate fungus garden mass. Two hundred μL of each crude extract (10 mg/mL), fraction (1 mg/mL), pure compound (0.25 mg/mL), or solvent-only control (*<u>SI Appendix</u>*, Table S4) was dripped onto the fungus garden surface. Fractions and pure compounds were diluted to approximate their concentrations in extracts of pure *Trichoderma* cultures. A still image of each treated fungus garden was taken before adding ants, ants were added, and then timelapse photos were taken every 5 s for 24 h to document ant behaviors. After 24 h ants were removed and a still image was taken before and after the waste created by the ants was collected into preweighed 2 mL tubes. Tubes were capped to prevent dehydration and then weighed again to calculate the total waste mass. Timelapse photography for each experiment was converted into movies using FFMPEG (http://ffmpeg.org) with the command:

~~~
ffmpeg -r 30 -f image2 -pattern_type glob -i ’*.JPG’ -s 4000x3000
-c:v mjpeg -q:v 10 ./output_timelapse.ffmpeg.30fps.q10.mov
~~~

Surprisingly, our initial experiments conducted early in the 2020 season did not recapitulate the previous induction of ant weeding behavior in response to crude *Trichoderma* extracts seen in 2019. As the season progressed (late August through September), colonies began responding to the treatments, and thus only experiments where the *Trichoderma* extract weeding response exceeded that of the negative controls were included in the final dataset (trials 9-16, and the replicate using colony JKH000396 in trial 17; *<u>SI Appendix</u>*, Table S4) We suspect this variability in response is due to seasonal and intercolony variability in *T. septentrionalis* activity, although further testing is necessary to confirm this. Ant weeding was then compared between treatments by averaging waste production per treatment across colony replicates.

### DNA extraction and sequencing

#### DNA extraction

DNA was extracted from approximately 0.05 g of fungus garden stored in DESS or fresh hyphae from *Trichoderma* culture plates as described in (59), except that the fungus garden samples used to generate the environmental ITS2 dataset were first washed thrice with sterile 1X PBS to remove PCR inhibitors and DNA was precipitated for 1 h at room temperature. All other samples were extracted using a modification of (59) wherein the extraction volume was halved (to 250 µL), 80% ice cold ethanol was used to wash DNA pellets, DNA pellets were air dried on a 37 °C heat block, DNA was incubated and gently agitated in 100 µL of nuclease-free water for ≥ 1 h at 37 °C to resuspend the DNA pellet and then quantified using the Qubit dsDNA high-sensitivity assay protocol and a Qubit 3.0 fluorimeter (Invitrogen, Carlsbad CA). Samples in the environmental ITS2 dataset were cleaned by excising 25-30 μL of each PCR product from 1% agarose gel using a sterile scalpel and the QIAGEN QIAquick Gel Extraction Kit following the manufacturer’s instructions.

#### Fungal ITS2 community amplicon sequencing

Before fungal community sequencing, DNA samples were first pre-screened for amplification using primers fITS7 and ITS4 targeting the ITS2 region of the fungal rRNA gene (*<u>SI Appendix</u>*, Table S3). Three μL (6 ng on average) of template DNA was added to 5 μL 5x GoTaq Green Reaction Mix Buffer (Promega, Madison, Wisconsin, USA), 0.625 units of GoTaq DNA Polymerase (Promega, Madison, Wisconsin, USA), 0.2 mM each dNTP, 0.15 μM of each primer, 0.464 μg/μL bovine serum albumin (BSA, New England BioLabs Inc. Ipswich, Massachusetts), and 1.5 mM of MgCl_2_, to which nuclease-free water was added to a volume of 25 μL. Thermocycling conditions were: 3 min at 95 °C, 30 cycles of 30 s at 95 °C, 45 s at 50 °C, and 45 s at 72 °C, followed by a 2 min final extension at 72 °C. Gel electrophoresis was used to confirm the expected band size of 300-500 bp. DNA extracts that failed to amplify were diluted 10-fold and re-screened, continuing to a maximum dilution factor of 1:50.

DNA extracts that successfully amplified during pre-screening and their corresponding negative control extracts were prepared for ITS2 community amplicon sequencing. Six μL (12 ng on average) of DNA from each extract was added to 10 μmol of each Illumina-barcoded version of fITS7 and ITS4 primers, 5 μL of 10x AccuPrime buffer (Invitrogen, Carlsbad, California), 0.4 mM MgSO_4_, 0.16 μg/μL BSA (New England BioLabs Inc. Ipswich, Massachusetts), 50 μM each dNTP, 0.2 μM of non-barcoded primers fITS7 and ITS4, and 1.25 units of AccuPrime polymerase. Nuclease-free water was added to reach a final volume of 50 μL per reaction. A PCR negative control using 6 μL nuclease-free water as template was included for each 96-well plate processed. This PCR master mix was split into triplicates (16.7 μL each reaction) using an epMotion 5075 liquid handling robot (Eppendorf, Hamburg, Germany). Thermocycler conditions were 2 min at 95 °C, 30 cycles of 95 °C for 15 s, 55 °C for 1 min, and 68 °C for 1 min, followed by final extension at 68 °C for 5 min before an indefinite hold at 4 °C. Triplicate PCR reactions were then re-pooled using the epMotion 5075 and DNA concentrations were quantified using a QIAxcel Advanced capillary electrophoresis system (QIAgen, Hilden, Germany). An equal mass of DNA (up to a maximum volume of 12 μL) was pooled into a single library, which was cleaned using Mag-Bind RXNPure plus beads (OMEGA, Norcross, GA) in a 1:0.8 ratio of sequencing library to bead volume. Cleaned libraries were adjusted to 1.1 ng/μL+/- 0.1 ng/μL using a Qubit dsDNA high-sensitivity assay and Qubit 3.0 fluorometer. Sequencing was performed using Illumina Miseq V2 2 x 250 bp reads.

### ITS2 community amplicon sequence analysis

For the Environmental ITS2 dataset, (NCBI accession: PRJNA763335, *<u>SI Appendix</u>*, Dataset S1) forward reads from 90 samples were processed using Trimmomatic v0.39 (65) with options SLIDINGWINDOW 5:20 and MINLEN:125 and then processed in R v.3.6.3 using the DADA2 v1.16.0 (66) ITS workflow (https://benjjneb.github.io/dada2/ITS_workflow.html, accessed May 26, 2021) except using only forward reads. All negative controls produced 0 reads so decontamination methods were not necessary for this dataset. Following DADA2, amplicon sequence variants (ASVs) were inputted into ITSx v1.1.3 (67) to extract the ITS2 region. ASVs without an ITSx-detected ITS2 region were filtered (n = 6), and ASVs with identical ITS2 regions (n = 41) were merged using the Phyloseq v1.26.1 (68) merge_taxa() command. Remaining ASVs were then taxonomically classified using the UNITE database (69) general FASTA release for Fungi 2 v.8.2 (https://doi.org/10.15156/BIO/786369). Read sets were rarefied to 12,531 reads, which removed 7 samples and 211 ASVs. ASVs not classified at the phylum level (n = 20) were removed using Phyloseq’s subset_taxa() command and ASVs that were not present at ≥ 1% abundance in at least one sample (n = 381) were assigned as “Other” using the metagMisc v.0.04 (https://github.com/vmikk/metagMisc) command phyloseq_filter_sample_wise_abund_trim(). The remaining 35 ASVs were assigned to 10 taxa: “Cultivar fungus”, *Meyerozyma*, *Trichoderma*, unclassified *Stephanosporaceae*, *Lecanicillium*, *Penicillium*, *Oberwinklerozyma*, *Cladosporium*, *Xylaria*, and *Scleroderma*. “Cultivar fungus” was manually defined as all taxa classified as genus *Leucocoprinus*, *Leucoagaricus*, or unclassified *Agaricaceae* confirmed as being best BLAST (70) matches to a higher attine cultivar fungus. For all ITS2 community analyses, ASVs were not classified past rank genus because the ITS2 region is too short to delineate species for many fungi, including *Trichoderma*.

For the time-course infection dataset (NCBI accession: <u>PRJNA743045</u>, *<u>SI Appendix</u>*, Dataset S2), sequencing resulted in 46 ITS2 community amplicon readsets including 4 negative controls. The number of reads per sample ranged from 6 to 23,953. This data was analyzed in the same way as the environmental ITS dataset except that after ITSx treatment and taxonomy assignment the prevalence protocol for decontam v1.12.0 (71) was used with a default threshold of 0.1 to determine if any ASVs were contaminant sequences. No contaminants were detected so negative controls were removed from further processing. This dataset was rarefied to 4,099 reads, which removed three ASVs and two samples. Two ASVs were removed that were unclassified past the Kingdom level, which left 31 unique ASVs. ASVs were then filtered to remove very rare taxa (ASVs present at < 1% abundance in at least one sample readset). This resulted in 14 unique ASVs that belonged to five taxa: “Cultivar fungus”, *Trichoderma*, *Candida*, *Vanrija*, and *Penicillium*.

### Extraction and fractionation of *Trichoderma* strain JKS001884

All solvents were HPLC grade and purchased from Sigma Aldrich (St. Louis, MO). Agar plates were cut into small pieces and extracted with 2:1 ethyl acetate:methanol (EtOAC:MeOH) three times and sonicated for 60 s. Solvent was collected after each extraction, filtered under vacuum, and dried using rotary evaporation.

Fractionation of *Trichoderma* strain JKS001884 was performed twice. Initial fractionation of a portion of MB1081 (1.48 g) began by dissolving it in a minimum amount of 50% aqueous MeOH, which was then loaded onto a Discovery^®^ DSC-18 SPE cartridge (10 g/50 mL). The column was conditioned and equilibrated with 150 mL portions of water, MeOH, and 10% aqueous MeOH prior to sample addition. Using a stepwise gradient, the sample was eluted under negative pressure using (A) 10% aqueous MeOH, (B) 25% aqueous MeOH, (C) 50% aqueous MeOH, (D) 75% aqueous MeOH, and 100% MeOH (E) that were eluted using 150, 150, 250, 300, and 300 mL portions, respectively, followed by a 100% acetonitrile (ACN) fraction (F), an organic wash with dichloromethane (DCM)/acetone/MeOH (G), and a final aqueous wash (H). The eight fractions (MB1081A-H) were dried under reduced pressure. For the second fractionation, crude extract MB1084 (413.0 mg) was dissolved in a minimum amount of 50% aqueous MeOH and then loaded onto a Discovery^®^ DSC-18 SPE cartridge (10 g/50 mL). Column conditions and fractionation were as described above for fractions A-E, except the sample was eluted with 135 mL of each solvent and no 100% ACN fraction was collected. An organic wash of 1:1:1:1 EtOAc:DCM:acetone:MeOH was collected (MB1084F), along with a final aqueous wash (MB1084G). The seven fractions (MB1084A-G) were dried under reduced pressure. All fractions were then evaluated using LC-MS/MS and in ant behavior assays.

### Molecular networking of fungus garden samples

Fungus garden samples from both field-collected and lab-raised gardens were extracted with 2:1 DCM:MeOH three times. Samples were sonicated, manually macerated with a metal spatula, and incubated in solvent for 15 min at room temperature, after which solvent was filtered, combined, and dried under nitrogen [see (72) for details].

#### LC-MS/MS conditions for molecular networking

Data acquisition corresponding to datasets MSV000082295, MSV000088621, and MSV000083723 were prepared as follows: samples were resuspended to a concentration of 1 µg/mL in 100% MeOH containing 2 µM sulfamethazine as an internal standard and LC-MS/MS analysis was performed using an UltiMate 3000 UPLC system (Thermo Scientific) with a Kinetex 1.7 µm C_18_ reversed phase UHPLC column (50 X 2.1 mm) and Maxis Q-TOF mass spectrometer (Bruker Daltonics) equipped with an ESI source. The column was equilibrated with 5% solvent B (LC-MS grade ACN, 0.1% formic acid) for 1 min, followed by a linear gradient from 5% B to 100% B over 8 min, and held at 100% B for 2 min. Then, the column was re-equilibrated via 100%–5% B in 0.5 min and maintained at 5% B for 2.5 min. A flow rate of 0.5 mL/min was used throughout the run. MS spectra were acquired in positive ion mode in the range of 100-2000 *m/z*. A mixture of 10 mg/mL each of sulfamethazine, sulfamethizole, sulfachloropyridazine, sulfadimethoxine, amitriptyline, and coumarin was run after every 96 injections for quality control. An external calibration with ESI-Low Concentration Tuning Mix (Agilent technologies) was performed prior to data collection and the internal calibrant hexakis(1H,1H,2H-perfluoroethoxy)phosphazene (CAS 186817-57-2) was used throughout the runs. A capillary voltage of 4500 V, nebulizer gas pressure (nitrogen) of 2 bar, ion source temperature of 200 °C, dry gas flow of 9 L/min, and a spectral rate of 3 Hz for MS1 and 10 Hz for MS2 was used. For acquiring MS/MS fragmentation, the 5 most intense ions per MS1 were selected, MS/MS active exclusion parameter was set to 2 and to release after 30 s, and the precursor ion was reconsidered for MS/MS if the current intensity/previous intensity ratio was higher than 2. An advanced stepping function was used to fragment ions, applying specified values for transferring ions into the collision cell per scan time according to Supplementary Table S5. CID energies for MS/MS data acquisition were used according to Supplementary Table S6.

#### Molecular networking

Networking analyses were performed with the molecular networking workflow (https://ccms-ucsd.github.io/GNPSDocumentation/networking/) on the GNPS website [http://gnps.ucsd.edu; (73)]. The data were filtered by removing all MS/MS fragment ions within ±17 Da of the precursor *m/z*. MS/MS spectra were window filtered by choosing only the top 6 fragment ions in the ±50 Da window throughout the spectrum. The precursor ion mass tolerance was set to 0.02 Da and a MS/MS fragment ion tolerance of 0.02 Da. A network was then created where edges were filtered to have a cosine score > 0.7 and ≥ 6 matched fragment ions. Edges between two nodes were kept in the network only if each node appeared in each other’s respective top 10 most similar nodes. Finally, the maximum size of a molecular family (also called a cluster or connected component of a plot) was set to 100, and the lowest scoring edges were removed from molecular families until the molecular family size was below this threshold. The spectra in the network were then searched against all GNPS spectral libraries. The library spectra were filtered in the same manner as the input data. All matches kept between network spectra and library spectra had a score > 0.7 and ≥ 6 matched fragment ions. Molecular networks were visualized using Cytoscape 3.7.2 (74). The molecular networking results corresponding to the fungus gardens collected from across the Eastern of the USA can be accessed at https://gnps.ucsd.edu/ProteoSAFe/status.jsp?task=428481037da64c989cbac2f84d866fba. The molecular networking results corresponding to the colonization experiment with isolates JKS001879, JKS001884, and JKS001921 can be accessed at https://gnps.ucsd.edu/ProteoSAFe/status.jsp?task=deaddf07450e435f91b3401cf4989f9f. The molecular networking results corresponding to the time-course colonization with JKS001884 strain can be accessed at https://gnps.ucsd.edu/ProteoSAFe/status.jsp?task=08e66fea0bea42e7b0655b1f877e5c67.

#### Feature-based molecular networking

Feature finding was performed with the open source MZmine software (75, 76), version 2.34 using the following settings: mass detection (MS1 detection at noise level of 1.0E4, MS2 detection at noise level of 1.0E2); chromatogram building (minimum time span: 0.01 min; minimum height: 3.0E4; *m/z* tolerance: 25 ppm); deconvolution [algorithm: baseline cut-off (minimum peak height: 1.0E4; peak duration range 0.01-1.0 min; baseline level: 1.0E2); *m/z* range for MS2 scan pairing: 0.01 Da; RT range for MS2 scan pairing: 0.3 min]; isotopic peak grouper (*m/z* tolerance: 25 ppm; RT tolerance: 0.2 min; maximum charge: 2); alignment (join aligner, *m/z* tolerance: 25 ppm; weight for *m/z*: 75; weight for RT: 25; RT tolerance: 0.2 min); gap filling (peak finder, intensity tolerance: 1%; *m/z* tolerance: 25 ppm; RT tolerance: 0.2 min; RT correction: checked); peak filter (peak area 1.0E4-1.0E7); peak row filtering to export .mgf file to GNPS (minimum peaks in a row: 2; RT: 1.00-14.00 min).

A molecular networking analysis of the time-course monitoring of pathogen colonization with strain JKS001884 of *T. septentrionalis* fungus gardens (Fig. S7-S8) was created with the feature-based molecular networking workflow (77). The data were filtered and the spectra were searched as described for molecular networking above. GNPS job link: https://gnps.ucsd.edu/ProteoSAFe/status.jsp?task=472a3fc2b798426dbb4361fda9bdbb7c.

### Metabolomics and statistical analyses of *Trichoderma* extracts and fractions

#### Untargeted MS data acquisition

MS data were acquired using a Waters Synapt G2-Si coupled to an Acquity UPLC system with an Acquity UPLC HSS T3 VanGuard pre-column (2.1 x 5 mm, 1.8 μM) and an Acquity UPLC HSS T3 column (2.1 x 150 mm, 1.8 μM) with a flow rate of 0.45 mL/min. Samples were prepared in a 1:1 mixture of MeOH/water and injected three times for technical replication, using a gradient elution with mobile phases A (water with 0.1% formic acid) and B (ACN with 0.1% formic acid) using the following conditions: 0.5 min hold at 95% A and 5% B, 3.5 min ramp to 60% B, 4 min ramp to 100% B, 1 min hold at 100% B, 0.2 min ramp back to 5% B and a re-equilibration hold at 5% B for 1.8 min. Spectrometric data were acquired using MS^E^ data-independent acquisition (DIA) with continuous MS survey scanning ranging from *m/z* 50-2,000 in positive-resolution mode. Data were acquired using the following parameters: 2 kV capillary voltage, 100 °C source temperature, 20 V sampling cone, 2 V collision energy, 800 L/h desolvation gas flow, and 80 V source offset. Real-time mass correction used a lockspray of 400 pg/μL leucine enkephalin solution (1:1 MeOH:water solution with 0.1% formic acid).

#### MS data pre-processing

MS data was processed using Progenesis QI (Nonlinear Dynamics, Milford, MA) to create a peak list for statistical analyses. An injection of the crude extract was selected as the alignment reference since all fractions share features with the crude extract. Peak picking parameters included fragment sensitivity of 0.2% of the base peak and adducts included M+H, M+2H, 2M+2H, 3M+3H, and M+Na. Export included compound ID, *m/z*, retention time, and raw abundance.

#### Metabolomics data processing, filtering, and statistical analyses

The peak list and fragment database were exported from Progenesis and analyzed with MPACT (78). Data were filtered using solvent blank, mispicked peak, reproducibility, and in-source fragment filters (*SI Appendi*x, Fig. S41). Blank filtering using MeOH blanks was applied based on a relative group parsing threshold of 0.05. A minimum reproducibility threshold of 0.5 median coefficient of variation (CV) among technical replicates was applied. The presence or absence of features in a group of samples was determined using a relative abundance threshold of 0.05 compared to the sample group in which a feature was most abundant. This threshold was applied during blank filtering to remove features whose abundance in solvent blanks was greater than 5% of their abundance in experimental samples. A 0.95 Spearman correlation threshold was used for identifying and removing in-source ion clusters. A correction for mispicked peaks was applied with a ringing mass window of 0.5 AMU, isotope mass window of 0.01 AMU, retention time window of 0.005 min, and a maximum isotopic mass shift of 4 AMU.

#### Analysis of peptaibol abundance

The Progenesis peak list was used to determine the percent abundance of each feature relative to the total ion abundance for each fraction (Fig. 4C). The percent abundance of each feature was obtained by averaging across technical replicates and then dividing by the average total ion abundance for each fraction, with heat map coloration ranging from 0-50%, established based on the most abundant feature with *m/z* 1197.7557, which constituted 47% of fraction E.

### Collection and fermentation of *Trichoderma arundinaceum* (MSX 70741) and *T. albolutescens* (MSX 57715)

#### Collection and identification of fungal strains

All peptaibols used for compound isolation were isolated from fungal cultures provided by Mycosynthetix, Inc., specifically *Trichoderma arundinaceum* (strain MSX70741) and *T. albolutescens* (strain MSX57715). Molecular taxonomical identification of both strains was performed using phylogenetic analysis on *RPB2* and *tef-1* sequence data following procedures described by Raja et al. (79). Full details pertaining to the collection of these samples and their identification using molecular approaches were published previously (80).

#### Fungal strain fermentation

Strains MSX70741 and MSX57715 were fermented following the procedures outlined previously (80, 81). Briefly, a seed culture of each fungal strain was grown on a malt extract agar slant for 7 d. Subsequently, a piece of mycelium was transferred into YESD media and incubated for seven days at 22 °C with shaking at ∼125 rpm. Separately, each seed culture was next propagated into either 250 mL Erlenmeyer flasks with 10 g of rice and 20 mL of water or 2.8 L Fernbach flasks containing 150 g of rice and 300 mL of water with vitamin solution; in all cases the rice was first autoclaved at 121 °C. The cultures were incubated at 22 °C for 2-3 weeks prior to extraction and purification of peptaibols (81).

### Extraction, isolation, and identification of compounds 1 and 2

Compounds **1** and **2** were obtained from strain MSX57715, following standard procedures (80, 81). Briefly, to the large-scale solid fermentation of strain MSX57715, a solution of chloroform (CHCl_3_):MeOH (1:1) was added, and the mixtures were shaken for 16 h at 100 rpm in a reciprocating shaker. The solutions were vacuum filtered, combined, and equal volumes of water and CHCl_3_ were added to a final volume of 2 L, whereupon the mixture was stirred for 2 h and then transferred into a separatory funnel. The organic layer was drawn off and evaporated to dryness. The extract was next defatted by partitioning between 300 mL of a mixture of 1:1 MeOH:ACN and 300 mL of hexanes in a separatory funnel. The bottom layer was collected and evaporated to dryness to give 8.0 g of extract. The defatted extract (7.9 g) was adsorbed onto a minimal amount of Celite 545 (Acros Organics). This material was fractionated via flash chromatography on a 120 g RediSep Rf Gold Si-gel column, using a gradient solvent system of CHCl_3_:MeOH at an 85 mL min^−1^ flow rate and 28.3 column volumes over 63.9 min, to afford five fractions (i.e., F1–F5). Part of fraction F5 (∼250 mg) was subjected to preparative HPLC using a Synergy column, Max-RP, 4 μm, 250 × 21.2 mm, and a gradient system initiated with 40:60 ACN:water (0.1% formic acid) to 100% ACN over 30 min at a flow rate of 21.0 mL min^−1^ to afford compounds **1** (6.6 mg) and **2** (7.0 mg).

Trichokindins VIII and IX (i.e., compounds **1** and **2**, respectively) were isolated from strain MSX57715. Their ^1^H NMR spectra were similar to what was observed with peptaibols previously isolated from the same strain and other *Trichoderma* species (80, 81). The HRESIMS data for **1** and **2** (*<u>SI Appendix</u>*, Figures S20 and S29) suggested molecular formulae of C_82_H_144_N_20_O_22_ and C_81_H_142_N_20_O_22_ , respectively, indicating an index of hydrogen deficiency of 21 for both compounds. These formulae were structurally similar to trichokindins I-V, a group of 18 residue peptaibols isolated from *T. harzianum* (82). However, the MS/MS fragmentation data for **1** and **2** were not a perfect match for the amino acid sequence reported for either of those molecules. For example, in **2** the full scan spectrum showed characteristic ions at *m*/*z* 1748.0651 [M + H]^+^, 1108.6349 (b_12_^+^), 640.4379 (y_6_^+^). Additional fragmentation of the b_12_^+^ ion provided information about the successive losses of Aib^12^ Leu^11^, Gly^10^, Aib^9^, Val^8^, Aib7-Gln^6^, Aib^5^-Aib^4^-Ala^3^, and Ser^2^-Aib^1^-Ac. MS/MS of fragment y_6_^+^ generated a sequence of ions corresponding to the loss of Leuol^18^, Gln^17^, Iva^16^, Aib^15^, and Leu^14^-Pro^13^. These data suggested that compound **2** was Ac-Aib^1^-Ser^2^-Ala^3^-Aib^4^-Aib^5^-Gln^6^-Aib^7^-Val^8^-Aib^9^-Gly^10^-Leu^11^-Aib^12^-Pro^13^-Leu^14^-Aib^15^-Iva^16^-Gln^17^-Leuol^18^. The planar structure of **2** was confirmed based on interpretation of 1D and 2D NMR data, including the use of ^1^H, ^13^C, COSY, HSQC, HMBC, and TOCSY experiments. Key correlations that supported the proposed structure are highlighted in *<u>SI Appendix</u>*, Figure S28. The presence of an isovaline at position 16 (Iva^16^) was supported by a triplet at δ_H_/δ_C_ (0.69, t, *J* = 7.5 Hz/ 7.2 ppm), which displayed correlations in the TOCSY and COSY experiments with hydrogens of a methylene group at δ_H_/δ_C_ (1.66, 2.13/26.0). In particular, the HMBC correlations of Iva^16^ γ-CH_3_ and β_2_-CH_3_ with the Iva^16^ α carbon supported this assignment. The structure of compound **1** was established similarly, combining spectroscopic and spectrometric data. The only differences were observed in the full scan MS spectrum, particularly for ions [M + H]^+^ and b_12_^+^, which displayed *m*/*z* values at 1762.0803 and 1122.6499, respectively, which were 14 Da more than those in compound **2**, indicating the incorporation of an extra methylene. Additionally, the ^1^H NMR spectra displayed signals for two triplet at δ_H_/δ_C_ 0.72/7.2 and 0.70/7.2 ppm, indicative of an additional Iva residue. Based on the analysis of the b_12_^+^ MS/MS spectrum, this residue was located at position five. Considering all the data obtained from mass spectrometry and 1D and 2D NMR experiments, the amino acid sequence of **1** was established as Ac-Aib^1^-Ser^2^-Ala^3^-Aib^4^-Iva^5^-Gln^6^-Aib^7^-Val^8^-Aib^9^-Gly^10^-Leu^11^-Aib^12^-Pro^13^-Leu^14^-Aib^15^-Iva^16^-Gln^17^-Leuol^18^. Compounds **1** and **2** were ascribed the trivial names trichokindins VIII and IX, respectively. These peptaibols share ∼70% of the core structure with other trichokindins, differing mainly by replacement of residues of Leu^8^, Ala^10^, and Aib^16^ in trichokindins I-VII by Val^8^, Gly^10^, and Iva^16^ in compounds **1** and **2**, respectively (82).

**Trichokindin VIII (1):** White powder; HRESIMS *m*/*z* 1748.0651 [M + H]^+^; calcd. for C_81_H_143_N_20_O_22_ 1748.0680. Δ = ‒1.7 ppm, Ω = 21. Structural characterization details are provided in *<u>SI Appendix</u>*, Table S7 and Figures S23 to S31.

**Trichokindin IX (2):** White powder; HRESIMS *m*/*z* 1762.0803 [M + H]^+^; calcd. for C_82_H_145_N_20_O_22_ 1762.0837. Δ = ‒1.9 ppm, Ω = 21. Structural characterization details are provided in *<u>SI Appendix</u>*, Table S7 and Figures S32 to S40.

## Supporting information

SI_revised

## Data availability

Sanger sequencing data generated in this study to identify *Trichoderma* isolates can be accessed from the National Center for Biotechnology Information (NCBI) GenBank (https://www.ncbi.nlm.nih.gov/genbank/) using accession numbers OM967104-OM967106 (ITS), ON113306-ON113308 (*RPB2*), and ON364341-ON364343 (*TEF1*). Fungal community sequencing data was deposited in NCBI under BioProject <u>PRJNA763335</u> and <u>PRJNA743045</u> for the wild fungus gardens and laboratory *Trichoderma* infections, respectively. Metabolomic data sets used in this work were deposited in the <u>Mass</u> Spectrometry Interactive <u>V</u>irtual <u>E</u>nvironment repository MassIVE (https://massive.ucsd.edu/ProteoSAFe/static/massive.jsp) under accession numbers MSV000082295 (environmental LC-MS/MS), MSV000088621 (*Trichoderma* infection), and MSV000083723 (time-course JKS001884 infection). All image and video data from *Trichoderma* infections and extract behavioral bioassays can be found at https://doi.org/10.7910/DVN/MAYHMQ. Code and additional metadata files used for ITS2 amplicon sequencing analysis and extract tests weeding quantification is available at https://github.com/kek12e/ms_peptaibol.

## Acknowledgments

Funding for this work was received from NSF grants IOS-1656475 (to J.L.K. and M.J.B.) and IOS-1656481 (to P.C.D.). A.M.C.-R. and P.C.D. were supported by the Gordon and Betty Moore Foundation through grant GBMF7622, the U.S. National Institutes of Health for the Center (P41 GM103484 and R01 GM107550) and Federal sub-award DE-SC0021340. The isolation of peptaibols from strains MSX70741 and MSX57715 were funded, in part, by the National Cancer Institute under grant P01 CA125066 (to N.H.O.). J.J.J.v.d.H. is grateful for funding from the Netherlands eScience Center (NLeSC) with an Accelerating Scientific Discoveries Grant (ASDI.2017.030). We thank Dr. Jeremy Balsbaugh and Dr. Jennifer Liddle from the UConn Proteomics and Metabolomics Facility for assistance in acquiring LC-MS/MS data, Dr. Kendra Maas from the UConn Microbial Analysis, Resources and Services (MARS) facility for assistance with ITS amplicon library preparation and sequencing, and Dr. Kim Diver for creating the map of ant colony collection sites. Further thanks are due to Evan Fox and Mariam Zedan for assistance with extractions and to members of the Klassen lab and state park and forest staff for assistance with fieldwork.

## Notes

**Competing Interest Statement:** K.E.K., S.P.P., A.M.C.R., J.R.-C., R.M.S., C.E.E., M.E., M.E.A., J.L.K, and M.J.B declare no competing interest. N.H.O., C.J.P. and H.A.R are members of the Scientific Advisory Board of Clue Genetics, Inc. N.H.O. is also a member of the Scientific Advisory Board of Mycosynthetix, Inc. J.J.J.v.d.H. is a member of the Scientific Advisory Board of NAICONS Srl. P.C.D. is a scientific co-founder and advisor to Ometa and Enveda with prior approval by UC-San Diego and an advisor to Cybele.

### Competing Interest Statement

K.E.K., S.P.P., A.M.C.R., J.R.-C., R.M.S., C.E.E., M.E., M.E.A., J.L.K, and M.J.B declare no competing interest. N.H.O., C.J.P. and H.A.R are members of the Scientific Advisory Board of Clue Genetics, Inc. N.H.O. is also a member of the Scientific Advisory Board of Mycosynthetix, Inc. J.J.J.v.d.H. is a member of the Scientific Advisory Board of NAICONS Srl. P.C.D. is a scientific co-founder and advisor to Ometa and Enveda with prior approval by UC-San Diego and an advisor to Cybele.

### Summary of Updates

Revisions to text, figures, and SI.

