## Supplementary material for "*Trachymyrmex septentrionalis* ants promote fungus garden hygiene using *Trichoderma*-derived metabolite cues": SI_revised

[a]Department of Molecular and Cell Biology, University of Connecticut, Storrs, CT, USA; [b]Division of Medicinal Chemistry, Department of Pharmaceutical Sciences, University of Connecticut, Storrs, CT, USA; [c]Collaborative Mass Spectrometry Innovation Center, Skaggs School of Pharmacy and Pharmaceutical Sciences, University of California San Diego, La Jolla, CA, USA; [d]Department of Chemistry and Biochemistry, University of North Carolina at Greensboro, Greensboro, NC, USA; [e]Department of Natural Products, Instituto de Química, Universidad Nacional Autónoma de México, Coyoacán, Mexico City, Mexico; [f]Department of Chemistry, University of Connecticut, Storrs, CT, USA; [g]Mycosynthetix, Inc., Hillsborough, NC, USA; [h]Section for Clinical Mass Spectrometry, Danish Center for Neonatal Screening, Department of Congenital Disorders, Statens Serum Institut, Copenhagen, Denmark, [i]Bioinformatics Group, Wageningen University, Wageningen, the Netherlands, [j]Department of Biochemistry, University of Johannesburg, Auckland Park, Johannesburg, South Africa, [k]Department of Microbiology and Immunology, University of Michigan, Ann Arbor, MI USA, [l]Department of Medicinal Chemistry, University of Michigan, Ann Arbor, MI, USA.

[1]K.E.K. and S.P.P. contributed equally to this work.

**This PDF file includes:**

Figures S1 to S41
Tables S1 to S8
Legend for Movie S1
Legends for Dataset S1 to S2

**Other supplementary materials for this manuscript include the following:**

Movie S1
Datasets S1 to S2

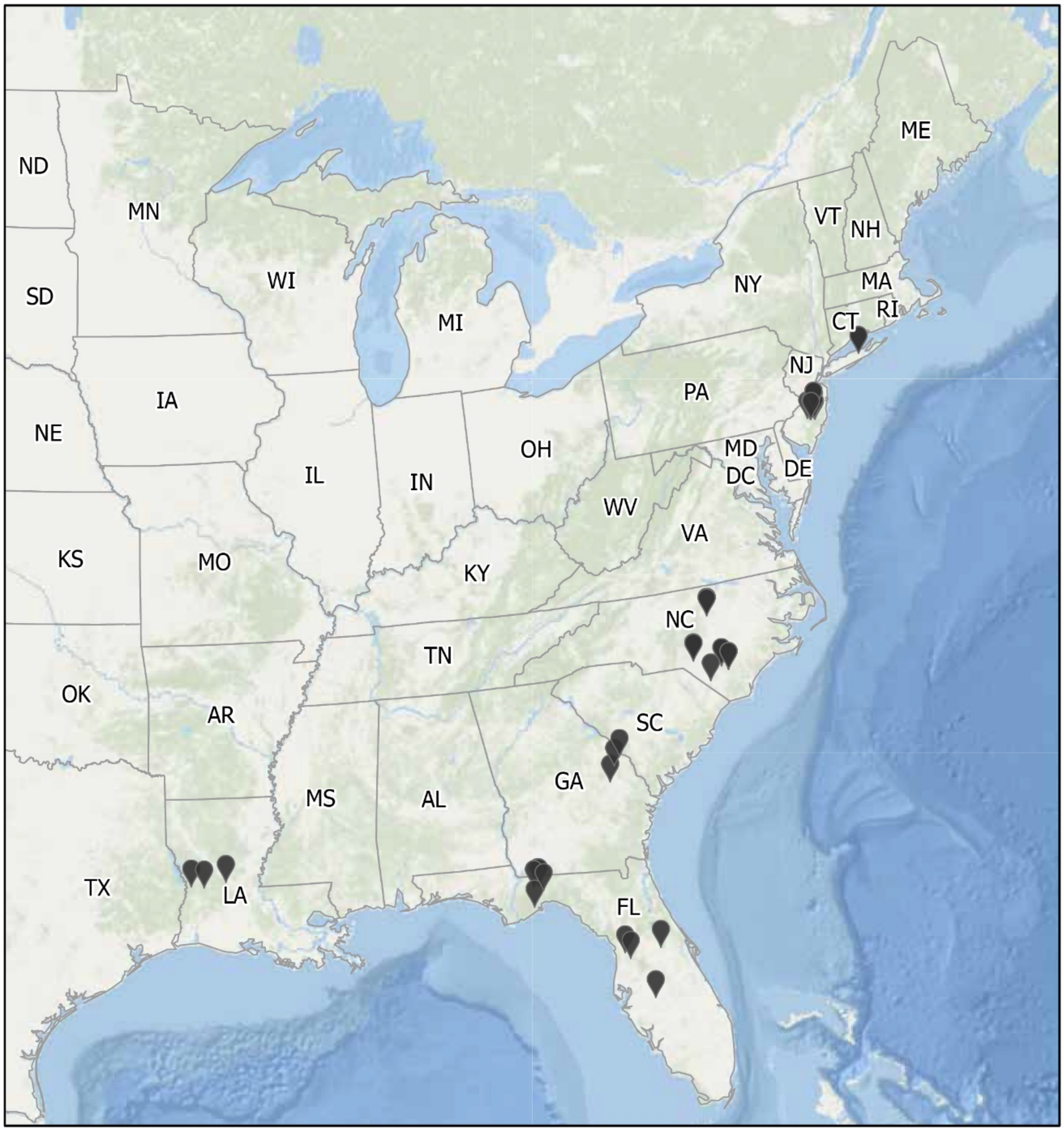

Map credit: Kim Diver, 2022
Data sources: Klassen Lab @ UConn (ant collection sites); Esri, GEBCO, DeLorme, NaturalVue (basemap)

**Fig. S1.** Locations of *T. septentrionalis* field-collections. GPS coordinates were mapped for the state forests and parks in the eastern United States from which the *T. septentrionalis* ant colonies used in this paper were collected. A total of 37 sites were sampled for this study. Map credit: Kim Diver, 2022; data sources: Klassen Lab GPS data, Esri, GEBCO, DeLorme, NaturalVue (basemap).

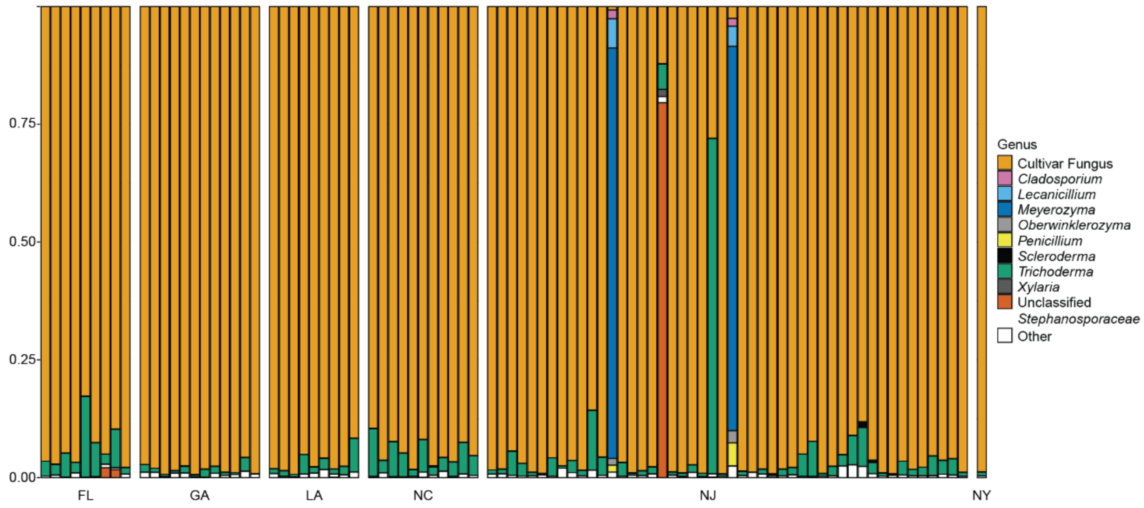

**Fig. S2.** Relative abundance of ITS2 community amplicons in field-sampled *T. septentrionalis* fungus gardens. The relative abundance of fungal taxa is shown for each of 90 field-samples representing 83 fungus gardens from 6 states selected from across *T. septentrionalis's* geographic range. Brackets that connect two stacked bars indicate the 7 colonies that were sampled twice. ASVs grouped as "Other" were < 1% abundant in all samples. All taxa were binned to the genus level except for those from the family *Stephanosporaceae*.

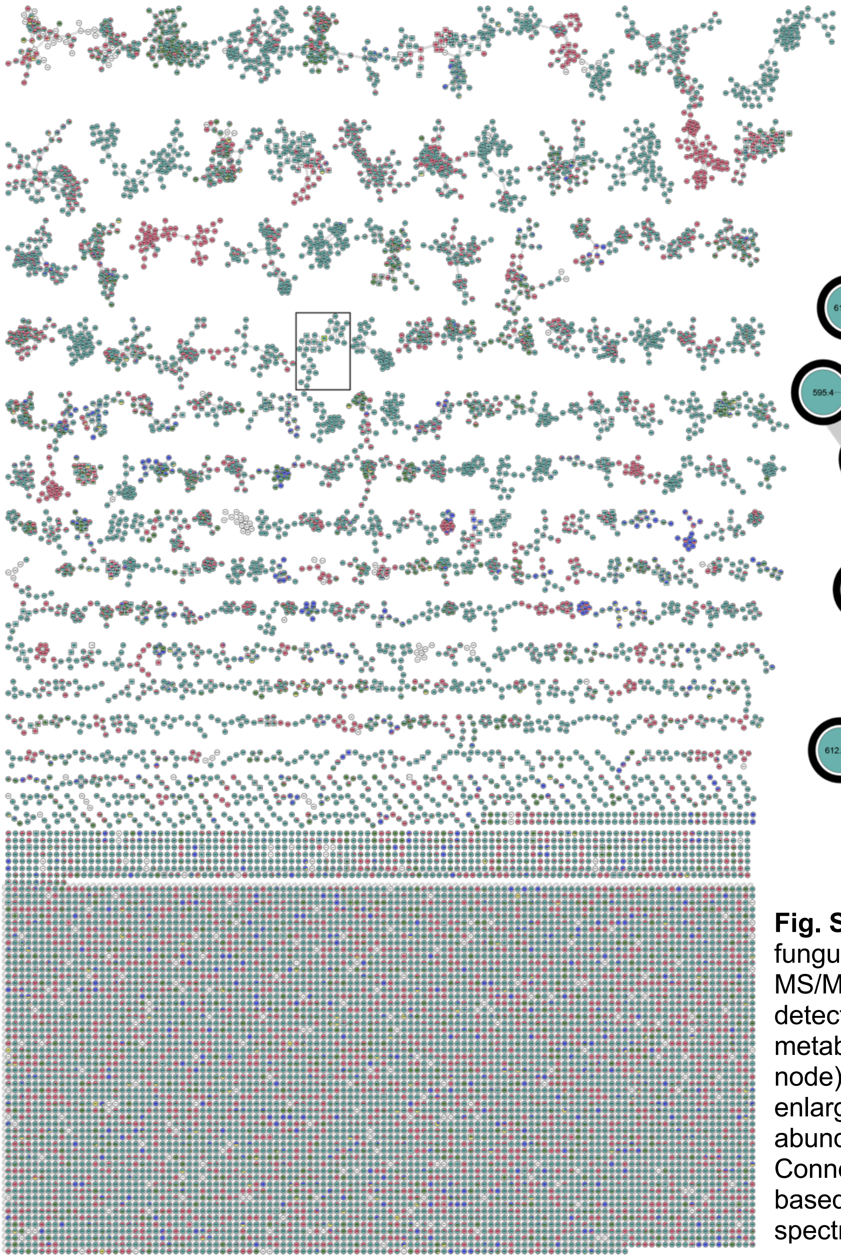

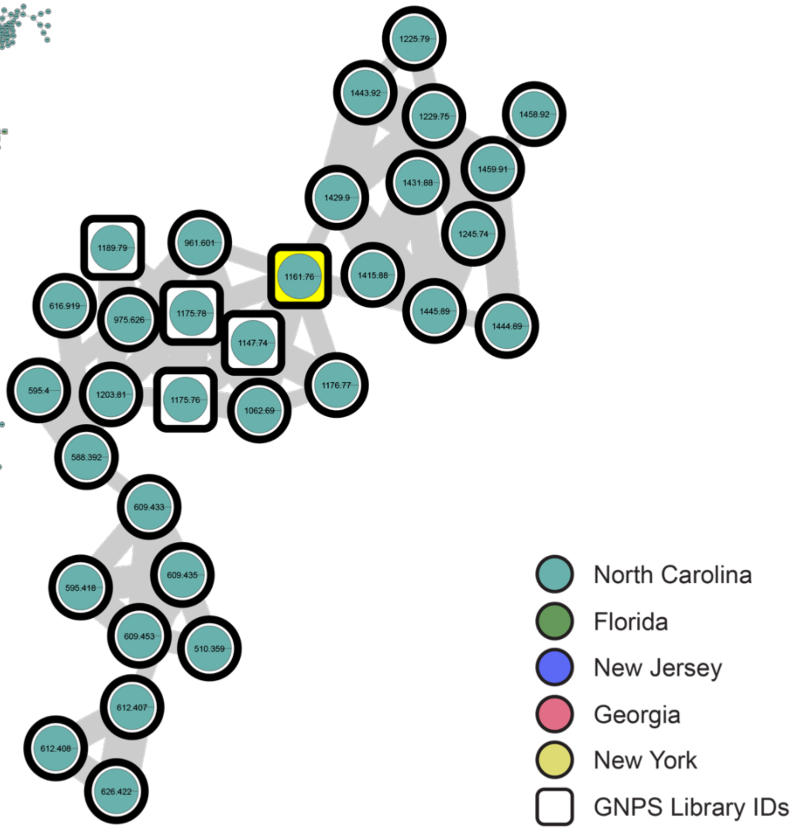

**Fig. S3.** Metabolic diversity in field-sampled *T. septentrionalis* fungus gardens from across their geographic range. Untargeted LC-MS/MS and molecular networking depict metabolite families detected in environmental samples of fungus gardens. A family of metabolites related to peptaibol trichodermide D (yellow square node) is marked by a black rectangle in the full network (left) and enlarged (right). Nodes are pie charts colored by the feature relative abundance for each US state in which they were detected. Connected nodes correspond to structurally related molecules based on their spectral similarity and square nodes represent spectral matches in the GNPS public libraries.

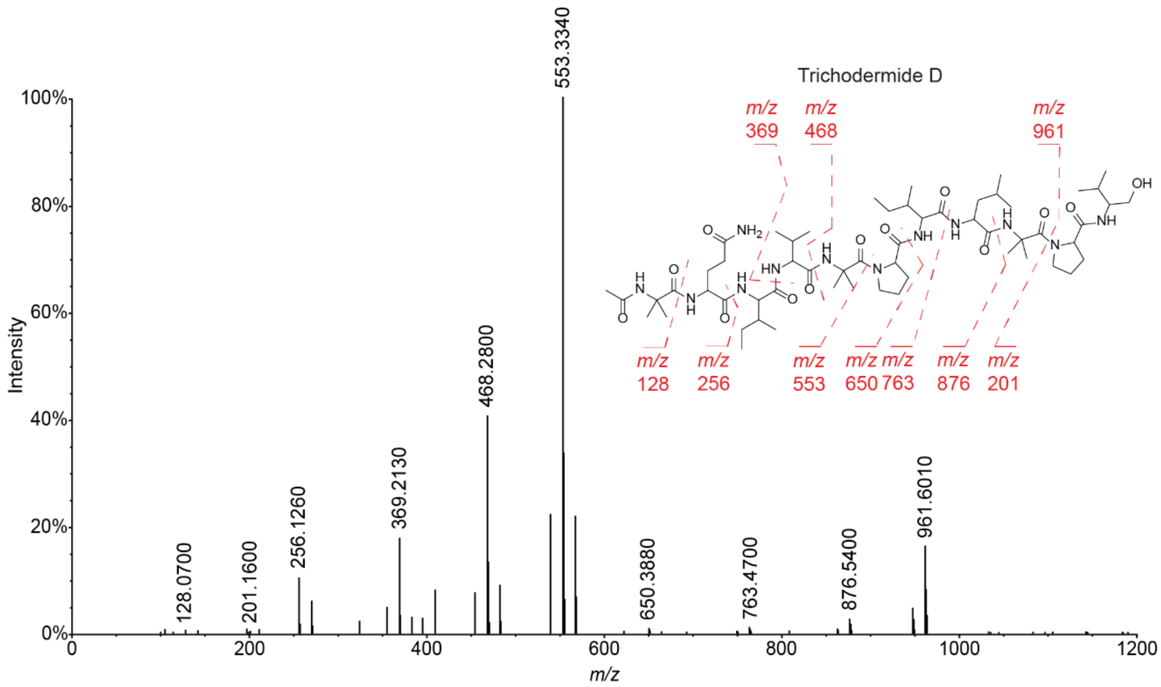

**Fig. S4.** Fragmentation pattern of a representative peptaibol with a mass spectrum consistent with trichodermide D (highlighted in yellow in the peptaibols' molecular family in Figs. 1*B* and S3).

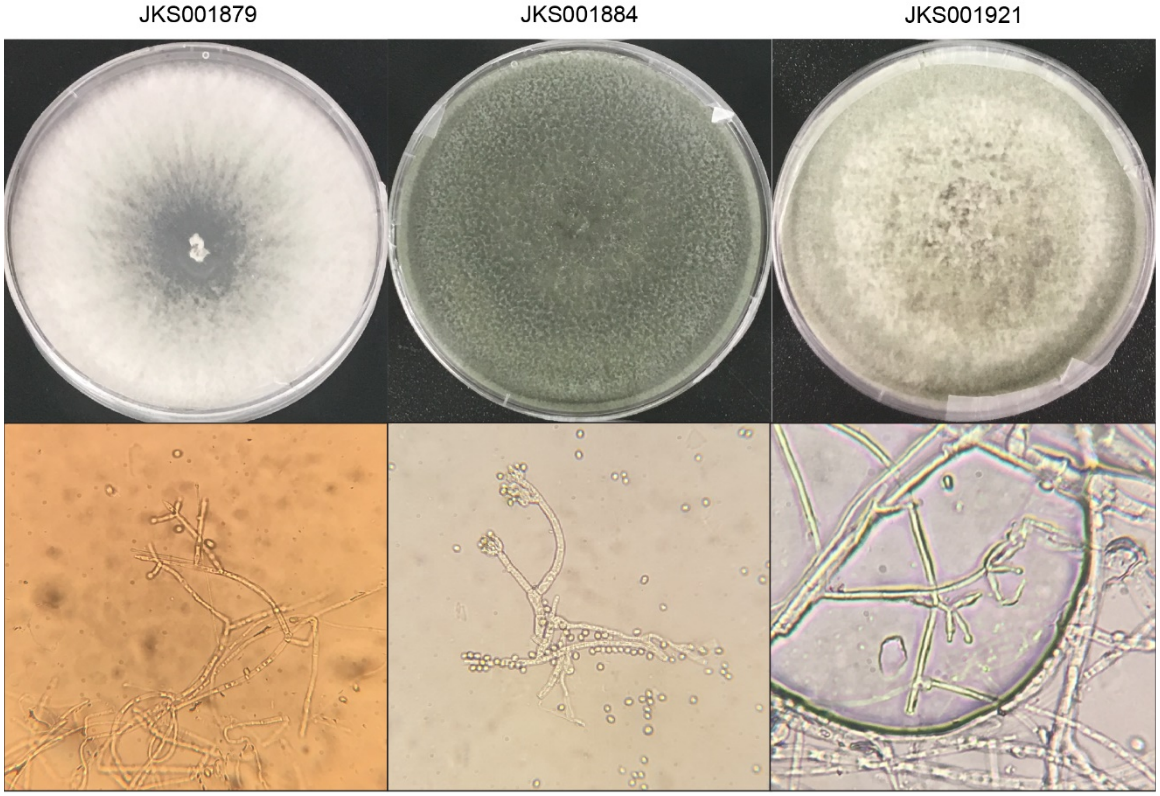

**Fig. S5.** Morphology of *Trichoderma* isolates JKS001879, JKS001884, and JKS001921, classified as *T. koningiopsis*, *T. virens*, and *T. simmonsii*, respectively. Top: Cultures grown for 7 days on potato dextrose agar + antibiotics (see methods). Bottom: Unstained conidiophores for each strain visualized using 1000x light microscopy.

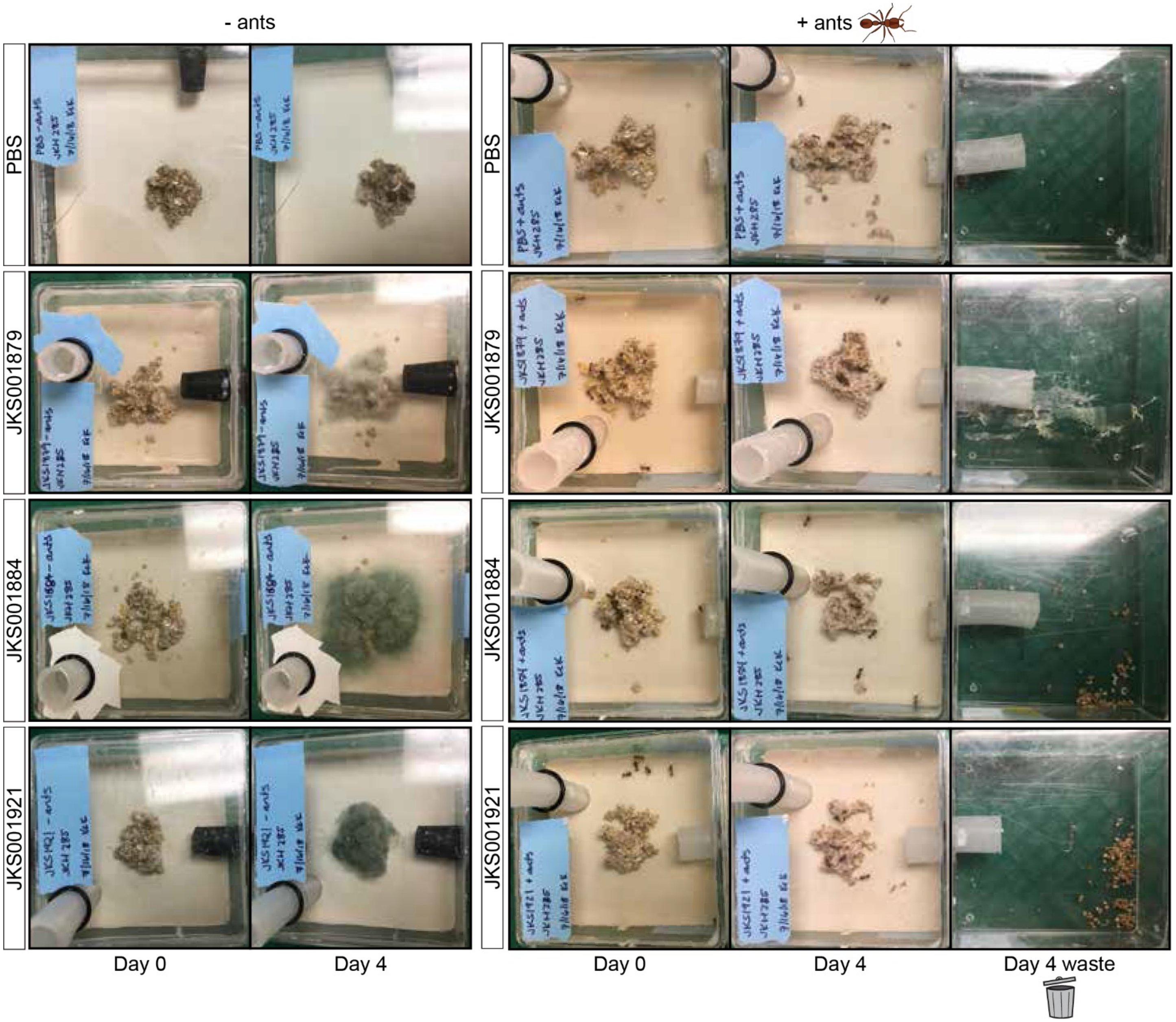
- ants
+ ants
PBS
JKS001879
JKS001884
JKS001921
Day 0
Day 4
Day 4 waste

**Fig. S6.** *Trichoderma* infection images. Images of the connected boxes where ants dispose of waste removed from the fungus garden show waste production for all treatments at Day 4 post-treatment. Treatments using *Trichoderma* strains JKS001884 and JKS001921 had more visible waste than those using *Trichoderma* JKS001879 or PBS. Waste boxes were empty on Day 0 (not shown). These images are of the same experiment as shown in Fig. 2*A* but with timepoint Day 0 and Day 4 waste images included.

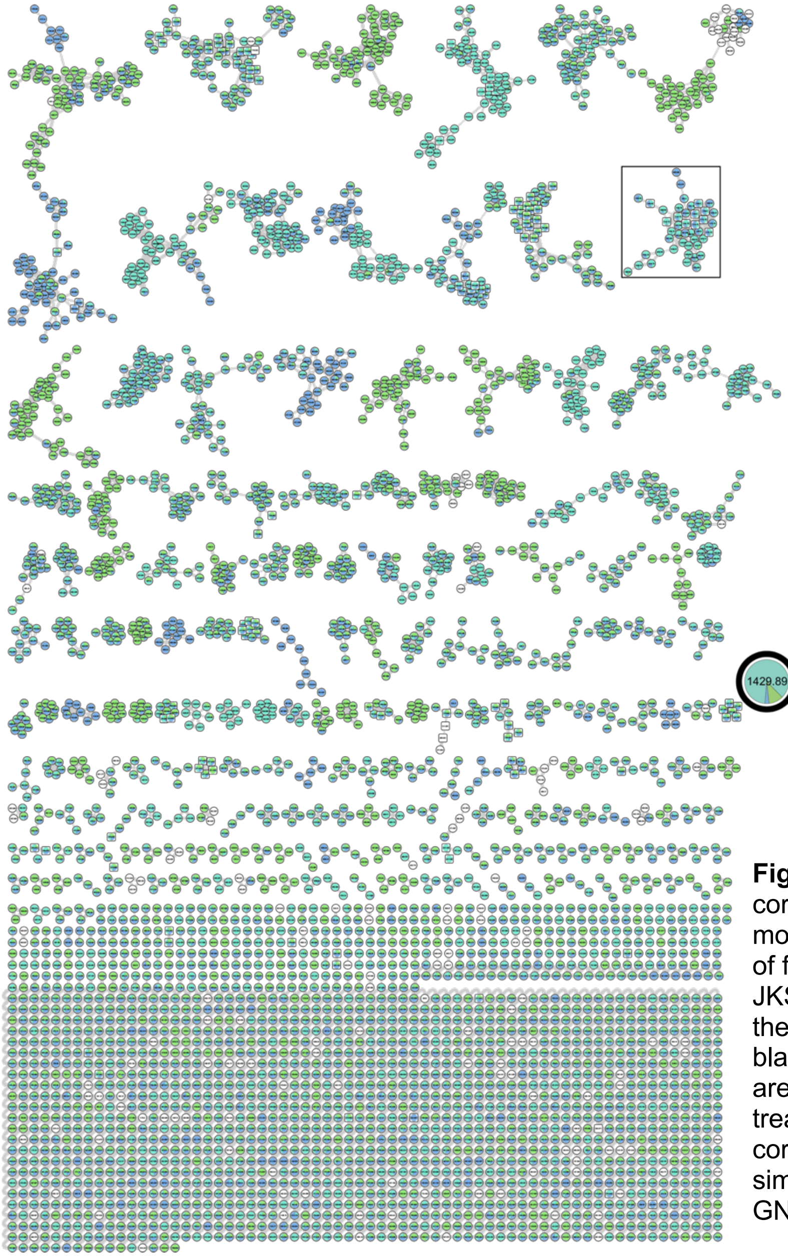

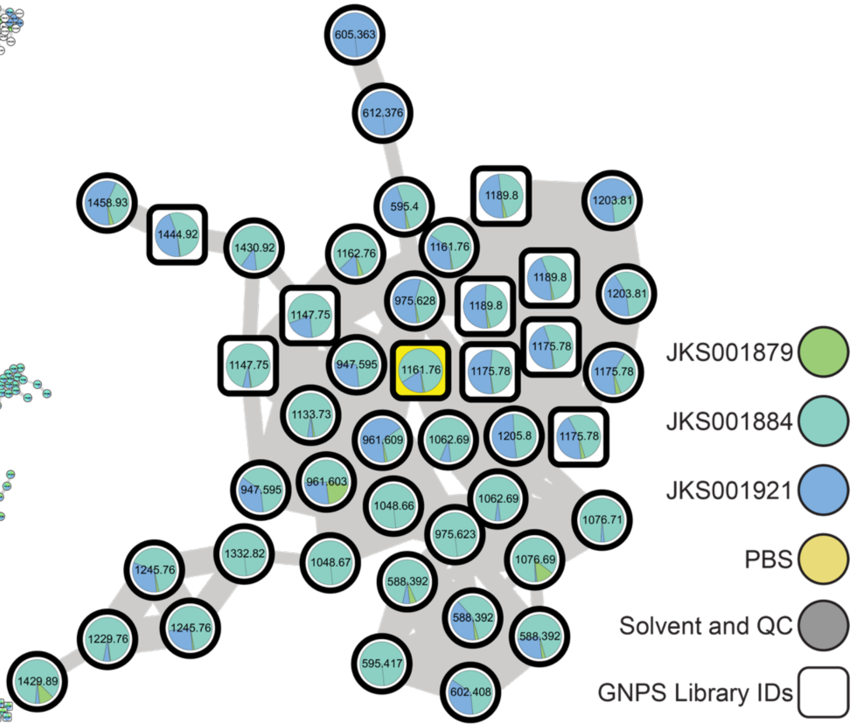

**Fig. S7.** Metabolic diversity in the *Trichoderma* infection experiment corresponding to Fig. 2*A* and S6. Untargeted LC-MS/MS and molecular networking depict metabolite families detected in samples of fungus gardens treated with *Trichoderma* sp. JKS001879, JKS001884, JKS001921, or PBS. A family of metabolites related to the peptaibol trichodermide D (yellow square node) is marked by a black rectangle in the full network (left) and enlarged (right). Nodes are pie charts colored by the feature relative abundance for each treatment in which they were detected. Connected nodes correspond to structurally related molecules based on their spectral similarity and square nodes represent spectral matches to the GNPS public libraries.

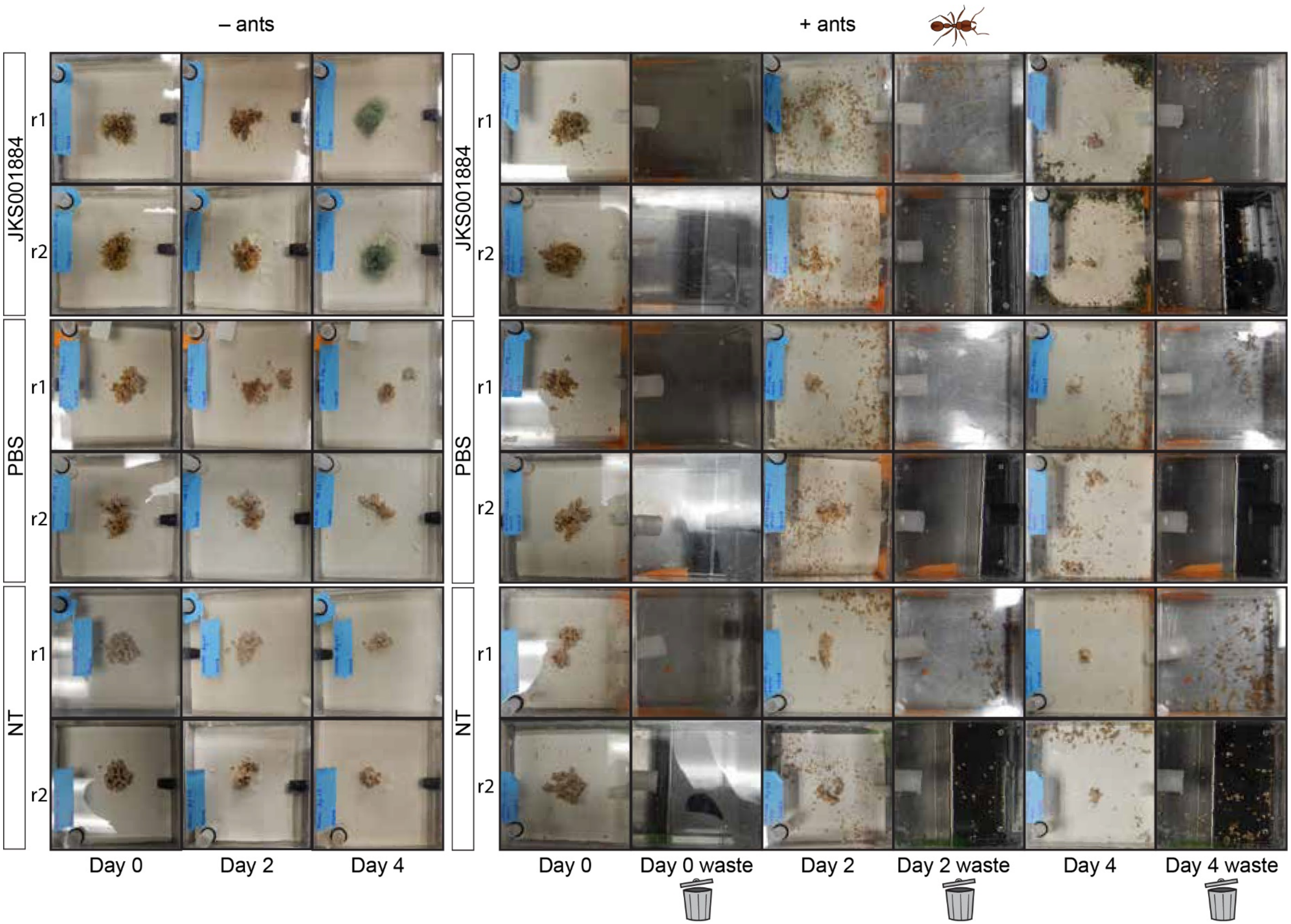
– ants
+ ants
JKS001884
PBS
NT
r1
r2
Day 0
Day 2
Day 4
Day 0 waste
Day 2 waste
Day 4 waste

**Fig. S8.** Time-course *Trichoderma* sp. JKS001884 infection experiment images showing the subcolonies subsampled for Fig. 2*B*-*C*. In addition to the expected weeding in response to *Trichoderma* treatment, ants also weeded their fungus gardens in the negative controls likely due to the disturbance of sampling ants and fungus garden for molecular analyses every other day (see Fig. 2*B*-*C*, S9-S11). However, only *Trichoderma*-treated waste became visibly infected by 4 days post-inoculation as seen by the dark green pieces of waste around the perimeter of the main box for those treatments. *Trichoderma* can also be seen as green growth on the fungus gardens in the *Trichoderma*-treated –ants treatments at day 4.

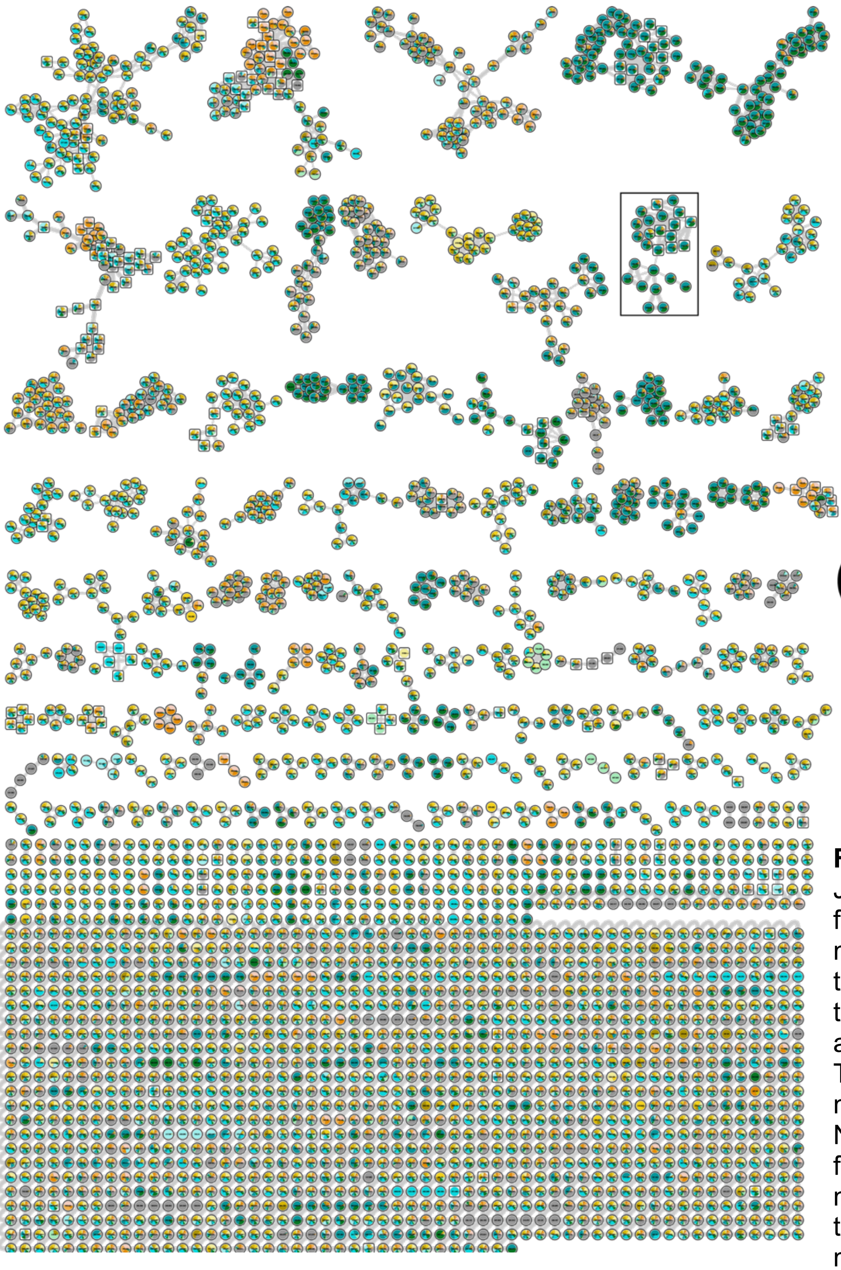

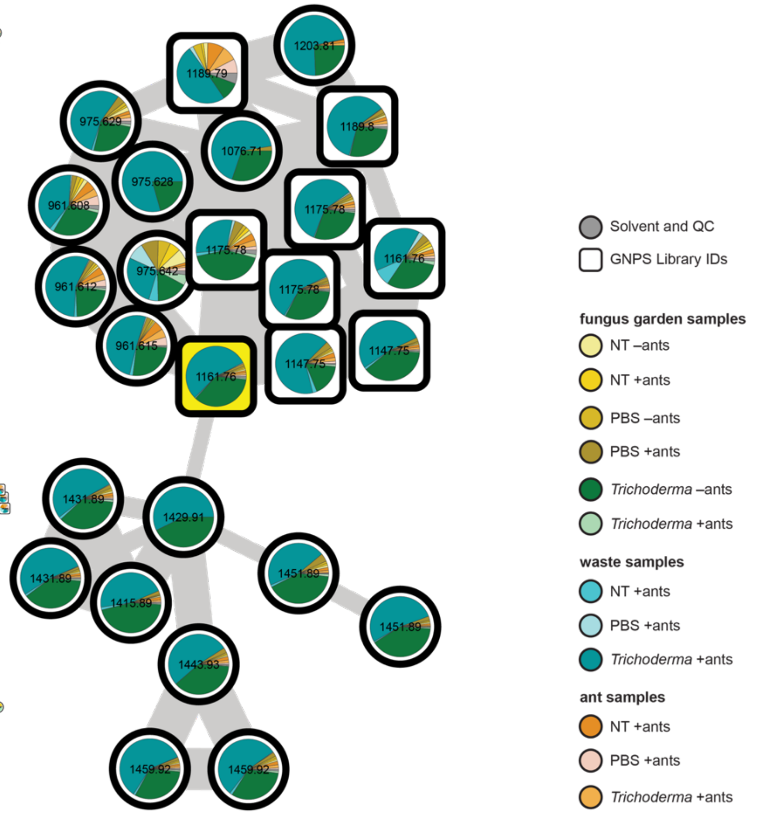

**Fig. S9.** Metabolic diversity in time-course *Trichoderma* sp. JKS001884 infection showing the relative abundance of each feature in each sample type. Untargeted LC-MS/MS and molecular networking depict metabolite families detected in the time-course infection samples. A family of metabolites related to peptaibol trichodermide D (yellow square node) is marked by a black rectangle in the full network (left) and enlarged (right). The enlarged peptaibol family here is the same as the top network in Fig. 2*C* but with the full legend of colors included. Nodes are pie charts colored by the feature relative abundance for each sample type in which they were detected. Connected nodes correspond to structurally related molecules based on their spectral similarity and square nodes represent spectral matches in the GNPS public libraries.

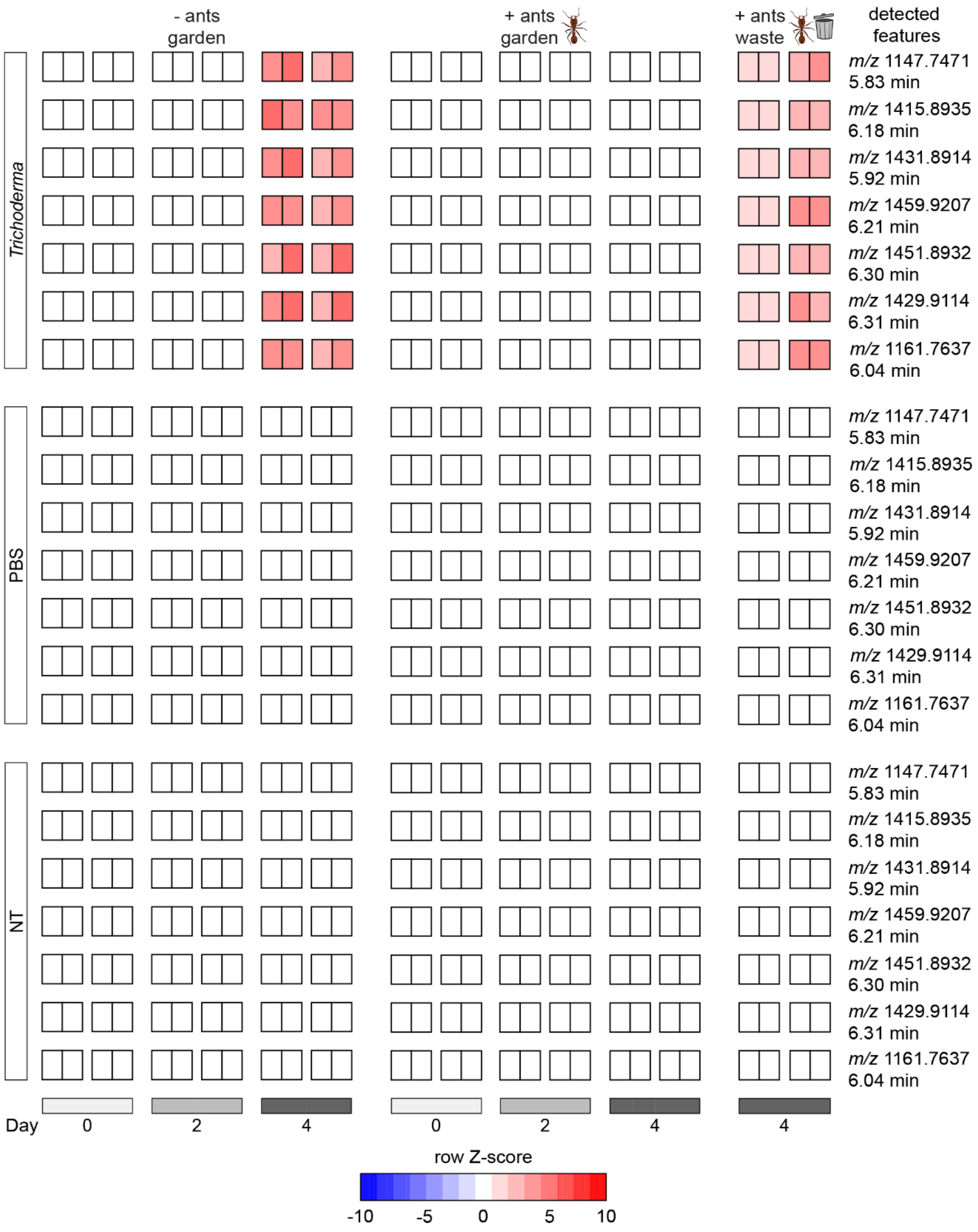

**Fig. S10.** Heatmap of peptaibol-related metabolites detected in samples from the time-course *Trichoderma* sp. JKS001884 infection. For each timepoint on the bottom x-axis there were two biological replicates (r1 and r2, see Fig. S8) and two technical replicates shown as squares colored by metabolite abundance using row Z-score. Technical replicates shown with no spacing between squares while biological replicates shown spaced slightly apart. The three treatment conditions on the left y-axis and individual rows correspond to 1 of the 7 detected peptaibol-like features listed on the right y-axis. The top x-axis separates the samples by treatment (+/- ants) and sample type (garden or waste). Peptaibol-related features were found at higher abundances in samples where we also visibly detected *Trichoderma* infection including the *Trichoderma*-treated fungus gardens without ants and the waste of the *Trichoderma*-treated gardens with ants (Fig. S8). The abundance of these detected features also correlated with the abundance of *Trichoderma* reads detected with ITS2 amplicon sequencing of parallel samples (Fig. 2*B*).

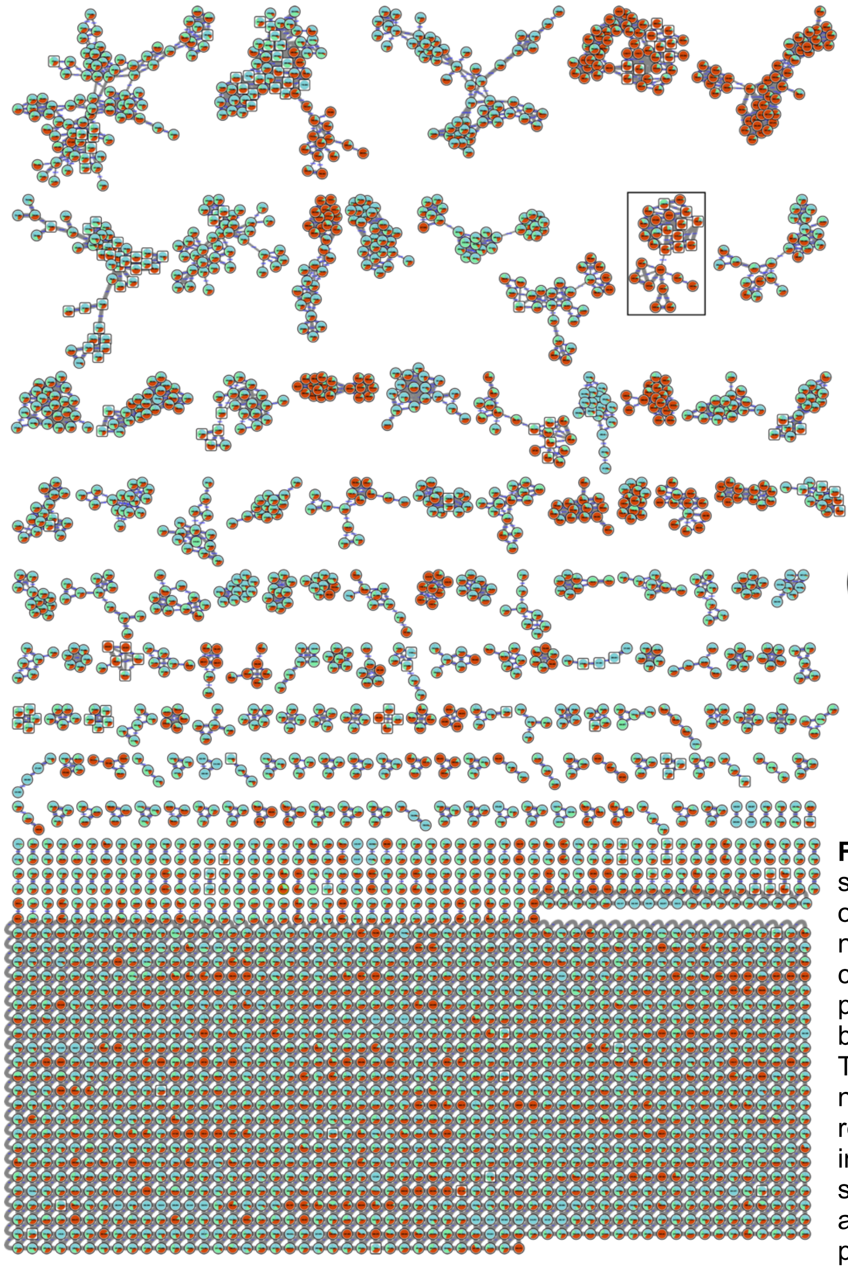

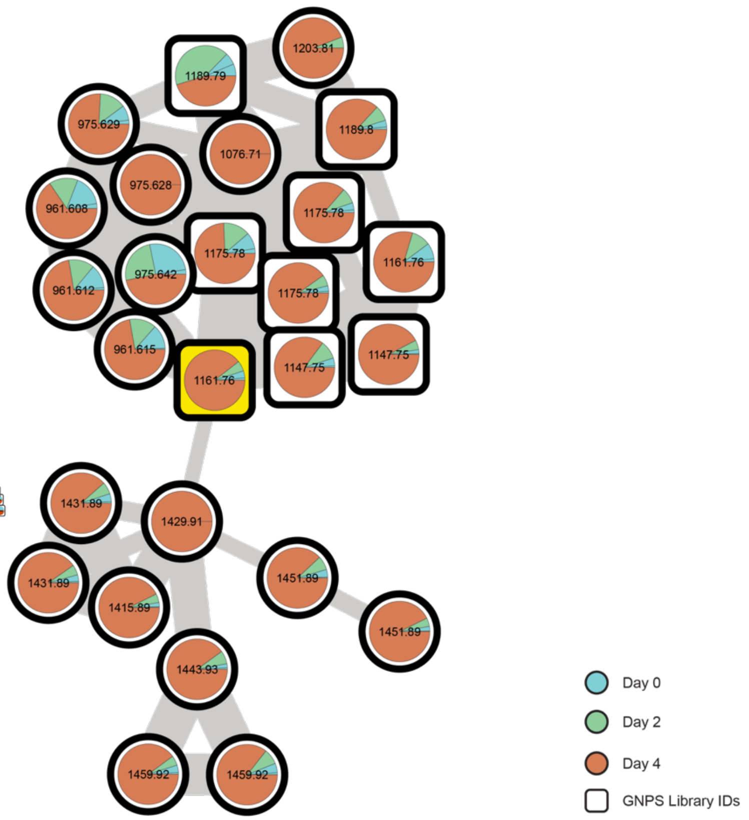

**Figure S11.** Metabolic diversity in time-course *Trichoderma* sp. JKS001884 infection showing feature relative abundance over infection time. Untargeted LC-MS/MS and molecular networking depict metabolite families detected in the time-course infection samples. A family of metabolites related to peptaibol trichodermide D (yellow square node) is marked by a black rectangle in the full network (left) and enlarged (right). The enlarged peptaibol family here is the same as the bottom network in Fig. 2*C*. Nodes are pie charts colored by the feature relative abundance for each day post experimental treatment in which they were detected. Connected nodes correspond to structurally related molecules based on their spectral similarity and square nodes represent spectral matches in the GNPS public libraries.

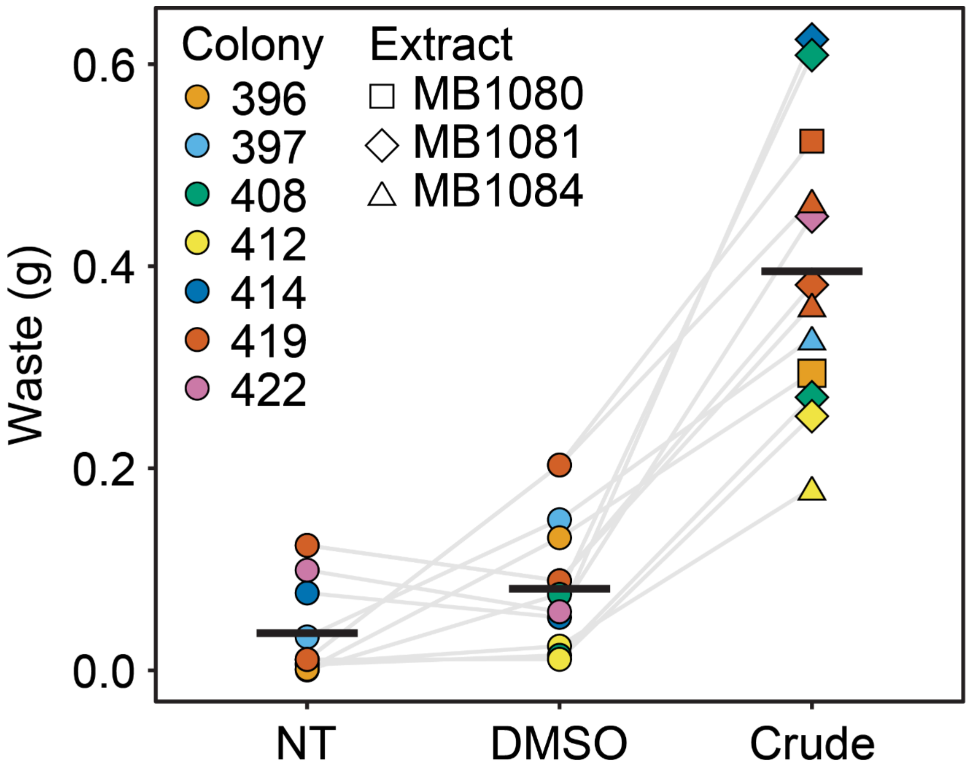

**Fig. S12.** Mass of waste produced by ants in response to crude *Trichoderma* extracts, showing the different extract IDs for the three independent batches of crude *Trichoderma* extract used for this study. The fungus gardens in *T. septentrionalis* subcolonies were treated with 10 mg/mL of crude extract in 0.5% DMSO, 0.5% DMSO, or were not treated, and waste produced by ants was collected and weighed after 24 h. Colors indicate unique ant colony IDs, and values from biologically independent trials are connected. This graph is identical to Fig. 3*A*, but includes the different extract IDs for the crude treatment. No obvious variability was seen in ant weeding response to different crude extracts.

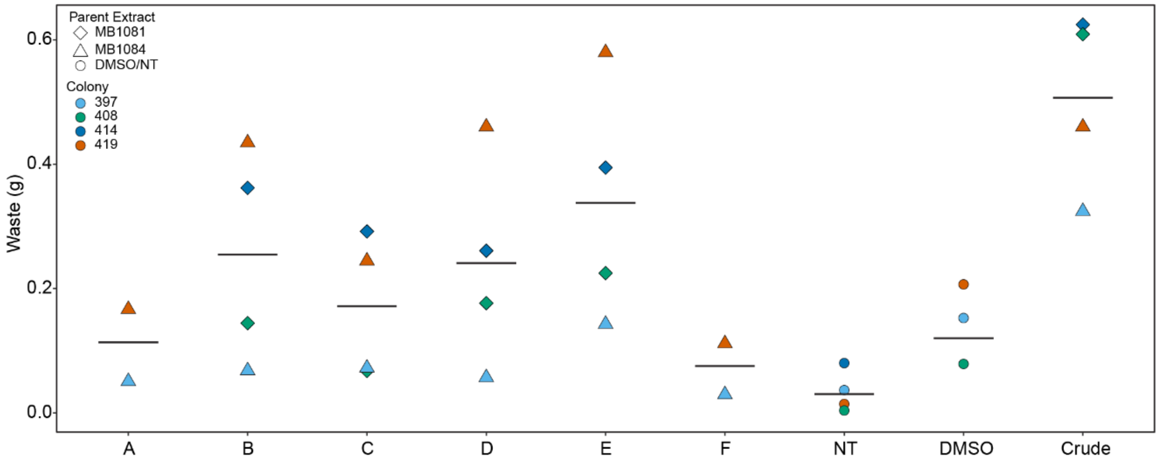

**Fig. S13.** Ant waste production in response to different extracts (Fig. 3*B*). Two independent crude extracts were fractionated (see methods) and tested for *T. septentrionalis* weeding response. Fractions were tested at 1 mg/mL and crude extracts were tested at 10 mg/mL, each dissolved in 0.5% DMSO. Waste produced by the ants was collected and weighed after 24 h. Independent ant colonies shown using color and the parent extract of the fractions shown using shapes. No obvious variability was seen in ant weeding response to fractions originating from different crude extracts.

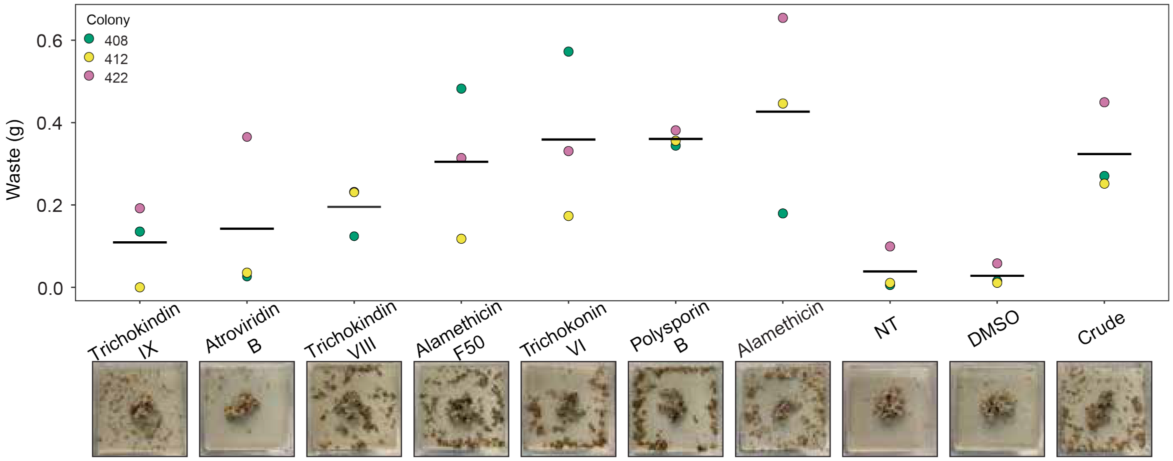

**Fig. S14.** Pure peptaibols induce ant weeding. Fungus gardens in *T. septentrionalis* subcolonies were treated with 0.25 mg/mL of each pure peptaibol dissolved in 0.5% DMSO, and waste created by ant weeding was weighed after 24 h. Images represent the weeding induced by each compound after 24 h, where the remaining fungus garden is in the center of the box and the waste is deposited by the ants away from this center fungus garden as they weed. All compounds elicited a stronger average weeding response than the NT and DMSO negative controls. Alamethicin F50, trichokinin VI, polysporin B, and alamethicin elicited average weeding similar to or greater than that elicited by the crude extract (tested at 10 mg/mL), while new compounds trichokindins VIII (**1**) and IX (**2**) and atroviridin B induced weaker mean weeding responses than the crude extract.

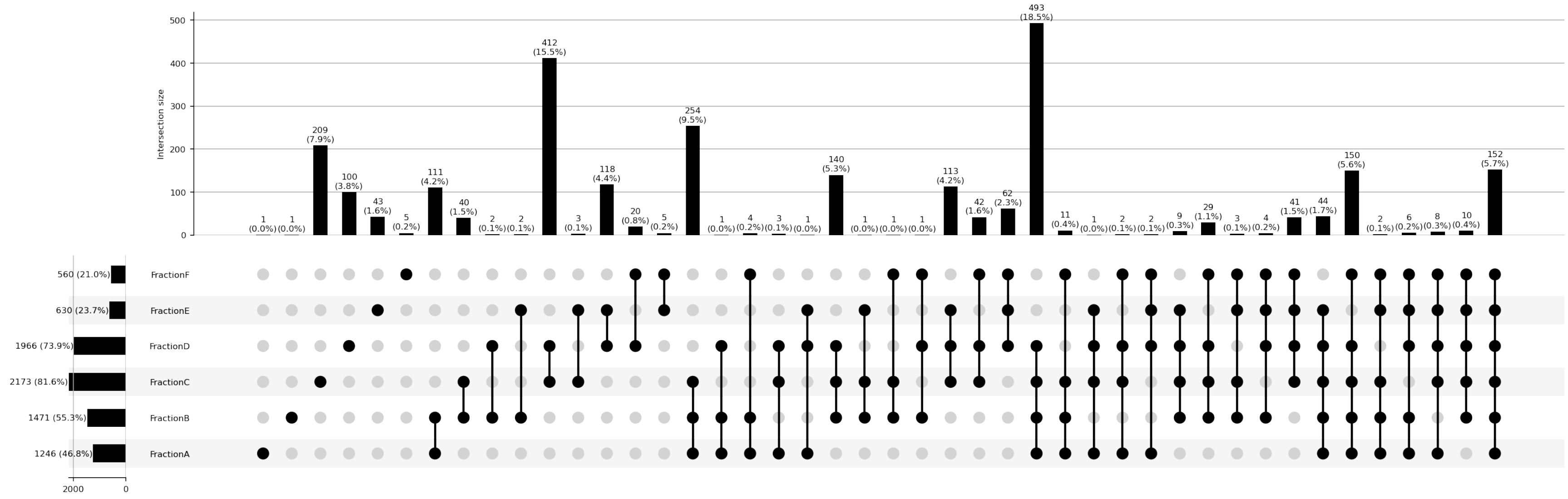

**Fig. S15.** UpSet plot indicating shared features of JKS001884 *Trichoderma* sp. fractions MB1084A-F. The bioactive fractions D and E exclusively shared 118 features, with an additional suite of features found to be highly enriched in fractions D and E but present in lower abundances in other fractions. Although fraction B induced similar waste production as fractions D and E, this fraction was found to be chemically dissimilar (B and D share only two features, B and E also share only two features). Fraction B was highly similar to fraction A (111 features shared), which was one of the least bioactive fractions.

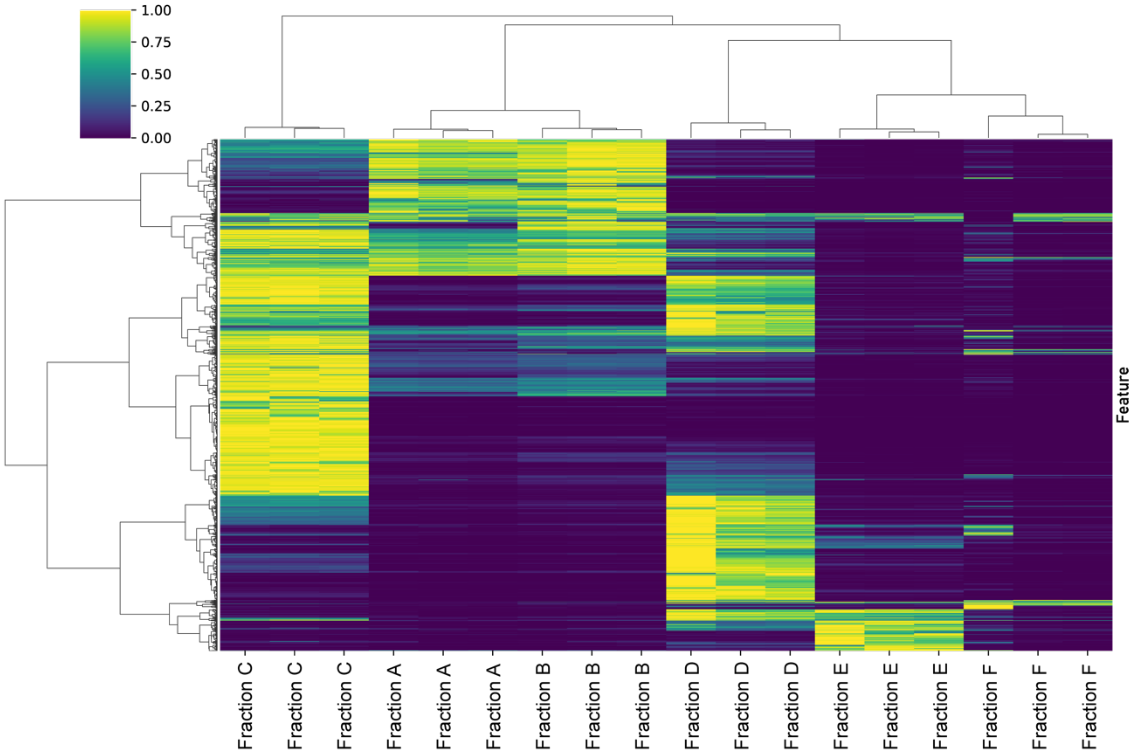

**Fig. S16.** Full heat map of metabolites from JKS001884 *Trichoderma* sp. fractions. The bioactive fractions D and E have several regions of similarity as highlighted in Fig. 4*A*. Although fraction B induced similar waste production as fractions D and E, this fraction has only minimal similarity in feature abundance with these two fractions and demonstrates much more similarity with the inactive fraction A.

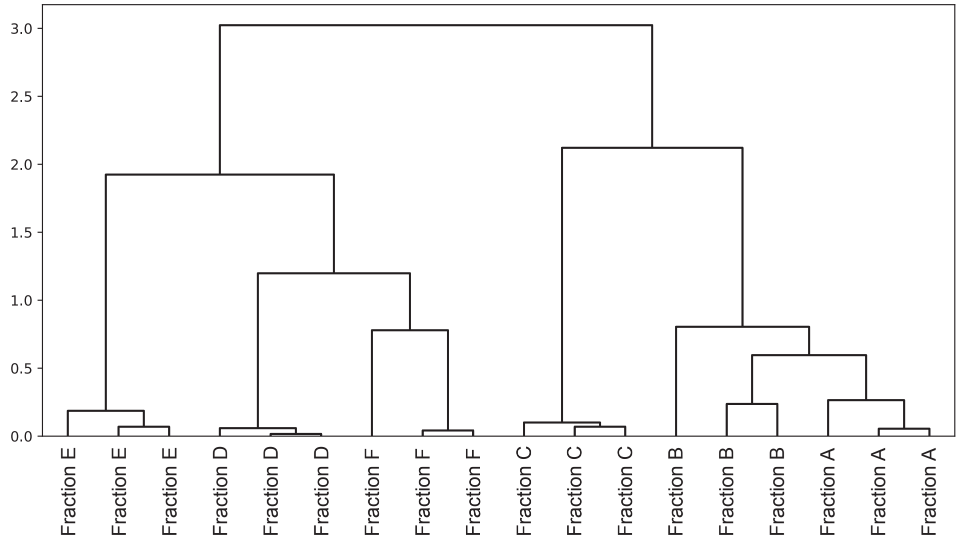

**Fig. S17.** Dendrogram of JKS001884 *Trichoderma* sp. fractions. Note that fractions D-F group separately from fractions A-C.

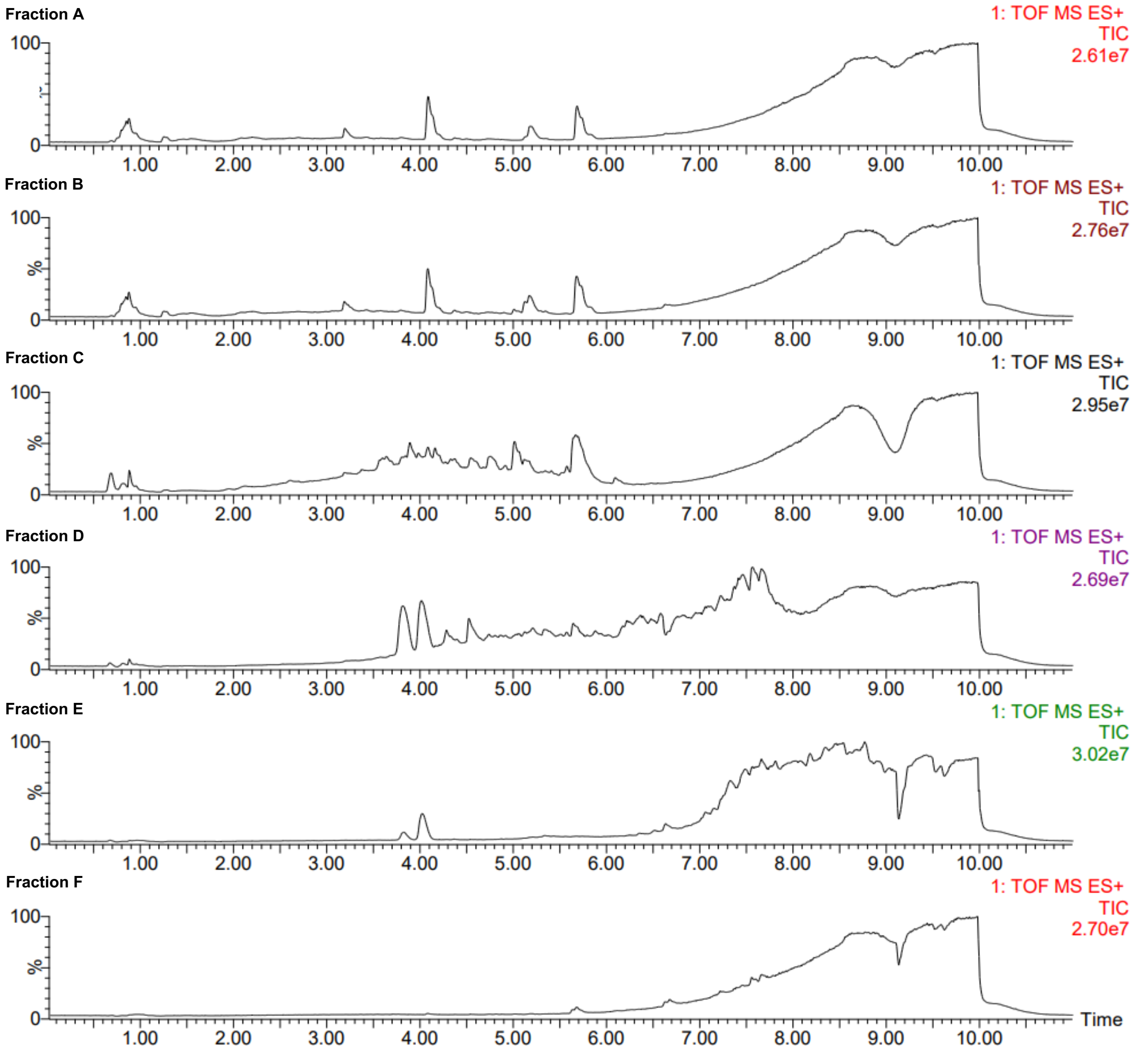

**Fig. S18.** Total ion chromatograms (TICs) of fractions from *Trichoderma* sp. JKS001884 including full runs of each injection. Runs are identical to those shown in Fig. 4*D*. Comparisons of total ion chromatograms indicated considerable overlap of peaks in both fractions D and E.

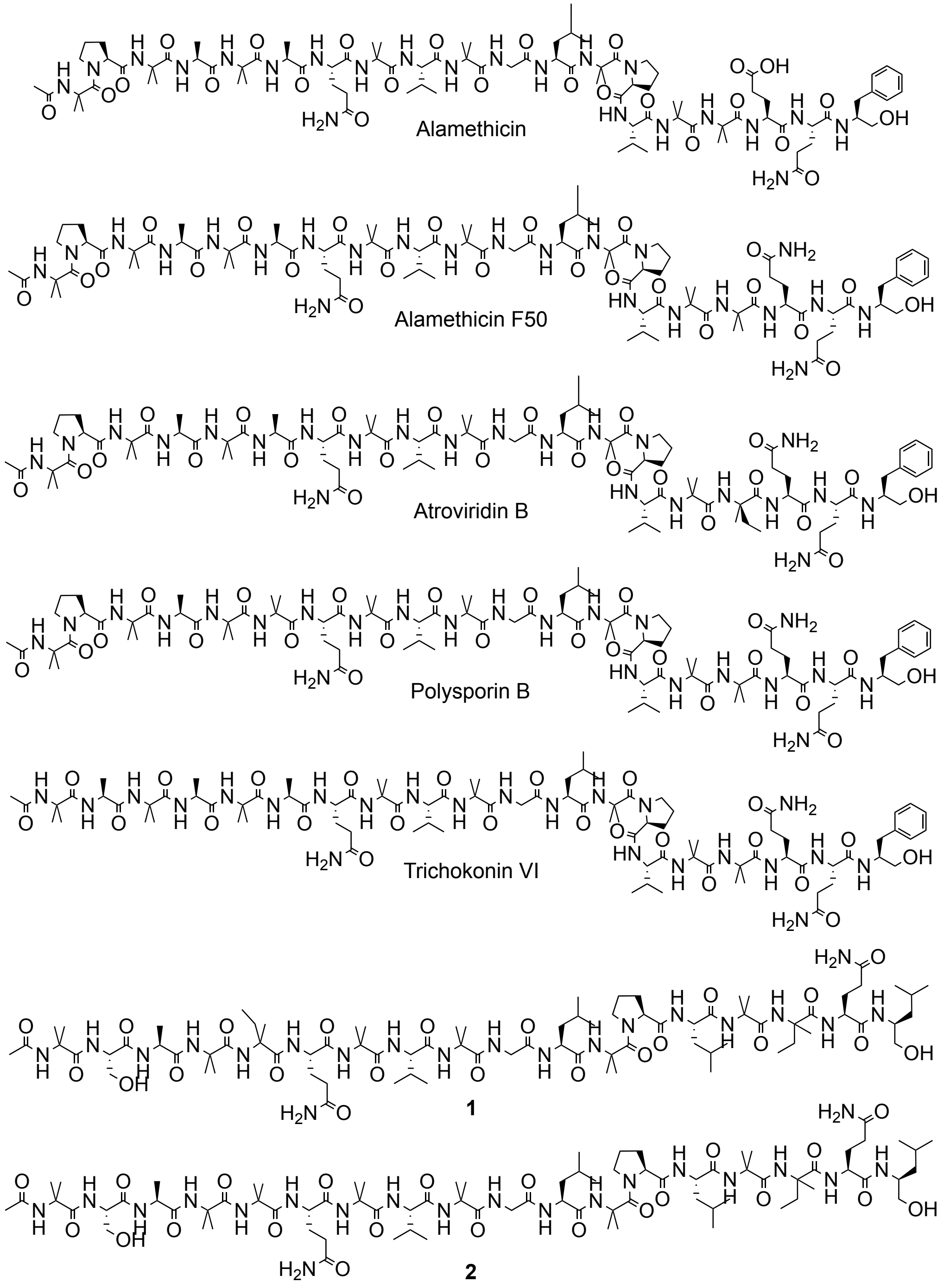

**Fig. S19.** Structures of two new peptaibol metabolites (**1** and **2**) and other pure peptaibols tested for ant behavioral activity.

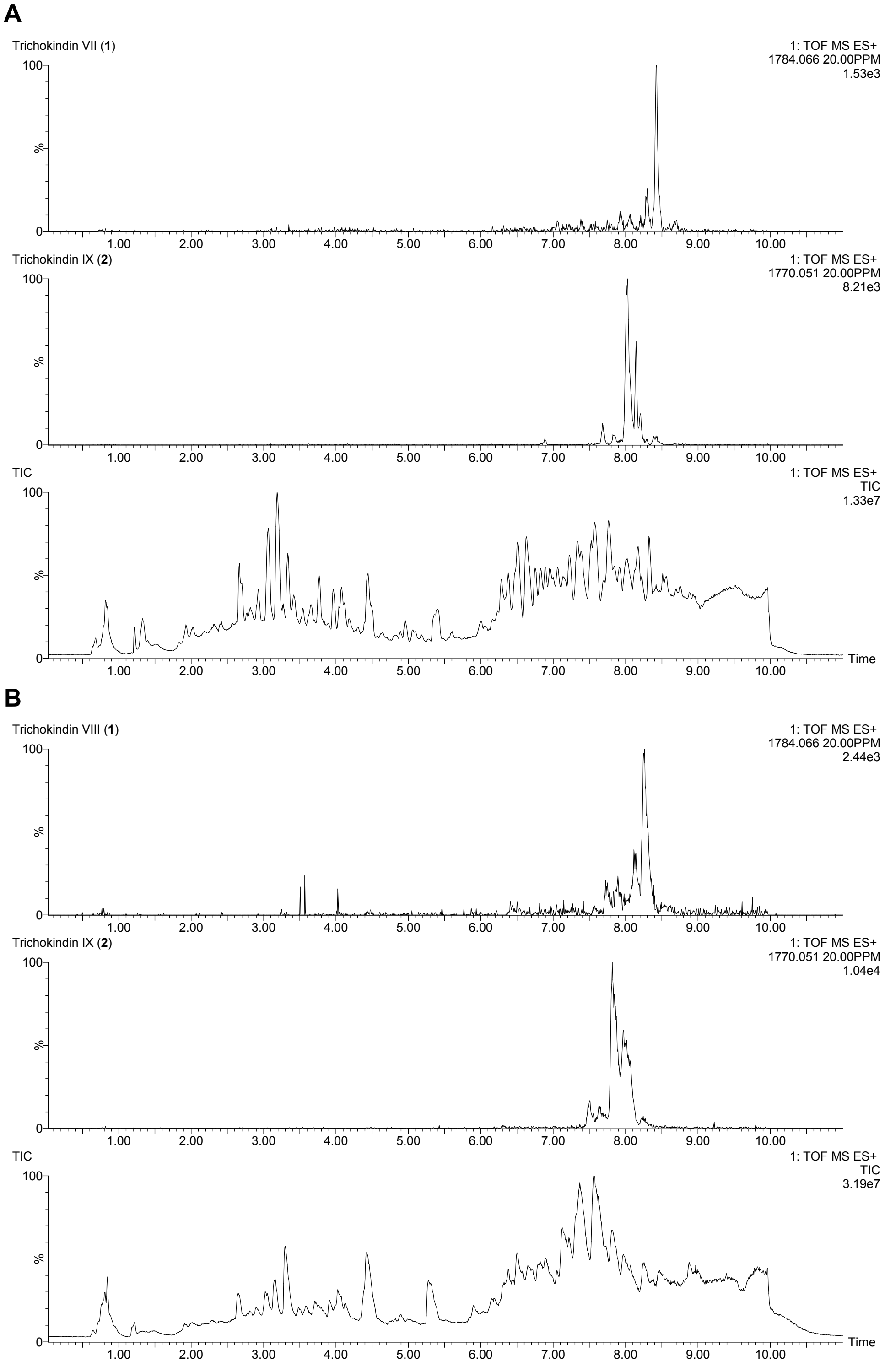

**Fig. S20.** Extracted ion chromatograms (EICs) for new compounds **1** and **2** in crude *Trichoderma* sp. JKS001884 extracts MB1081 (A) and MB1084 (B). Total ion chromatograms (TICs) shown below each EIC.

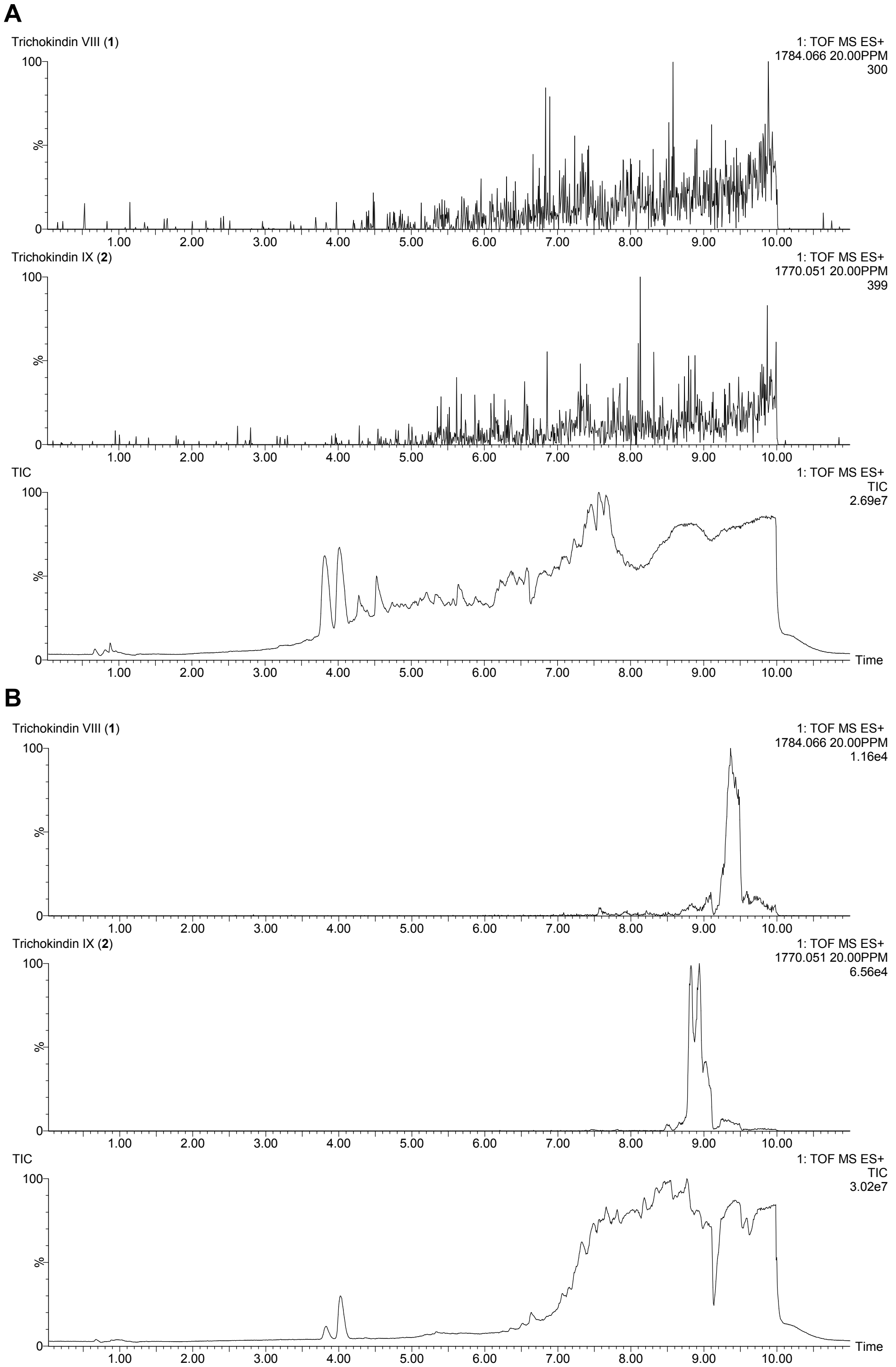

**Fig. S21.** Extracted ion chromatograms (EICs) of new compounds **1** and **2** in fractions D (A) and E (B) from *Trichoderma* sp. JKS001884 extract MB1084. Total ion chromatograms (TICs) shown below each EIC.

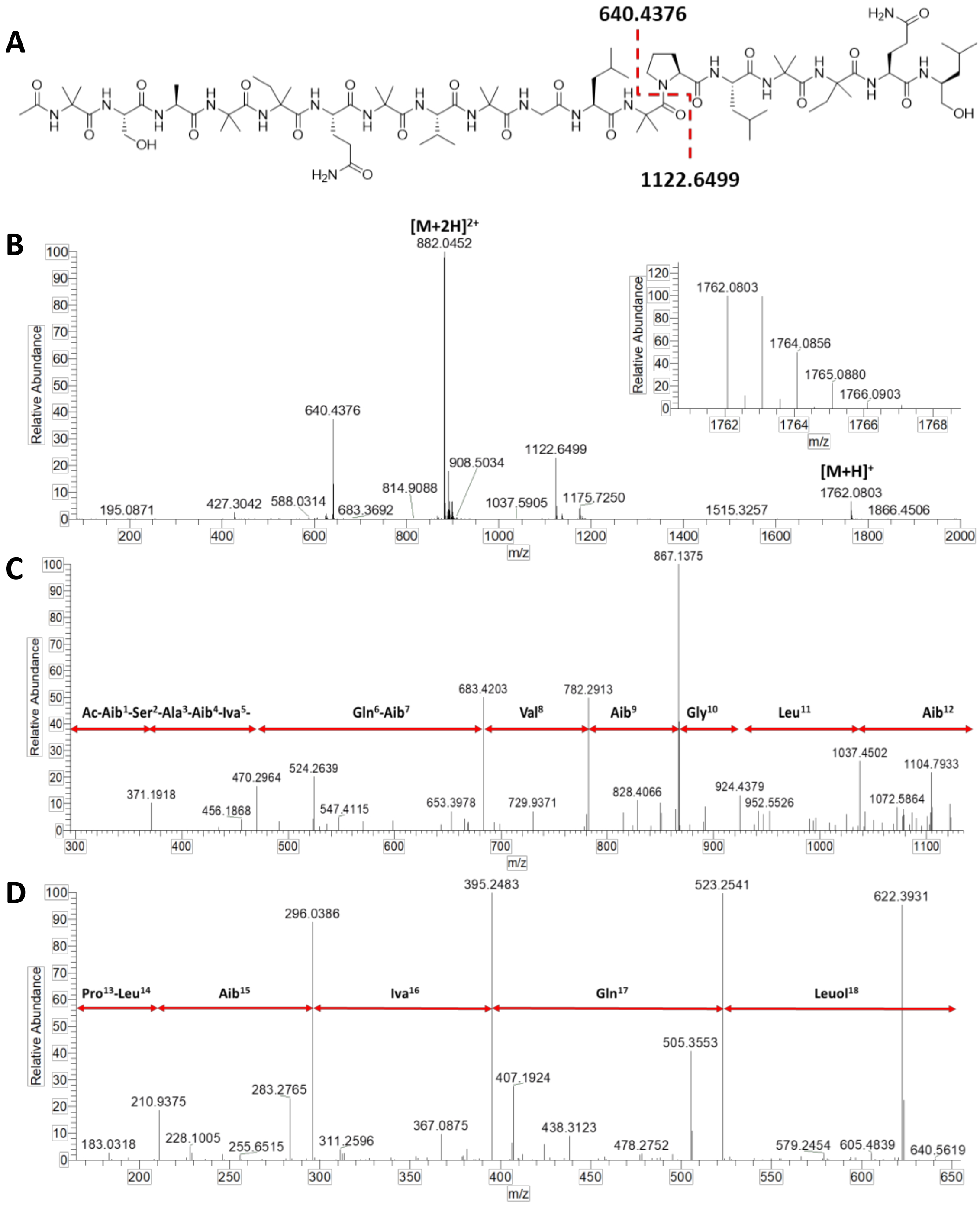

**Fig. S22.** Positive HRESIMS data of compound **1**. A) Structure of **1**. B) Full scan showing in source fragmentation. C and D) HRESIMS$^2$ of fragments $b_{12}^+$ and $y_6^+$, respectively.

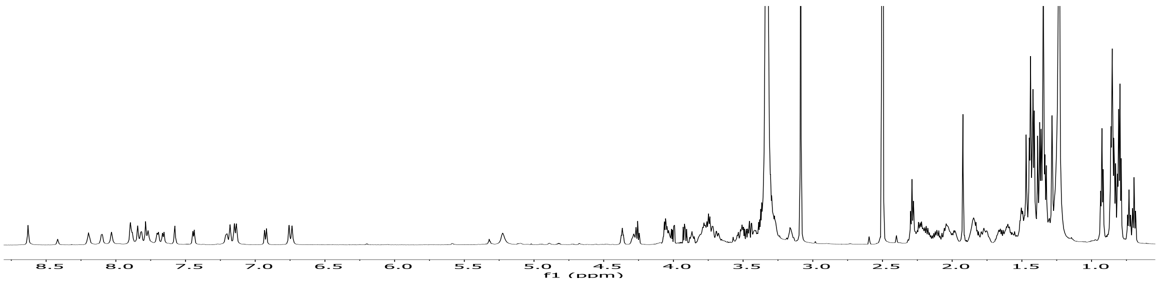

**Fig. S23.** $^{1}$H NMR data for compound **1** (700 MHz in DMSO-$d_6$).

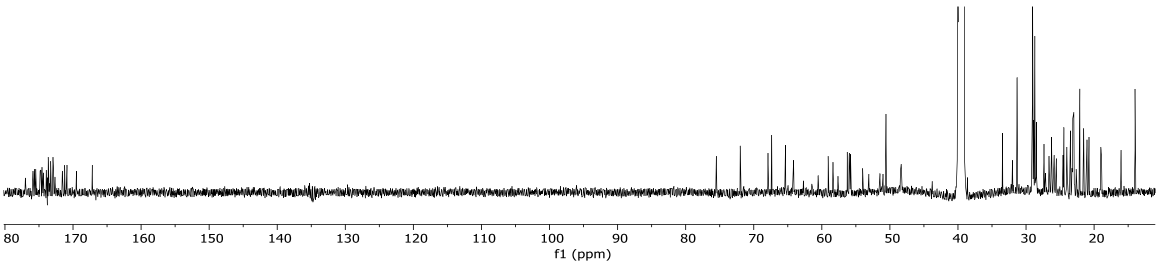

**Fig. S24.** $^{13}$C NMR data for compound **1** (175 MHz in DMSO-$d_6$).

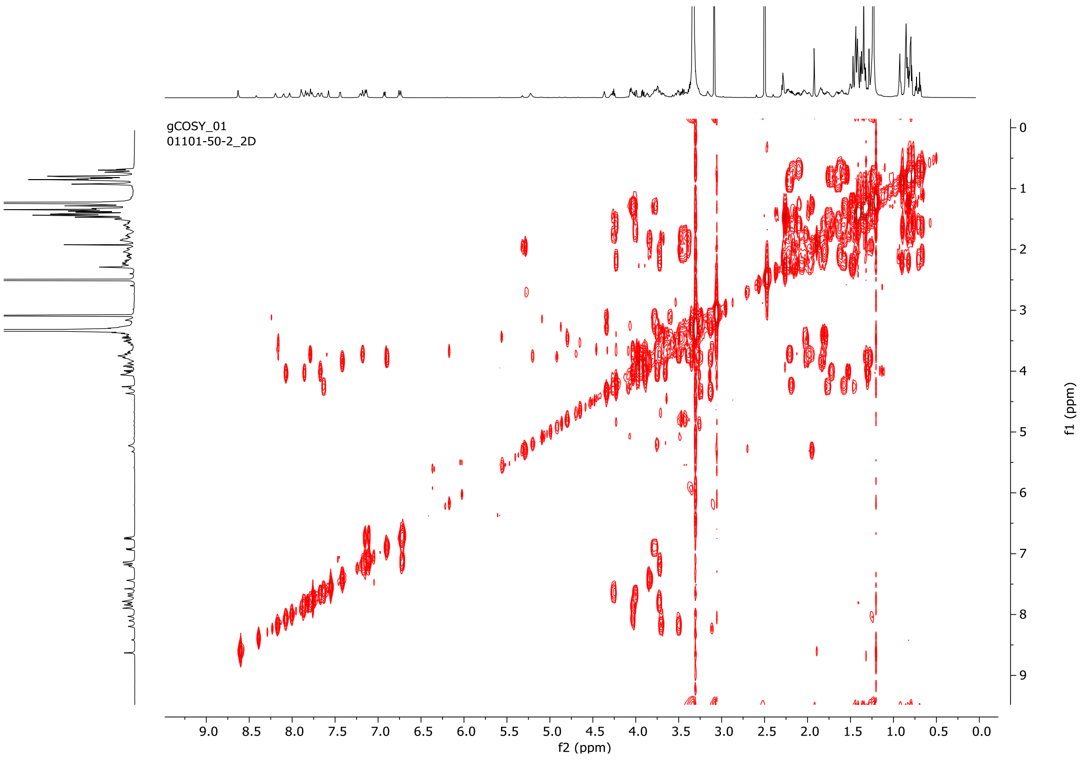

**Fig. S25.** $^{1}H$-$^{1}H$ COSY NMR data for compound **1** (DMSO-$d_6$).

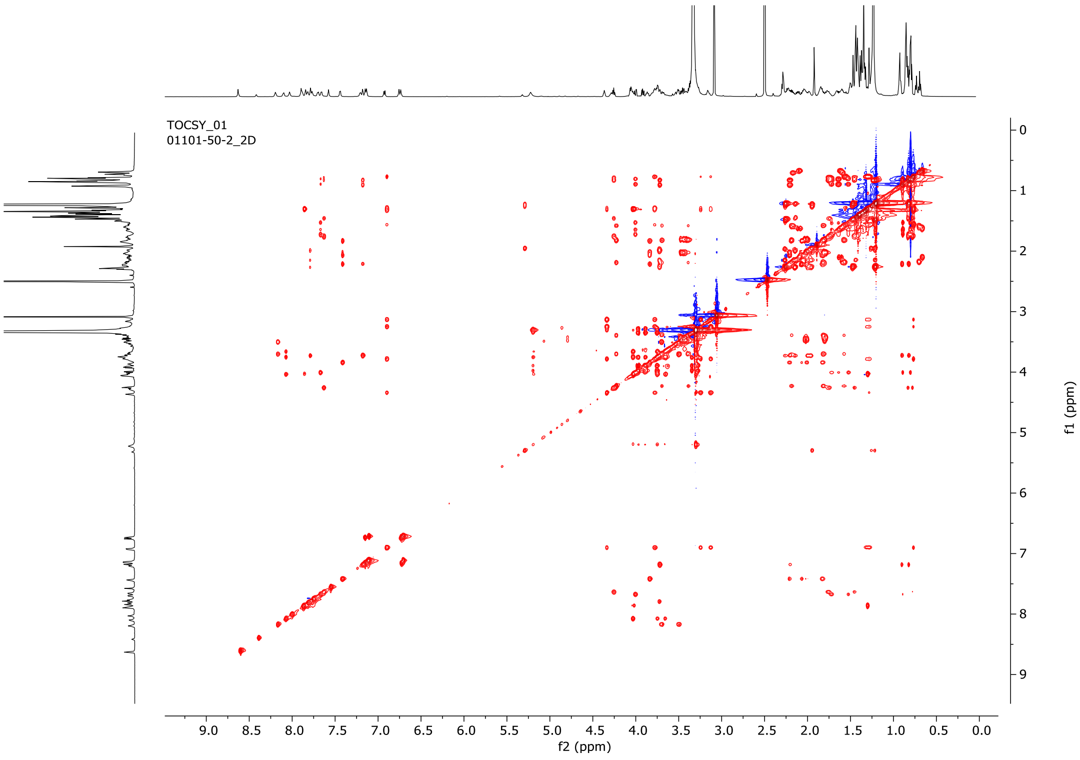

**Fig. S26.** $^{1}H$-$^{1}H$ TOCSY NMR data for compound **1** (DMSO-$d_6$).

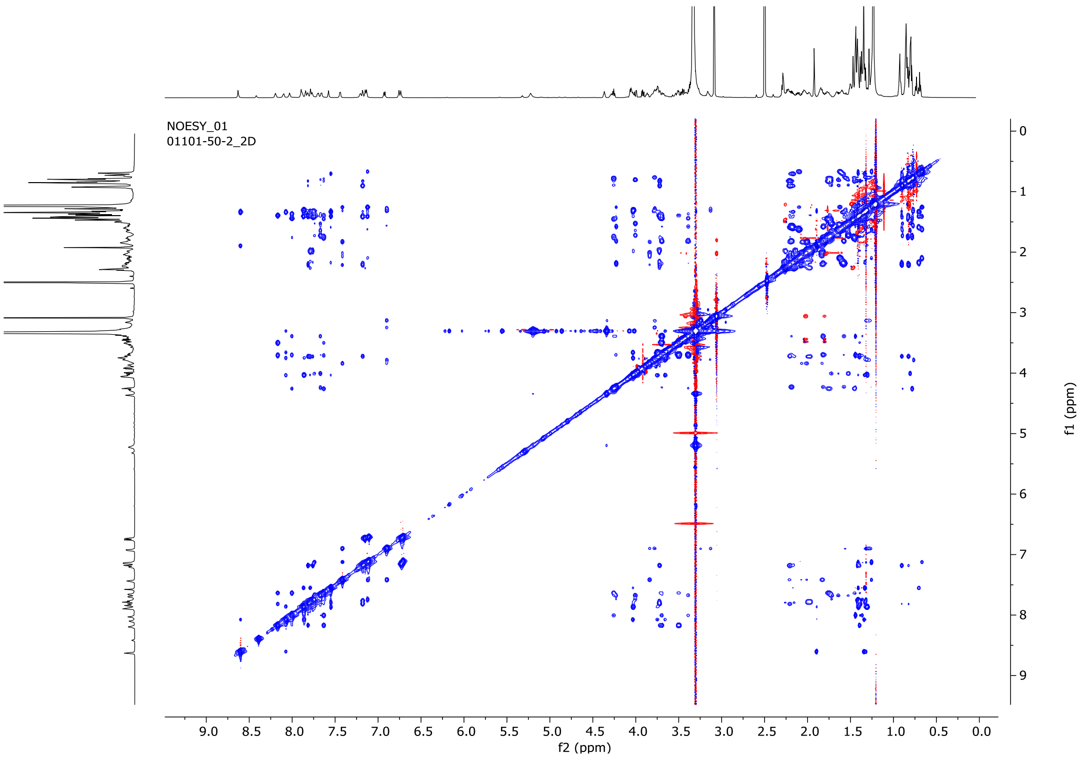

**Fig. S27.** $^{1}$H-$^{1}$H NOESY NMR data for compound **1** (DMSO-$d_6$).

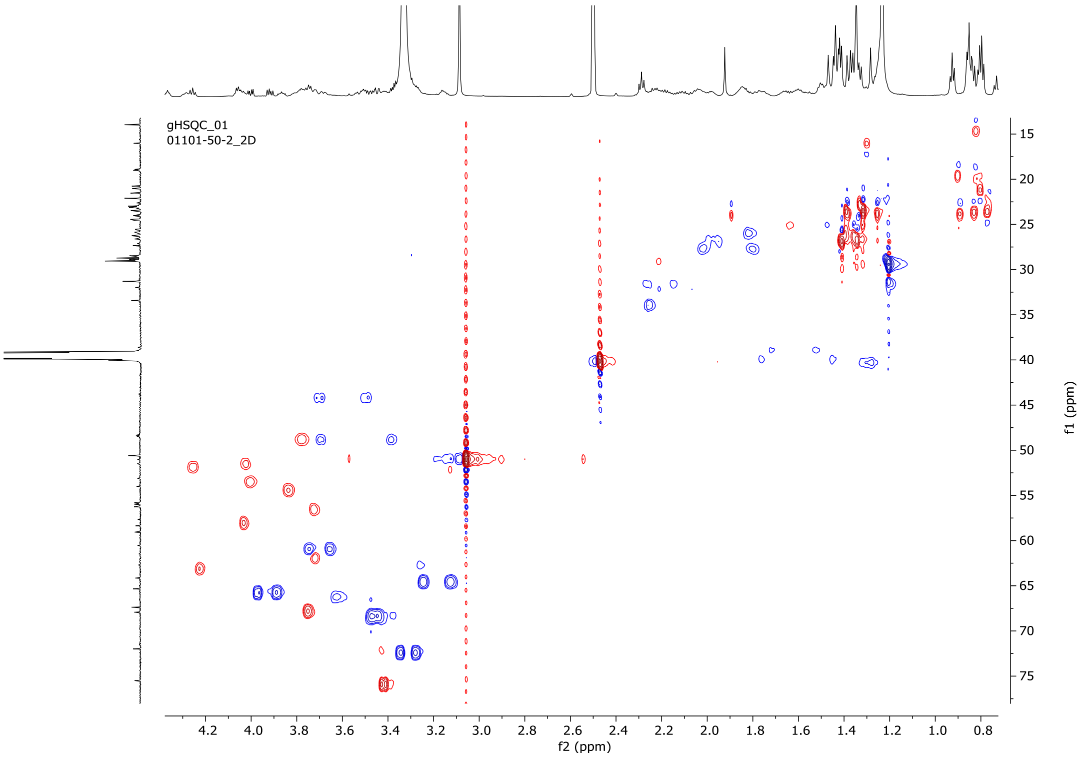

**Fig. S28.** $^{1}$H-$^{13}$C HSQC NMR data for compound **1** (DMSO-$d_6$).

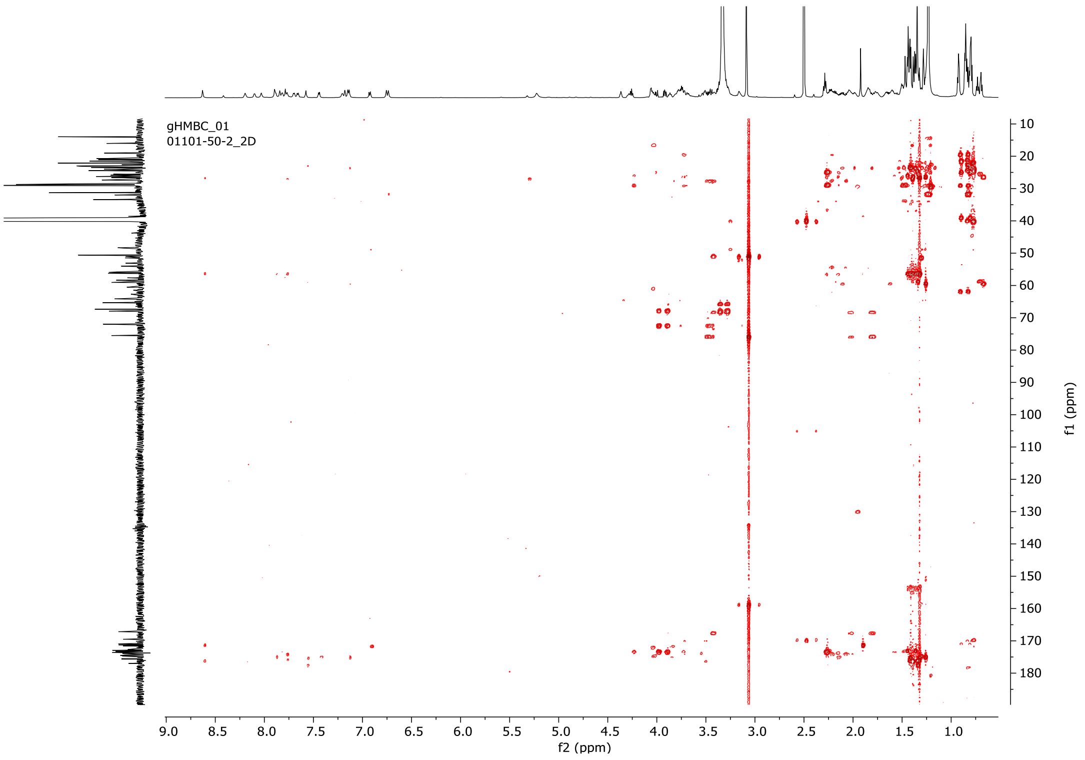

**Fig. S29.** $^{1}H$-$^{13}C$ HMBC NMR data for compound **1** (DMSO-$d_6$).

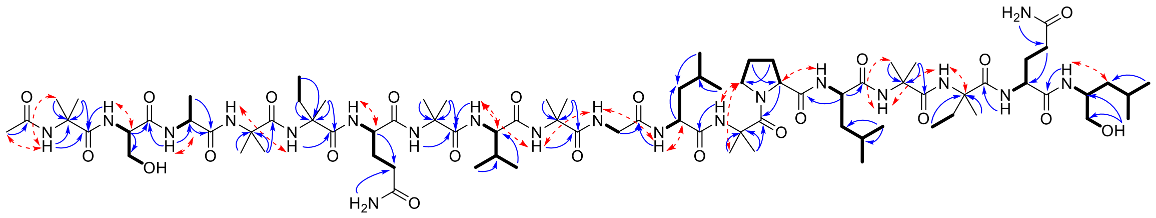

**Fig. S30.** Key HMBC (navy arrows), NOESY (red-dashed arrows) and TOCSY (bolded bounds) correlations for compound **1**.

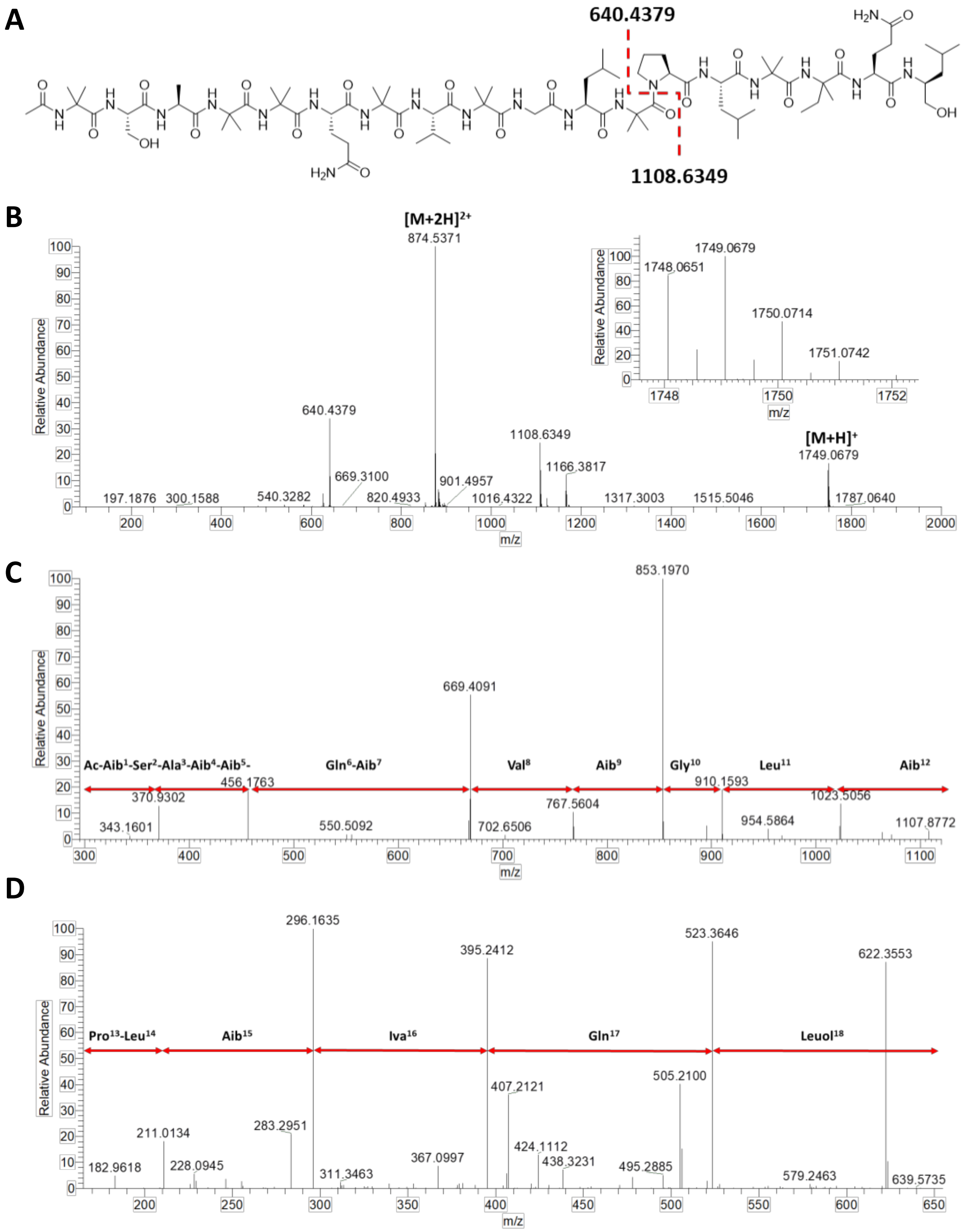

**Fig. S31.** Positive HRESIMS data of compound **2**. A) Structure of **2**. B) Full scan showing in source fragmentation. C and D) HRESIMS$^2$ of fragments $b_{12}^+$ and $y_6^+$, respectively.

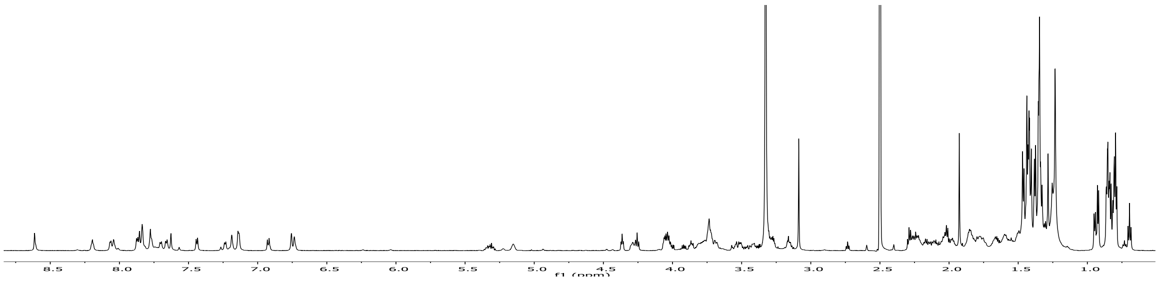

**Fig. S32.** $^{1}H$ NMR data for compound **2** (700 MHz in DMSO-$d_6$).

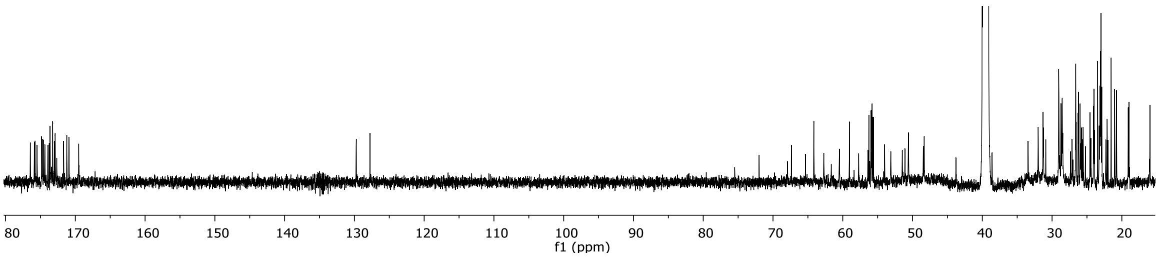

**Fig. S33.** $^{13}C$ NMR data for compound **2** (175 MHz in DMSO-$d_6$).

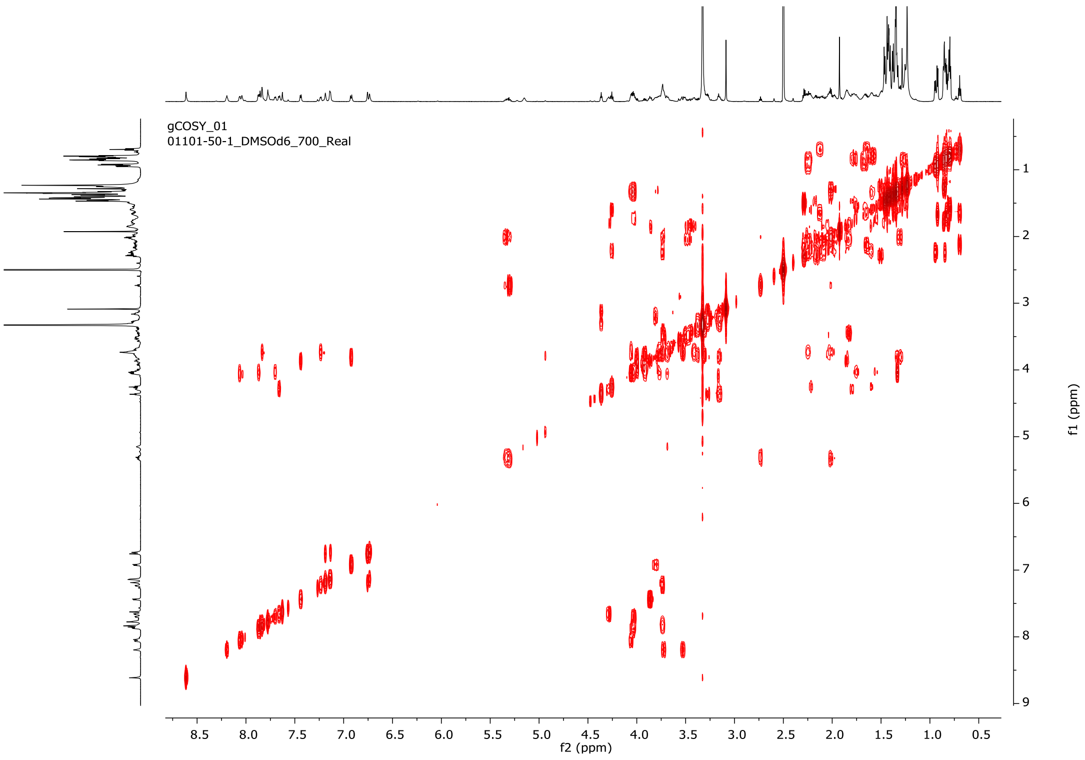

**Fig. S34.** [1]H-[1]H COSY NMR data for compound **2** (DMSO-$d_6$).

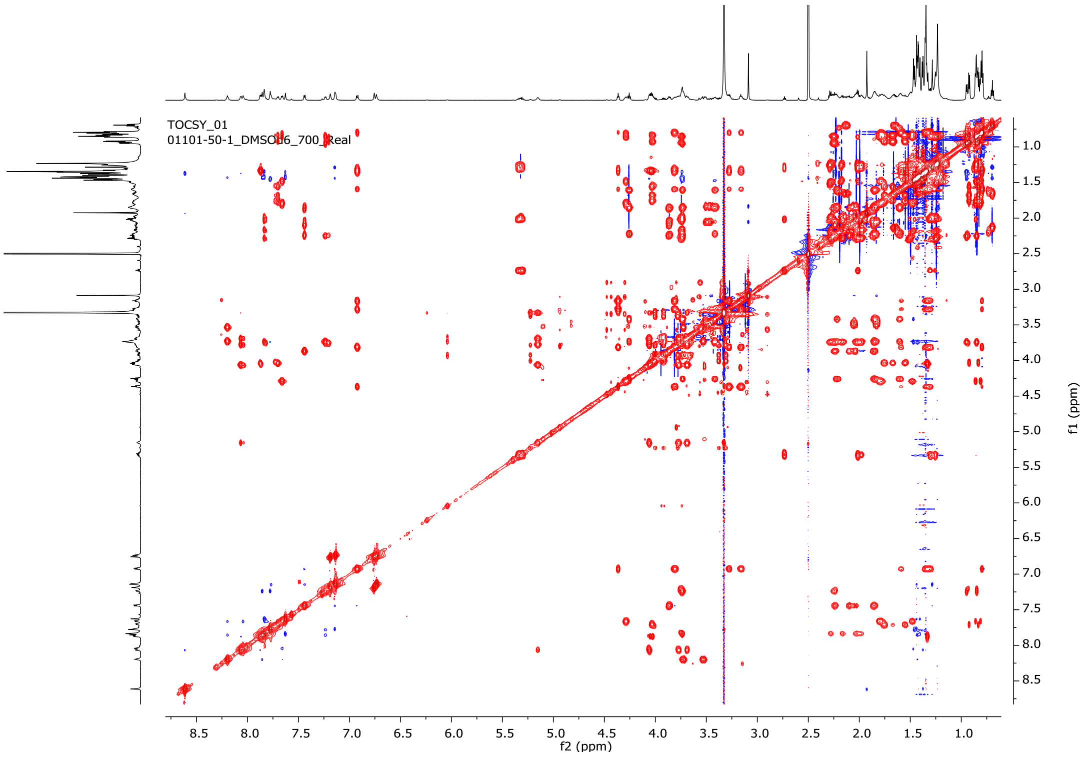

**Fig. S35.** [1]H-[1]H TOCSY NMR data for compound **2** (DMSO-$d_6$).

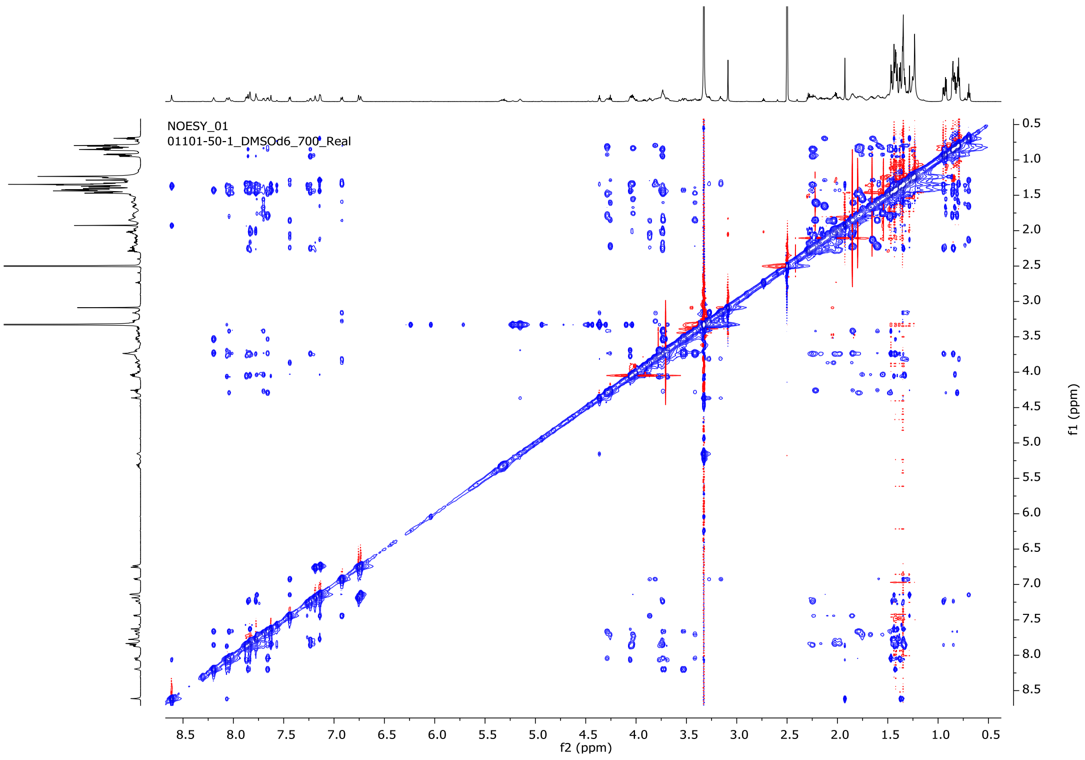

**Fig. S36.** $^1$H-$^1$H NOESY NMR data for compound **2** (DMSO-$d_6$).

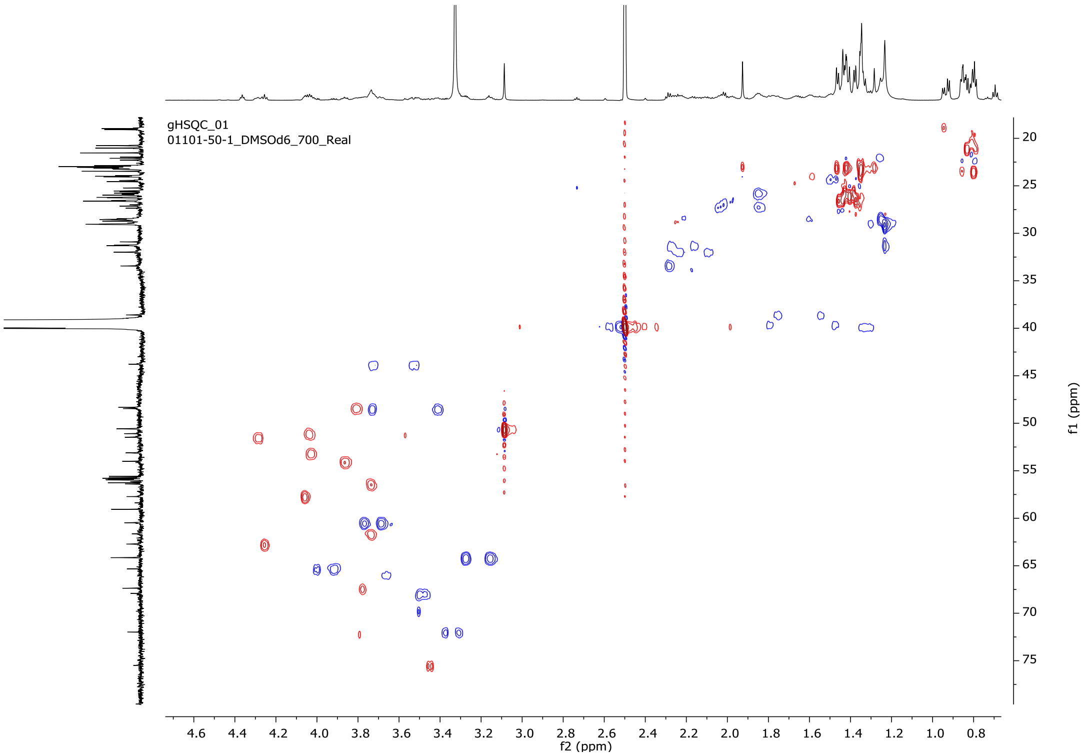

**Fig. S37.** $^1$H-$^{13}$C HSQC NMR data for compound **2** (DMSO-$d_6$).

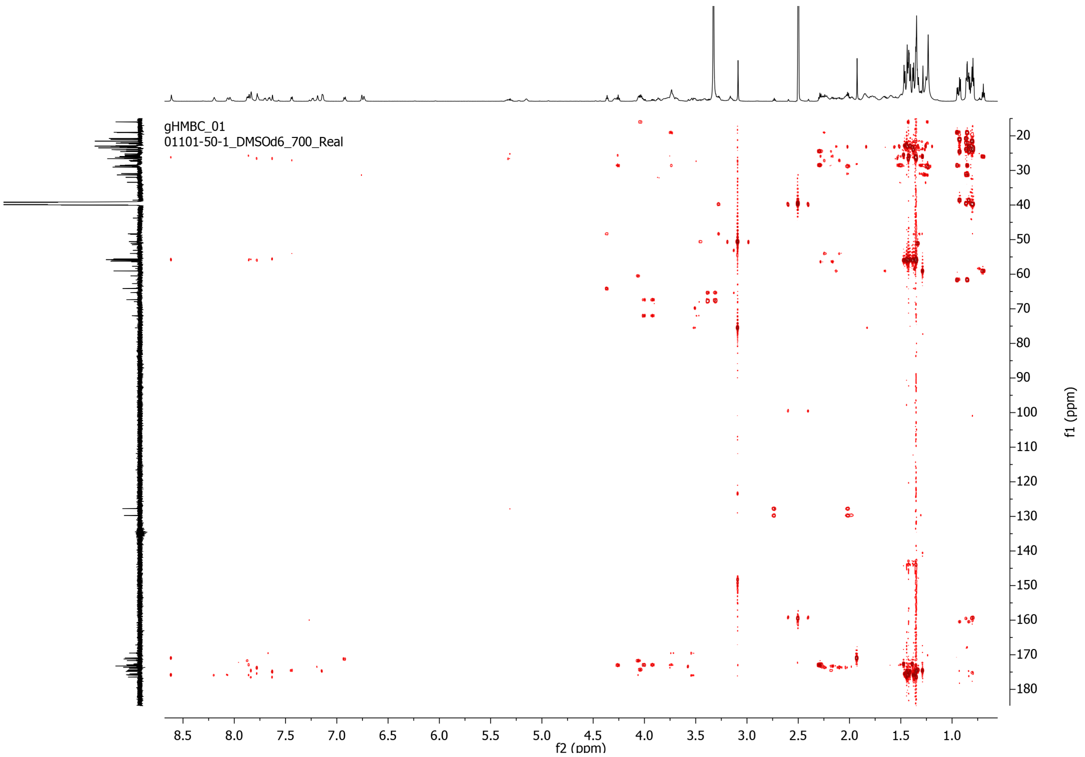

**Fig. S38.** $^{1}H$-$^{13}C$ HMBC NMR data for compound **2** (DMSO-$d_6$).

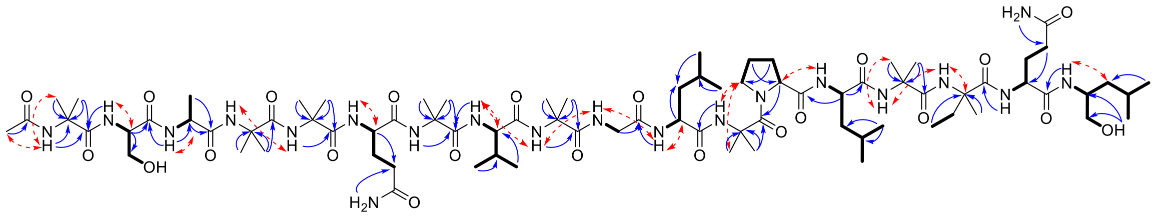

**Fig. S39.** Key HMBC (navy arrows), NOESY (red-dashed arrows) and TOCSY (bolded bounds) correlations for compound **2**.

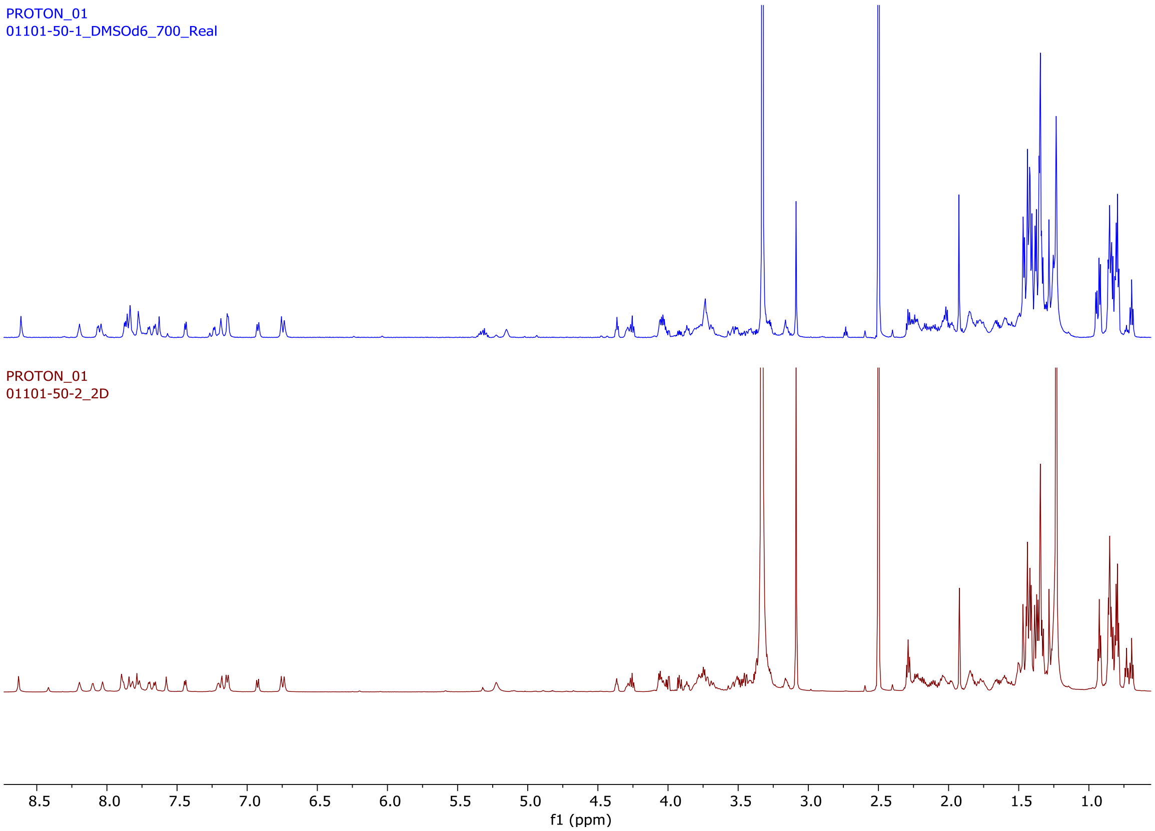

**Fig. S40.** Comparison of $^{1}$H-NMR data for compounds **2** (navy) and **1** (maroon) (both at 700 MHz in DMSO-$d_6$).

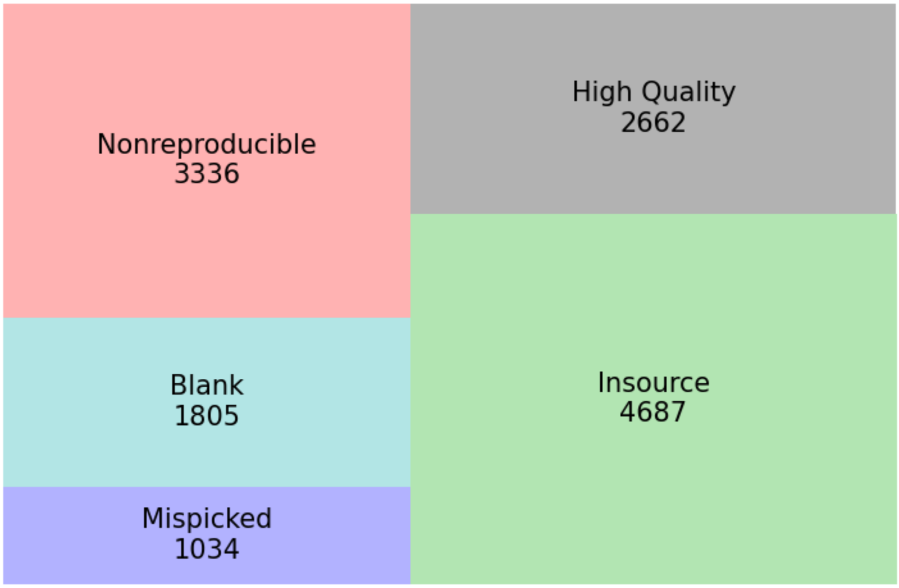

**Fig. S41.** Treemap generated in MPACT describing features of JKS001884 *Trichoderma* sp. fractions indicating features that were found to be non-reproducibly, highly present in blanks, mispicked, or the result of in-source fragmentation.

**Table S1.** Field collection metadata for all *T. septentrionalis* colonies used in this study. The "Dataset" column indicates what each colony was used for.

| Dataset | Colony ID | State | Park | Date of Collection |
|---|---|---|---|---|
| Environmental ITS2 | JKH000038 | New Jersey | Brendon T. Byrne State Forest | 2014-06-24 |
| Environmental ITS2 | JKH000043 | New Jersey | Wharton State Forest | 2014-06-25 |
| Environmental ITS2 | JKH000048 | New Jersey | Wharton State Forest | 2014-06-25 |
| Environmental ITS2 | JKH000052 | New Jersey | Wharton State Forest | 2014-07-23 |
| Environmental ITS2 | JKH000055 | New Jersey | Wharton State Forest | 2014-06-26 |
| Environmental ITS2 | JKH000056 | New Jersey | Wharton State Forest | 2014-06-26 |
| Environmental ITS2 | JKH000057 | New Jersey | Wharton State Forest | 2014-06-26 |
| Environmental ITS2 | JKH000063 | New York | Robert Cushman Murphy County Park | 2014-07-17 |
| Environmental ITS2 | JKH000065 | New Jersey | Brendon T. Byrne State Forest | 2014-07-22 |
| Environmental ITS2 | JKH000067 | New Jersey | Brendon T. Byrne State Forest | 2014-07-22 |
| Environmental ITS2 | JKH000068 | New Jersey | Brendon T. Byrne State Forest | 2014-07-22 |
| Environmental ITS2 | JKH000072 | New Jersey | Wharton State Forest | 2014-07-23 |
| Environmental ITS2 | JKH000073 | New Jersey | Wharton State Forest | 2014-07-23 |
| Environmental ITS2 | JKH000074 | New Jersey | Wharton State Forest | 2014-07-23 |
| Environmental ITS2 | JKH000075 | New Jersey | Wharton State Forest | 2014-07-23 |
| Environmental ITS2 | JKH000076 | New Jersey | Wharton State Forest | 2014-07-23 |
| Environmental ITS2 | JKH000077 | New Jersey | Wharton State Forest | 2014-07-23 |
| Environmental ITS2 | JKH000078 | New Jersey | Wharton State Forest | 2014-07-23 |
| Environmental ITS2 | JKH000079 | New Jersey | Wharton State Forest | 2014-07-23 |
| Environmental ITS2 | JKH000082 | New Jersey | Wharton State Forest | 2014-07-24 |
| Environmental ITS2 | JKH000109 | Florida | Withlacoochee State Forest | 2014-11-15 |
| Environmental ITS2 | JKH000128 | Georgia | George L. Smith State Park | 2015-05-06 |
| Environmental ITS2 | JKH000129 | Georgia | George L. Smith State Park | 2015-05-06 |
| Environmental ITS2 | JKH000130 | Georgia | George L. Smith State Park | 2015-05-06 |
| Environmental ITS2 | JKH000131 | Georgia | George L. Smith State Park | 2015-05-06 |
| Environmental ITS2 | JKH000133 | Georgia | George L. Smith State Park | 2015-05-06 |
| Environmental ITS2 | JKH000134 | Georgia | Yuchi Wildlife Management Area | 2015-05-07 |
| Environmental ITS2 | JKH000135 | Georgia | Yuchi Wildlife Management Area | 2015-05-07 |
| Environmental ITS2 | JKH000136 | Georgia | Yuchi Wildlife Management Area | 2015-05-07 |
| Environmental ITS2 | JKH000141 | Georgia | Magnolia Springs State Park | 2015-05-08 |
| Environmental ITS2 | JKH000142 | Georgia | Magnolia Springs State Park | 2015-05-08 |
| Environmental ITS2 | JKH000156 | North Carolina | Jones Lake State Park | 2015-06-10 |
| Environmental ITS2 | JKH000163 | North Carolina | Singletary Lake State Park | 2015-06-10 |
| Environmental ITS2 | JKH000164 | North Carolina | Singletary Lake State Park | 2015-06-10 |

| | | | | |
|---|---|---|---|---|
| Environmental ITS2 | JKH000165 | North Carolina | Lumber River State Park | 2015-06-11 |
| Environmental ITS2 | JKH000168 | North Carolina | Lumber River State Park | 2015-06-11 |
| Environmental ITS2 | JKH000171 | North Carolina | Lumber River State Park | 2015-06-11 |
| Environmental ITS2 | JKH000172 | North Carolina | Lumber River State Park | 2015-06-12 |
| Environmental ITS2 | JKH000174 | North Carolina | Lumber River State Park | 2015-06-12 |
| Environmental ITS2 | JKH000181 | New York | Robert Cushman Murphy County Park | 2015-07-24 |
| Environmental ITS2 | JKH000185 | Louisiana | Alexander State Forest Wildlife Management Area | 2016-05-26 |
| Environmental ITS2 | JKH000186 | Louisiana | Alexander State Forest Wildlife Management Area | 2016-05-26 |
| Environmental ITS2 | JKH000187 | Louisiana | Clear Creek Wildlife Management Area | 2016-05-28 |
| Environmental ITS2 | JKH000188 | Louisiana | Clear Creek Wildlife Management Area | 2016-05-28 |
| Environmental ITS2 | JKH000192 | Louisiana | Fort Polk Wildlife Management Area | 2016-05-29 |
| Environmental ITS2 | JKH000195 | Louisiana | Fort Polk Wildlife Management Area | 2016-05-29 |
| Environmental ITS2 | JKH000196 | Louisiana | Alexander State Forest Wildlife Management Area | 2016-05-30 |
| Environmental ITS2 | JKH000198 | Louisiana | Alexander State Forest Wildlife Management Area | 2016-05-30 |
| Environmental ITS2 | JKH000199 | Louisiana | Alexander State Forest Wildlife Management Area | 2016-05-30 |
| Environmental ITS2 | JKH000204 | Florida | Tate's Hell State Forest | 2016-06-27 |
| Environmental ITS2 | JKH000207 | Florida | Lake Talquin State Forest | 2016-06-28 |
| Environmental ITS2 | JKH000212 | Florida | Lake Talquin State Forest | 2016-06-28 |
| Environmental ITS2 | JKH000213 | Florida | Lake Talquin State Forest | 2016-06-29 |
| Environmental ITS2 | JKH000215 | Florida | Lake Talquin State Forest | 2016-06-29 |
| Environmental ITS2 | JKH000219 | Florida | Wakulla State Forest | 2016-06-30 |
| Environmental ITS2 | JKH000226 | Florida | Wakulla State Forest | 2016-06-30 |
| Environmental ITS2 | JKH000232 | New Jersey | Brendon T. Byrne State Forest | 2017-07-24 |
| Environmental ITS2 | JKH000233 | New Jersey | Brendon T. Byrne State Forest | 2017-07-24 |
| Environmental ITS2 | JKH000234 | New Jersey | Brendon T. Byrne State Forest | 2017-07-24 |
| Environmental ITS2 | JKH000235 | New Jersey | Brendon T. Byrne State Forest | 2017-07-24 |
| Environmental ITS2 | JKH000237 | New Jersey | Brendon T. Byrne State Forest | 2017-07-24 |
| Environmental ITS2 | JKH000240 | New Jersey | Brendon T. Byrne State Forest | 2017-07-24 |
| Environmental ITS2 | JKH000241 | New Jersey | Brendon T. Byrne State Forest | 2017-07-24 |
| Environmental ITS2 | JKH000242 | New Jersey | Brendon T. Byrne State Forest | 2017-07-24 |
| Environmental ITS2 | JKH000243 | New Jersey | Wharton State Forest | 2017-07-25 |
| Environmental ITS2 | JKH000244 | New Jersey | Wharton State Forest | 2017-07-25 |
| Environmental ITS2 | JKH000245 | New Jersey | Wharton State Forest | 2017-07-25 |
| Environmental ITS2 | JKH000247 | New Jersey | Wharton State Forest | 2017-07-25 |
| Environmental ITS2 | JKH000248 | New Jersey | Wharton State Forest | 2017-07-25 |
| Environmental ITS2 | JKH000253 | New Jersey | Wharton State Forest | 2017-07-26 |
| Environmental ITS2 | JKH000256 | New Jersey | Wharton State Forest | 2017-07-26 |
| Environmental ITS2 | JKH000258 | New Jersey | Wharton State Forest | 2017-07-26 |
| Environmental ITS2 | JKH000260 | New Jersey | Wharton State Forest | 2017-07-26 |
| Environmental ITS2 | JKH000262 | New Jersey | Wharton State Forest | 2017-07-26 |

| | | | | |
|---|---|---|---|---|
| Environmental ITS2 | JKH000263 | New Jersey | Wharton State Forest | 2017-07-26 |
| Environmental ITS2 | JKH000264 | New Jersey | Wharton State Forest | 2017-07-27 |
| Environmental ITS2 | JKH000265 | New Jersey | Wharton State Forest | 2017-07-27 |
| Environmental ITS2 | JKH000266 | New Jersey | Wharton State Forest | 2017-07-27 |
| Environmental ITS2 | JKH000267 | New Jersey | Wharton State Forest | 2017-07-27 |
| Environmental LC-MS/MS | JKH000070 | New Jersey | Brendon T. Byrne State Forest | 2014-07-22 |
| Environmental LC-MS/MS | JKH000073 | New Jersey | Wharton State Forest | 2014-07-23 |
| Environmental LC-MS/MS | JKH000074 | New Jersey | Wharton State Forest | 2014-07-23 |
| Environmental LC-MS/MS | JKH000076 | New Jersey | Wharton State Forest | 2014-07-23 |
| Environmental LC-MS/MS | JKH000133 | Georgia | George L. Smith State Park | 2015-05-06 |
| Environmental LC-MS/MS | JKH000145 | Louisiana | Alexander Wildlife Management Area | 2015-05-09 |
| Environmental LC-MS/MS | JKH000150 | North Carolina | William B. Umstead State Park | 2015-06-09 |
| Environmental LC-MS/MS | JKH000151 | North Carolina | William B. Umstead State Park | 2015-06-09 |
| Environmental LC-MS/MS | JKH000152 | North Carolina | William B. Umstead State Park | 2015-06-09 |
| Environmental LC-MS/MS | JKH000154 | North Carolina | William B. Umstead State Park | 2015-06-09 |
| Environmental LC-MS/MS | JKH000155 | North Carolina | William B. Umstead State Park | 2015-06-09 |
| Environmental LC-MS/MS | JKH000156 | North Carolina | Jones Lake State Park | 2015-06-01 |
| Environmental LC-MS/MS | JKH000157 | North Carolina | Jones Lake State Park | 2015-06-01 |
| Environmental LC-MS/MS | JKH000158 | North Carolina | Jones Lake State Park | 2015-06-01 |
| Environmental LC-MS/MS | JKH000160 | North Carolina | Jones Lake State Park | 2015-06-01 |
| Environmental LC-MS/MS | JKH000161 | North Carolina | Singletary Lake State Park | 2015-06-01 |
| Environmental LC-MS/MS | JKH000162 | North Carolina | Singletary Lake State Park | 2015-06-01 |
| Environmental LC-MS/MS | JKH000163 | North Carolina | Singletary Lake State Park | 2015-06-01 |
| Environmental LC-MS/MS | JKH000164 | North Carolina | Singletary Lake State Park | 2015-06-01 |
| Environmental LC-MS/MS | JKH000165 | North Carolina | Lumber River State Park | 2015-06-11 |
| Environmental LC-MS/MS | JKH000167 | North Carolina | Lumber River State Park | 2015-06-11 |
| Environmental LC-MS/MS | JKH000168 | North Carolina | Lumber River State Park | 2015-06-11 |
| Environmental LC-MS/MS | JKH000169 | North Carolina | Lumber River State Park | 2015-06-11 |
| Environmental LC-MS/MS | JKH000170 | North Carolina | Lumber River State Park | 2015-06-11 |
| Environmental LC-MS/MS | JKH000171 | North Carolina | Lumber River State Park | 2015-06-11 |
| Environmental LC-MS/MS | JKH000173 | North Carolina | Lumber River State Park | 2015-06-12 |
| Environmental LC-MS/MS | JKH000174 | North Carolina | Lumber River State Park | 2015-06-12 |
| Environmental LC-MS/MS | JKH000175 | North Carolina | Lumber River State Park | 2015-06-12 |
| Environmental LC-MS/MS | JKH000176 | North Carolina | Lumber River State Park | 2015-06-12 |
| Environmental LC-MS/MS | JKH000177 | North Carolina | Lumber River State Park | 2015-06-12 |
| Environmental LC-MS/MS | JKH000179 | North Carolina | Lumber River State Park | 2015-06-12 |
| Environmental LC-MS/MS | JKH000180 | North Carolina | Lumber River State Park | 2015-06-12 |
| Environmental LC-MS/MS | JKH000181 | New York | Robert Cushman Murphy County Park | 2015-07-24 |
| Environmental LC-MS/MS | JKH000183 | New York | Robert Cushman Murphy County Park | 2015-07-24 |

| | | | | |
|---|---|---|---|---|
| Environmental LC-MS/MS | JKH000206 | Florida | Lake Talquin State Forest | 2016-06-28 |
| Environmental LC-MS/MS | JKH000208 | Florida | Lake Talquin State Forest | 2016-06-28 |
| Environmental LC-MS/MS | JKH000209 | Florida | Lake Talquin State Forest | 2016-06-28 |
| Environmental LC-MS/MS | JKH000211 | Florida | Lake Talquin State Forest | 2016-06-28 |
| Environmental LC-MS/MS | JKH000212 | Florida | Lake Talquin State Forest | 2016-06-28 |
| Environmental LC-MS/MS | JKH000213 | Florida | Lake Talquin State Forest | 2016-06-29 |
| Environmental LC-MS/MS | JKH000214 | Florida | Lake Talquin State Forest | 2016-06-29 |
| Environmental LC-MS/MS | JKH000215 | Florida | Lake Talquin State Forest | 2016-06-29 |
| Environmental LC-MS/MS | JKH000217 | Florida | Lake Talquin State Forest | 2016-06-29 |
| Environmental LC-MS/MS | JKH000219 | Florida | Wakulla State Forest | 2016-06-30 |
| Environmental LC-MS/MS | JKH000221 | Florida | Wakulla State Forest | 2016-06-30 |
| Environmental LC-MS/MS | JKH000223 | Florida | Wakulla State Forest | 2016-06-30 |
| Environmental LC-MS/MS | JKH000225 | Florida | Wakulla State Forest | 2016-06-30 |
| Environmental LC-MS/MS | JKH000226 | Florida | Wakulla State Forest | 2016-06-30 |
| Environmental LC-MS/MS | JKH000227 | Florida | Wakulla State Forest | 2016-06-30 |
| *Trichoderma* isolations | JKH000037 | New Jersey | Brendon T. Byrne State Forest | 2014-06-24 |
| *Trichoderma* isolations | JKH000038 | New Jersey | Brendon T. Byrne State Forest | 2014-06-24 |
| *Trichoderma* isolations | JKH000066 | New Jersey | Brendon T. Byrne State Forest | 2014-07-22 |
| *Trichoderma* infection | JKH000285 | New York | Robert Cushman Murphy County Park | 2018-06-20 |
| Time-course infection | JKH000292 | New Jersey | Wharton State Forest | 2018-06-27 |
| 2019 extract experiments | JKH000365 | North Carolina | Lumber River State Park | 2019-06-05 |
| 2019 extract experiments | JKH000377 | North Carolina | William B. Umstead State Park | 2019-06-06 |
| 2019 extract experiments | JKH000380 | North Carolina | William B. Umstead State Park | 2019-06-06 |
| 2020 extract experiments | JKH000397 | New Jersey | Wharton State Forest | 2020-06-24 |
| 2020 extract experiments | JKH000408 | New Jersey | Wharton State Forest | 2020-06-25 |
| 2020 extract experiments | JKH000409 | New Jersey | Wharton State Forest | 2020-06-25 |
| 2020 extract experiments | JKH000410 | New Jersey | Wharton State Forest | 2020-06-25 |
| 2020 extract experiments | JKH000411 | New Jersey | Wharton State Forest | 2020-06-25 |
| 2020 extract experiments | JKH000412 | New Jersey | Wharton State Forest | 2020-06-25 |
| 2020 extract experiments | JKH000414 | New Jersey | Wharton State Forest | 2020-06-25 |
| 2020 extract experiments | JKH000419 | New Jersey | Brendan T. Byrne State Forest | 2020-06-26 |
| 2020 extract experiments | JKH000422 | New Jersey | Brendan T. Byrne State Forest | 2020-06-26 |

**Table S2. *Trichoderma* isolations and identification.** The three *Trichoderma* strains used in this study with their associated isolation metadata and the sequences used for species classification.

| Strain ID | Classification | Colony ID | GenBank ID | Gene |
|---|---|---|---|---|
| JKS001879 | *Trichoderma koningiopsis* | JKH000037 | OM967104 | ITS |
| | | | ON364341 | TEF1 |
| | | | ON113306 | RPB2 |
| JKS001884 | *Trichoderma virens* | JKH000038 | OM967105 | ITS |
| | | | ON364342 | TEF1 |
| | | | ON113307 | RPB2 |
| JKS001921 | *Trichoderma simmonsii* | JKH000066 | OM967106 | ITS |
| | | | ON364343 | TEF1 |
| | | | ON113308 | RPB2 |

**Table S3.** Primer sequences used in this study.

| Primer name | Primer sequence (5' to 3') | Reference |
|---|---|---|
| fITS7 | GTG ART CAT CGA ATC TTT G | Ihrmark et al. 2012 |
| ITS4 | TCC TCC GCT TAT TGA TAT GC | White et al. 1990 |
| ITS1 | TCC GTA GGT GAA CCT GCG G | White et al. 1990 |
| EF1 | ATG GGT AAG GAR GAC AAG AC | Cai and Druzhinina 2021 |
| EF2 | GGA RGT ACC AGT SAT CAT GTT | Cai and Druzhinina 2021 |
| fRPB2-5f | GAY GAY MGW GAT CAY TTY GG | Liu et al. 1999 |
| fRPB2-7cr | CCC ATR GCT TGY TTR CCC AT | Liu et al. 1999 |

**Table S4:** Summer 2020 ant waste production bioassays. Multiple trials were sometimes run using the same control subcolonies.

| Exp. # | Date | Treatment(s) | Controls (+/–)[a] | Abbrev. Colony ID | Excluded from analysis |
|---|---|---|---|---|---|
| 1 | 7-16 | ant number[b] | | 408 | yes – experimental design test, negative controls not tested |
| | | | | 414 | yes – experimental design test, negative controls not tested |
| | | | | 416 | yes – experimental design test, negative controls not tested |
| 2 | 7-22 | lab acclimation[c] | MB1081, DMSO, NT | 410 | yes – before seasonal threshold |
| | | | | 411 | yes – not lab acclimated |
| 3 | 7-23 | lab acclimation | MB1081, DMSO, NT | 422 | yes – negative control weeding equal to crude weeding |
| | | | | 409 | yes – not lab acclimated |
| 4 | 7-27 | fractions B-F | MB1081, DMSO, NT | 422 | yes – before seasonal threshold |
| 5 | 7-29 | fractions B-F | MB1081, DMSO, NT | 397 | yes – negative control waste exceeded crude waste |
| 6 | 7-31 | extract batch[d] * treatment dose[e] | MB1081, MB0895, DMSO, NT | 410 | yes – negative control waste exceeded crude waste |
| 7 | 8-11 | garden disturbance[f] | | 422 | yes – experimental design test, extracts not tested |
| 8 | 8-23 | humidity[g] | | 412 | yes – experimental design test, extracts not tested |
| 9 | 8-24 | humidity * crude extract | MB1084, DMSO, NT | 412 | **no** – only 100% humidity used |
| 10 | 8-28 | humidity * crude extract | MB1081, MB1084, DMSO, NT | 419 | **no** – only 100% humidity used |
| 11 | 9-01 | fractions B-E, G | MB1081, DMSO, NT | 414 | **no** |
| 12 | 9-06 | fractions B-E, G | MB1081, DMSO, NT | 408 | **no** |
| 13 | 9-08 | pure compounds | MB1081, DMSO, NT | 408 | **no** |
| 14 | 9-10 | pure compounds | MB1081, DMSO, NT | 422 | **no** |
| 15 | 9-13 | fractions A-F | MB1084, DMSO, NT | 397 | **no** |
| 16 | 9-16 | pure compounds | MB1081, DMSO, NT | 412 | **no** |

| | | | | | |
|---|---|---|---|---|---|
| 17 | 9-20 | *Trichoderma* pre-extraction growth time[h] | MB1080, DMSO, NT | 414 | yes – garden and waste appeared infected at sampling |
| | | | | 396 | **no** |
| 18 | 9-22 | pure compounds | MB1084, DMSO, NT | 392 | yes – crude extract did not produce weeding response |
| 19 | 9-24 | *Trichoderma* pre-extraction growth time; fractions A-F | MB1084, DMSO, NT | 419 | **no** – only MB1080 (7 d) was used from growth experiment |

[a]Crude *Trichoderma* extract was used as the positive control. Its induction of ant weeding was repeatedly demonstrated in 2019 and was the phenotype that we aimed to replicate in 2020. These crude extracts contain the metabolites tested in the fractionation and pure compound experiments. DMSO treatment (0.5%) was used as the negative solvent-only control because all extracts and compounds were dissolved in 0.5% DMSO. The weeding response to DMSO was expected to be similar to the no treatment (NT) control, which was minimally manipulated and thus not expected to display an ant weeding response.
[b]This experiment compared the amount of waste produced by ant subcolonies treated with crude extract but containing either 10, 8, or 7 ants.
[c]These experiments tested the response of both acclimated and non-acclimated subcolonies to crude extract. Colonies were considered acclimated when the fungus garden color changed from brown (it's natural color in the field) to yellow due to the incorporation of sterile cornmeal in the lab.
[d]These experiments compared two replicate extracts from the same *Trichoderma* strain. MB0895 was extracted in 2019 and kept at -20°C; MB1081 was extracted in 2020.
[e]Two different volumes of *Trichoderma* innocula were compared: 200 µL and 400 µL
[f]This experiment tested the effect of physically manipulating fungus gardens using forceps, and compared fungus gardens that were touched minimally to those that were touched an additional 5 or 10 times.
[g]These experiments tested the impact of humidity on ant weeding. Box humidity was adjusted by adding water to the base plaster 1 mL at a time until it was saturated (100% humidity). This volume was then halved (50% humidity), quartered (25% humidity), or no water was added (0% humidity).
[h]*Trichoderma* was grown for 3 d (MB1079), 7 d (MB1080), or 14 d (MB1082) prior to extraction.

**Table S5.** Values for transferring ions into the collision cell per scan time for metabolomics data acquisition.

| Time[a](%) | Collision RF[b] (Vpp) | Transfer Time (µs) | Collision (%) |
|---|---|---|---|
| 0 | 450.0 | 70.0 | 125 |
| 25 | 550.0 | 75.0 | 100 |
| 50 | 800.0 | 90.0 | 100 |
| 75 | 1100.0 | 95.0 | 75 |

[a] Collision stepping switch time (proportion of the scan time)
[b] RF, radio frequency; Vpp, Peak-to-Peak voltage Vpp, volts peak to peak

**Table S6.** CID energies for MS/MS data acquisition for metabolomics data acquisition.

| Type | Mass | Width | Collision | Charge State |
|---|---|---|---|---|
| Base | 100.00 | 4.00 | 22.00 | 1 |
| Base | 100.00 | 4.00 | 18.00 | 2 |
| Base | 300.00 | 5.00 | 27.00 | 1 |
| Base | 300.00 | 5.00 | 22.00 | 2 |
| Base | 500.00 | 6.00 | 35.00 | 1 |
| Base | 500.00 | 6.00 | 30.00 | 2 |
| Base | 1000.00 | 8.00 | 45.00 | 1 |
| Base | 1000.00 | 8.00 | 35.00 | 2 |
| Base | 2000.00 | 10.00 | 50.00 | 1 |
| Base | 2000.00 | 10.00 | 50.00 | 2 |

The mass of the internal calibrant was excluded from the MS/MS list using a mass range of *m/z* 621.5–623.0.

**Table S7.** NMR data for compounds **1** and **2** (700 MHz for $^1$H and 175 MHz for $^{13}$C, DMSO-$d_6$).

| residue | position | **1** | | | **2** | | |
|---|---|---|---|---|---|---|---|
| | | $\delta_C$ | type | $\delta_H$, m, (J in Hz) | $\delta_C$ | type | $\delta_H$, m (J in Hz) |
| Ac | C=O | 171.2 | C | | 170.9 | C | |
| | $CH_3$ | 23.5 | $CH_3$ | 1.92, s | 23.0 | $CH_3$ | 1.93, s |
| Aib$^1$ | C=O | 175.9 | C | | 175.9 | C | |
| | $\alpha$ | 56.3 | C | | 55.8 | C | |
| | $\beta_1$ | 26.6 | $CH_3$ | 1.35, s | 26.6 | $CH_3$ | 1.37, s |
| | $\beta_2$ | 26.6 | $CH_3$ | 1.44, s | 26.6 | $CH_3$ | 1.44, s |
| | NH | | | 8.63, s | | | 8.61, s |
| Ser$^2$ | C=O | 171.6 | C | | 171.7 | C | |
| | $\alpha$ | 57.6 | CH | 4.06* | 57.7 | CH | 4.06* |
| | $\beta$ | 60.5 | $CH_2$ | 3.69, m | 60.5 | $CH_2$ | 3.70, m |
| | | | | 3.77, m | | | 3.77, m |
| | NH | | | 8.10, d (5.8) | | | 8.06, d (5.9) |
| | OH | | | 5.23, brs | | | 5.15 brs |
| Ala$^3$ | C=O | 174.4 | C | | 174.4 | C | |
| | $\alpha$ | 55.0 | C | 4.05* | 51.1 | C | 4.04* |
| | $\beta$ | 16.0 | $CH_3$ | 1.33* | 16.0 | $CH_3$ | 1.34* |
| | NH | | | 7.89, d* | | | 7.87, d (6.7) |
| Aib$^4$ | C=O | 173.0 | C | | 174.5 | C | |
| | $\alpha$ | 56.2 | C | | 59.1 | C | |
| | $\beta_1$ | 23.2 | $CH_3$ | 1.41, s | 23.1 | $CH_3$ | 1.28, s |
| | $\beta_2$ | 26.2 | $CH_3$ | 1.37, s | 26.2 | $CH_3$ | 1.34, s |
| | NH | | | 7.84, s | | | 7.83, s |
| Iva$^5$/Aib$^5$ | C=O | 175.7 | C | | 176.4 | C | |
| | $\alpha$ | 58.4 | C | | 55.7 | C | |
| | $\beta_{1a}$ | 26.3 | $CH_2$ | 1.61* | 27.1 | $CH_3$ | 1.35, s |
| | $\beta_{1b}$ | | | 2.19* | - | - | - |
| | $\beta_2$ | 26.6 | $CH_3$ | 1.43, s | 26.6 | $CH_3$ | 1.44, s |
| | $\gamma$ | 7.2 | $CH_3$ | 0.72, t (7.5) | - | - | - |
| | NH | | | 7.58, s | | | 7.62, s |
| Gln$^6$ | C=O | 174.7 | | | 174.7 | | |
| | $\alpha$ | 54.0 | CH | 3.86, ddd (11.3, 7.4, 4.0) | 54.0 | CH | 3.88* |
| | $\beta$ | 25.6 | $CH_2$ | 1.85, m | 27.1 | $CH_2$ | 1.84, m |
| | $\gamma_{1a}$ | 32.0 | $CH_2$ | 2.09* | 32.0 | $CH_2$ | 2.10* |
| | $\gamma_{1b}$ | | | 2.23* | | | 2.29* |
| | $\delta$ | 173.8 | C | | 173.7 | C | |
| | NH | | | 7.44, d (7.4) | | | 7.44, d (7.4) |
| | $NH_2$ | | | 6.74, brs | | | 6.73, brs |
| | | | | 7.13, brs | | | 7.14, brs |
| Aib$^7$ | C=O | 175.5 | C | | 175.9 | C | |
| | $\alpha$ | 56.3 | C | | 55.8 | C | |
| | $\beta_1$ | 26.3 | $CH_3$ | 1.37, s | 26.5 | $CH_3$ | 1.37, s |
| | $\beta_2$ | 25.8 | $CH_3$ | 1.42, s | 26.2 | $CH_3$ | 1.40, s |
| | NH | | | 7.79, s | | | 7.78, s |
| Val$^8$ | C=O | 172.7 | C | | 171.7 | C | |
| | $\alpha$ | 61.5 | CH | 3.74* | 61.7 | CH | 3.76* |
| | $\beta$ | 28.7 | CH | 2.24* | 28.7 | CH | 2.24* |
| | $\gamma_1$ | 23.5 | $CH_3$ | 0.85, d (6.6) | 23.5 | $CH_3$ | 0.86, d (6.5) |
| | $\gamma_2$ | 19.6 | $CH_3$ | 0.93, d (6.5) | 19.1 | $CH_3$ | 0.92, d (6.5) |
| | NH | | | 7.20, d (7.0) | | | 7.24, d (6.9) |
| Aib$^9$ | C=O | 173.9 | C | | 175.9 | C | |
| | $\alpha$ | 56.0 | C | | 55.8 | C | |
| | $\beta_1$ | 25.9 | $CH_3$ | 1.39, s | 26.5 | $CH_3$ | 1.37, s |
| | $\beta_2$ | 23.1 | $CH_3$ | 1.47, s | 26.2 | $CH_3$ | 1.40, s |
| | NH | | | 7.88, s | | | 7.87, s |
| Gly$^{10}$ | C=O | 169.5 | C | | 169.5 | C | |
| | $\alpha_1$ | 43.8 | $CH_2$ | 3.53* | 43.8 | $CH_2$ | 3.53* |
| | $\alpha_2$ | | | 3.73* | | | 3.73* |
| | NH | | | 8.19, t (5.8) | | | 8.19, t (5.0) |
| Leu$^{11}$ | C=O | 174.8 | C | | 174.7 | C | |
| | $\alpha$ | 51.5 | CH | 4.28, m | 51.5 | CH | 4.29, t (8.0) |
| | $\beta_{1a}$ | 40.0 | $CH_2$ | 1.50* | 40.0 | $CH_2$ | 1.49* |
| | $\beta_{1b}$ | | | 1.81* | | | 1.80* |
| | $\gamma$ | 24.1 | CH | 1.78* | 24.1 | CH | 1.78* |
| | $\delta_1$ | 23.2 | $CH_3$ | 0.81, d (6.2) | 23.5 | $CH_3$ | 0.81, d (6.2) |
| | $\delta_2$ | 23.3 | $CH_3$ | 0.86, d (6.2) | 23.5 | $CH_3$ | 0.86, d (6.2) |
| | NH | | | 7.66, d (8.0) | | | 7.66, d (7.9) |
| Aib$^{12}$ | C=O | 173.0 | C | | 174.5 | C | |
| | $\alpha$ | | C | | 55.9 | C | |
| | $\beta_1$ | 23.1 | $CH_3$ | 1.42, s | 23.1 | $CH_3$ | 1.42, s |
| | $\beta_2$ | 26.2 | $CH_3$ | 1.44, s | 26.5 | $CH_3$ | 1.46, s |
| | NH | | | 8.03, s | | | 8.04, s |

| | | | | | | | |
|---|---|---|---|---|---|---|---|
| Pro[13] | C=O | 173.9 | C | | 173.7 | C | |
| | $\alpha$ | 62.7 | CH | 4.26, t (8.0) | 62.7 | CH | 4.26, t (8.0) |
| | $\beta$ | 32.0 | $CH_2$ | 2.21* | 33.4 | $CH_2$ | 2.23* |
| | $\gamma_{1a}$ | 25.6 | $CH_2$ | 1.60, m | 25.8 | $CH_2$ | 1.60, m |
| | $\gamma_{1b}$ | | | 1.85* | | | 1.85* |
| | $\delta_{1a}$ | 48.4 | $CH_2$ | 3.41, dd (11.3, 2.7) | 48.4 | $CH_2$ | 3.42, dd (11.4, 2.7) |
| | $\delta_{1b}$ | | | 3.73* | | | 3.73* |
| Leu[14] | C=O | 173.0 | C | | 173.0 | C | |
| | $\alpha$ | 53.1 | CH | 4.03, m | 53.1 | CH | 4.02, m |
| | $\beta_{1a}$ | 38.6 | $CH_2$ | 1.55* | 38.6 | $CH_2$ | 1.55* |
| | $\beta_{1b}$ | | | 1.76* | | | 1.75* |
| | $\gamma$ | 24.6 | CH | 1.67, m | 24.6 | CH | 1.60, m |
| | $\delta_1$ | 20.8 | $CH_3$ | 0.83, d (6.5) | 20.8 | $CH_3$ | 0.83, d (6.6) |
| | $\delta_2$ | 19.0 | $CH_3$ | 0.92, d (6.5) | 19.0 | $CH_3$ | 0.92, d (6.5) |
| | NH | | | 7.70, d (7.3) | | | 7.70, d (7.1) |
| Aib[15] | C=O | 175.5 | C | | 174.8 | C | |
| | $\alpha$ | 55.9 | C | | 56.3 | C | |
| | $\beta_1$ | 23.5 | $CH_3$ | 1.41, s | 23.3 | $CH_3$ | 1.41, s |
| | $\beta_2$ | 26.3 | $CH_3$ | 1.45, s | 26.5 | $CH_3$ | 1.44, s |
| | NH | | | 7.77, s | | | 7.86, s |
| Iva[16] | C=O | 174.9 | C | | 171.2 | C | |
| | $\alpha$ | 59.1 | C | | 59.1 | C | |
| | $\beta_{1a}$ | 26.3 | $CH_2$ | 1.66* | 26.0 | $CH_2$ | 1.66* |
| | $\beta_{1b}$ | | | 2.13* | | | 2.13* |
| | $\beta_2$ | 7.2 | $CH_3$ | 0.70, t (7.5) | 7.2 | $CH_3$ | 0.69, t (7.5) |
| | $\gamma$ | 23.5 | $CH_3$ | 1.28, s | 23.1 | $CH_3$ | 1.28, s |
| | NH | | | 7.14, s | | | 7.13, s |
| Gln[17] | C=O | | | | | | |
| | $\alpha$ | 56.3 | CH | 3.75* | 56.4 | CH | 3.73* |
| | $\beta$ | 26.3 | $CH_2$ | 2.01* | 27.0 | $CH_2$ | 2.01* |
| | $\gamma_{1a}$ | 31.3 | $CH_2$ | 2.17* | 31.3 | $CH_2$ | 2.17, m |
| | $\gamma_{1b}$ | | | 2.29, m | | | 2.28, m |
| | $\delta$ | 173.3 | C | | 173.2 | C | |
| | NH | | | 7.82, d (4.5) | | | 7.83, d* |
| | $NH_2$ | | | 6.76, brs | | | 6.76, brs |
| | | | | 7.18, brs | | | 7.19, brs |
| Ileol[18] | | | | | | | |
| | $\alpha$ | 48.3 | CH | 3.80* | 48.4 | CH | 3.81* |
| | $\beta$ | 40.0 | $CH_2$ | 1.33* | 40.0 | $CH_2$ | 1.33* |
| | $\gamma$ | 24.6 | CH | 1.59* | 24.1 | CH | 1.59* |
| | $\delta$ | 21.5 | $CH_3$ | 0.79, d (6.5) | 21.0 | $CH_3$ | 0.80, d (6.3) |
| | $\gamma_2$ | 23.1 | $CH_3$ | 0.80, d (6.5) | 21.0 | $CH_3$ | 0.80, d (6.3) |
| | $\beta'_1$ | 64.2 | $CH_2$ | 3.16, m | 64.2 | $CH_2$ | 3.17, m |
| | $\beta'_2$ | | | 3.27, m | | | 3.27, m |
| | NH | | | 6.93, d (9.3) | | | 6.94, d (9.3) |
| | OH | | | 4.37, t (5.9) | | | 4.37, t (5.9) |

***Overlapped signals. Note: Signals for Aib residues may be interchangeable.**

**Table S8.** Percent abundance for each of the three most abundant metabolites for each JKS001884 *Trichoderma* sp. fraction (from Fig. 4C; percentages calculated using total ion abundance per fraction). Together, the ion abundances of these three features represented a combined 35.5% and 75.5% of total metabolite abundance in fractions D and E, respectively. Further exploration of these features confirmed that all three represent peptaibols, two of which have masses consistent with the peptaibols trichodermides B/C and D/E, although several other peptaibols have similar masses.

| Metabolite | *m/z* | %A | %B | %C | %D | %E | %F |
|---|---|---|---|---|---|---|---|
| Peptaibol_1197 | 1197.7557 | 1.7 | 1.7 | 0.8 | 9.5 | 47.0 | 7.9 |
| Peptaibol_1183 | 1183.7406 | 0.6 | 0.7 | 0.4 | 12.0 | 15.8 | 3.8 |
| Peptaibol_1452 | 1452.8756 | 0.2 | 0.0 | 0.1 | 14.1 | 12.7 | 1.8 |

**Movie S1** (separate file). Preliminary *Trichoderma* extract bioassay performed Aug. 2, 2019. This experiment tested crude *Trichoderma* extract (MB0895 10mg/mL in 1% DMSO) and negative controls (1% DMSO and NT) in three replicates on subcolonies of JKH000365, JKH000377, and JKH000380. Ten ants were included in each box. This experiment was video recorded for 8.5 h (as much as the SD card could record) and then sped to 500x the original speed. Side waste/food boxes were connected to the main fungus garden boxes for this experiment that contained sterile cornmeal in a plastic weigh boat. The boxes in the video from left to right and top to bottom are: JKH365-MB0895 waste/food, JKH377-MB0895 waste/food, JKH380-MB0895 waste/food, JKH380-NT fungus garden, JKH380-NT waste/food, JKH365-MB0895 fungus garden, JKH377-MB0895 fungus garden, JKH380-MB0895 fungus garden, JKH377-NT fungus garden, JKH377-NT waste/food, JKH365-DMSO fungus garden, JKH377-DMSO fungus garden, JKH380-DMSO fungus garden, JKH360-NT fungus garden, JKH360-NT waste/food, JKH365-DMSO waste/food, JKH377-DMSO waste/food, JKH380-waste/food.

**Dataset S1** (separate file). Sequencing and analysis metadata for the environmental ITS2 community amplicon sequencing dataset. For each sample the number of reads and ASVs are tracked through each step of the analysis pipeline.

**Dataset S2** (separate file). Sequencing and analysis metadata for the time-course *Trichoderma* JKS001884 infection ITS2 community amplicon sequencing dataset. For each sample the number of reads and ASVs are tracked through each step of the analysis pipeline.
